# Distinct regulation of *Hox* genes by Polycomb Group genes in a crustacean

**DOI:** 10.1101/2022.03.27.485719

**Authors:** Dennis A Sun, Yuri Takahashi, Rebecca J Chang, Nipam H Patel

## Abstract

Non-insect crustaceans exhibit tremendous body plan diversity. The evolution of diverse patterns of *Hox* gene expression has been implicated as a primary driver of body plan evolution between crustacean groups, but the mechanisms underlying *Hox* regulatory evolution remain unknown. We identify Polycomb and Trithorax Group proteins, crucial for proper *Hox* regulation across bilaterians, in the genome of the amphipod crustacean *Parhyale hawaiensis*, and demonstrate their essential functions in crustacean *Hox* regulation and embryonic development using CRISPR-Cas9 mutagenesis. Examination of *Hox* misexpression patterns between individual *Hox* genes with respect to timing, tissue, and PcG mutant gene in mutant embryos reveals that each crustacean *Hox* gene follows its own idiosyncratic regulatory mechanism. These results suggest a distinct regulation of *Hox* genes that may have enabled crustacean body plan evolution.

**Primary Findings:** - The genome of the amphipod crustacean *Parhyale hawaiensis* contains all core Polycomb Group (PcG) and Trithorax Group (TrxG) proteins

- CRISPR-Cas9 mutagenesis of PcG proteins induces homeotic transformations and misexpression of *Hox* genes that differ from similar experiments in insects

- PcG knockout embryos show proper initiation of *Hox* expression boundaries at early developmental stages

- Each of the three posterior *Hox* genes in *Parhyale* displays distinct patterns of misexpression in response to PcG knockout

- *Hox* regulation appears to occur via different mechanisms in the nervous system vs. limbs

- PcG phenotypes reveal the potential for distinct layers of *Hox* regulation in crustaceans

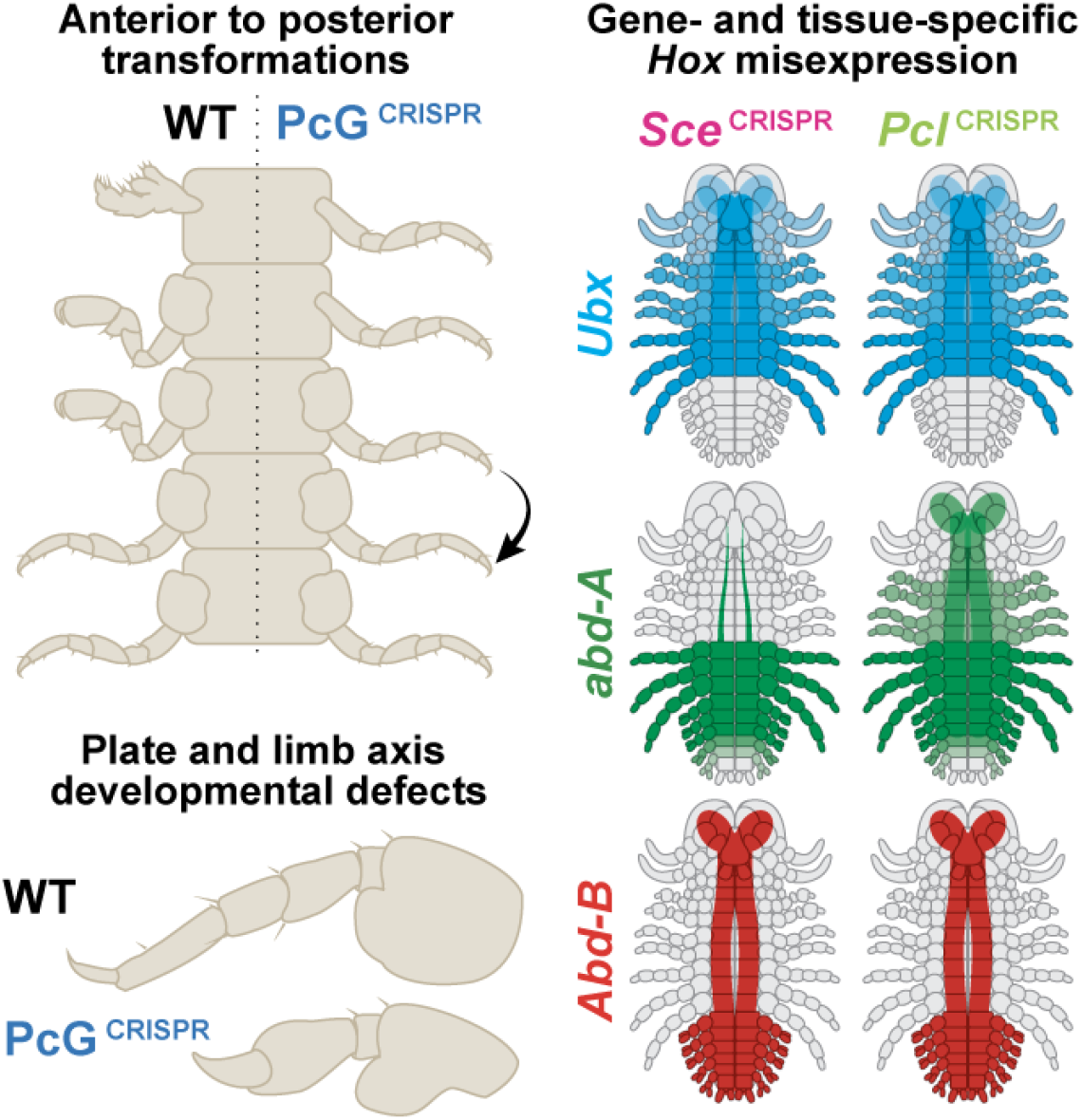

## Introduction

Non-insect crustaceans are a group of organisms with tremendous variation in body organization. Unlike hexapods or chelicerates, which stereotypically only have head and thoracic appendages, crustaceans generally have one appendage for every body segment (Angelini and Kaufman, 2005; Hughes and Kaufman, 2002). Moreover, unlike the limbs found in myriapods (millipedes and centipedes), which are largely serially identical along the body axis, crustaceans have a tremendous variety of limbs for different functions, such as feeding, grasping, walking, or swimming.

Individual groups of crustaceans have evolved different arrangements of these appendages, which are referred to as bauplans, or body plans (Averof and Akam, 1995; Deutsch and Mouchel-Vielh, 2003). Some of these body plans are familiar to most people, such as the decapod body plans of shrimp, crabs, and lobsters, or the isopod body plan of pill bugs and woodlice (Schwentner et al., 2017, 2018; Wolfe et al., 2019). Beyond those with familiar body plans, many other crustacean groups exist, including remipedes, fairy shrimp, and shield shrimp, each with its own unique arrangement of limbs along the anterior-posterior axis. As a result of these Swiss Army knife-like bodies, crustaceans are able to survive in numerous aquatic and terrestrial environments.

The evolution of crustacean body plan diversity is hypothesized to be a consequence of variations in the expression of *Homeotic (Hox)* genes, a family of genes crucial for establishing identity along the anterior-posterior axis in diverse animal groups (Averof and Akam, 1995; Averof and Patel, 1997; Hrycaj and Wellik, 2016; Krumlauf, 1994; Panganiban et al., 1995). For example, shifts in the expression domain of the *Hox* gene *Ultrabithorax* have been correlated with the emergence of novel identities across the crustacean tree (Averof and Patel, 1997). These results have suggested that crustacean *Hox* regulatory network evolution may be a driving force in crustacean body plan evolution.

Recent experiments in the amphipod crustacean *Parhyale hawaiensis* have begun to directly reveal the function of *Hox* genes in patterning crustacean body plans. Using CRISPR-Cas9 mutagenesis, RNA interference, and heat shock-induced misexpression, previous work has characterized the expression (Liubicich et al., 2009; Serano et al., 2016) and function (Liubicich et al., 2009; Martin et al., 2015; Pavlopoulos et al., 2009) of each of the *Parhyale Hox* genes (summarized in Supp. Fig. 1.1D).

For example, the *Hox* gene *Ultrabithorax (Ubx)* is expressed in most of the thoracic appendages (T2-8) in *Parhyale*: at lower levels in the claws (T2/3) and higher levels in the walking legs (T4/5) and jumping legs (T6-8). RNAi knockdown and CRISPR knockout of *Ubx* reveal that this gene is necessary for the forward walking leg (T4/5) identity, and overexpression experiments have revealed that ectopic *Ubx* expression can induce transformations of anterior body segments towards T4/5 walking leg identity (Supp Fig. 1.1F). These results indicate that *Ubx* expression is both necessary and sufficient for establishing T4/5 forward walking leg identity.

Moreover, knockout experiments examining multiple *Hox* genes in *Parhyale* have indicated that *Hox* genes also regulate one another (Jarvis et al., 2022). For example, upon CRISPR-Cas9 knockout of the *Hox* gene *Abdominal-B*, normally found in the abdominal appendages (A1-6), expression of *Ubx* expands into the previous *Abd-B* domain, suggesting that *Abd-B* represses *Ubx* (summarized in Supp. Fig. 1.1E). Knockout of both *Abd-B* and *Ubx* results in expansion of *Scr* expression towards the posterior, inducing homeotic transformations to T1 identity in the regions lacking *abd-A* expression; thus, *Ubx* and *Abd-B* repress *Scr*. These results suggest that *Hox* cross-regulatory mechanisms are essential for proper body organization in *Parhyale*.

While these studies have revealed the function of *Hox* genes in establishing regional identity and in cross-regulation, the upstream mechanisms of crustacean *Hox* regulation remain unknown. In insects, early *Hox* regulation appears to be governed at two phases: early establishment of *Hox* boundaries by upstream transcription factors, and maintenance of proper *Hox* expression through later stages of development (Akam, 1987; Denell, 1978; Irish et al., 1989; Maeda and Karch, 2006, 2015; Puro and Nygrén, 1975; Wedeen et al., 1986). For example, in *Drosophila*, the gap and pair-rule gene networks activate or repress *Hox* genes through special cis-regulatory elements known as initiator elements, which establish active or repressive chromatin in specific regions of the genome for each body segment (Maeda and Karch, 2006, 2015; Peifer et al., 1987)(Supp. Fig. 1A). These chromatin marks are laid down and maintained by the activity of Polycomb Group (PcG) and Trithorax Group (TrxG) genes, two families of general transcriptional repressors and activators, respectively (Geisler and Paro, 2015; Kassis et al., 2017; Schuettengruber et al., 2017).

PcG and TrxG genes were first identified as *Hox* regulators in *Drosophila* due to the homeotic transformations induced by perturbation of PcG/TrxG function. For example, maternal-zygotic null embryos for many PcG genes exhibit homeotic transformations of all thoracic segments towards the most posterior identity (A8)(Breen and Duncan, 1986; Martin and Adler, 1993)(Supp. Fig. 1B-C). Within these mutants, most of the *Drosophila Hox* genes (such as *Scr*, *Dfd*, *Antp*, *Ubx*, *abd-A*, and *Abd-B*) appear to be broadly misexpressed anterior to their wildtype boundary and in a variety of different embryonic tissues, including the embryonic ectoderm, nervous system, and mesoderm (Simon et al., 1992; Wedeen et al., 1986). Despite this broad misexpression of many genes, it is proposed that the cross-regulatory repression of more anterior *Hox* genes by *Abd-B* results in a uniform transformation of all body segments to the most posterior identity (Breen and Duncan, 1986; Simon et al., 1992; Struhl and White, 1985).

In contrast, some TrxG genes were observed to induce homeotic transformations in the opposite direction – posterior to anterior – as a result of decrease or loss of proper *Hox* expression (Breen and Harte, 1991, 1993)(Supp. Fig. 1B). Others TrxG genes were identified through their dominant suppressor effects on PcG mutants, indicating their competing roles in *Hox* regulation (Fauvarque et al., 2001; Kennison and Tamkun, 1988).

These two families of genes appear to have globally similar functions in vertebrate *Hox* regulation. Loss of PcG or TrxG function in mice often results in *Hox* misexpression and homeotic transformations (Akasaka et al., 2001; Brinkmeier et al., 2015; Guenther et al., 2005; Isono Kyo-ichi et al., 2005; del Mar Lorente et al., 2000; Wang et al., 2007; Yu Benjamin D. et al., 1998), suggesting that the ancestral bilaterian *Hox* cluster was regulated by PcG/TrxG function. However, mouse *Hox* regulation differs from that of *Drosophila* in that *Hox* expression is first initiated in a temporal sequence within a single tissue region (the posterior primitive streak), rather than being initiated in spatially separate regions (Deschamps and van Nes, 2005; Deschamps and Wijgerde, 1993; Kmita and Duboule, 2003; Mallo and Alonso, 2013; Wang et al., 2004). This early activation is then followed by an expansion of the expression domains towards their final expression patterns, a process not governed by cell migration, but instead by progressive changes to histone marks associated with PcG/TrxG function (Chambeyron and Bickmore, 2004; de Graaff Wim et al., 2003; Deschamps and van Nes, 2005; Deschamps and Wijgerde, 1993; Rastegar Mojgan et al., 2004; Soshnikova and Duboule, 2009). Thus, in vertebrates, the final expression pattern of *Hox* genes appears to be both established and maintained by PcG/TrxG function.

The molecular functions of PcG/TrxG complexes across organisms are diverse, but primarily occur via the chromatin modifications they establish. Multiple PcG complexes exist, but are broadly broken down into three groups: Polycomb Repressive Complex 1 (PRC1), Polycomb Repressive Complex 2 (PRC2), and Pleiohomeotic Repressive Complex (PhoRC) (Fig. 1A, adapted from Erokhin et al., 2018; Schuettengruber et al., 2017). Within these broader groups, additional sub-complexes have been characterized, such as PRC1 vs. dRAF, and PRC2.1 vs. PRC2.2, each of which has its own particular biochemical function. TrxG complexes are more diverse both in function and number, including BAP/PBAP, TAC1, COMPASS and COMPASS-like, and other complexes (Fig. 1A, adapted from Kassis et al., 2017). Many of the TrxG complexes directly reverse the biochemical function of PcG complexes; as such, the two groups of genes are often considered to function in opposition (Geisler and Paro, 2015).

**Figure 1:**
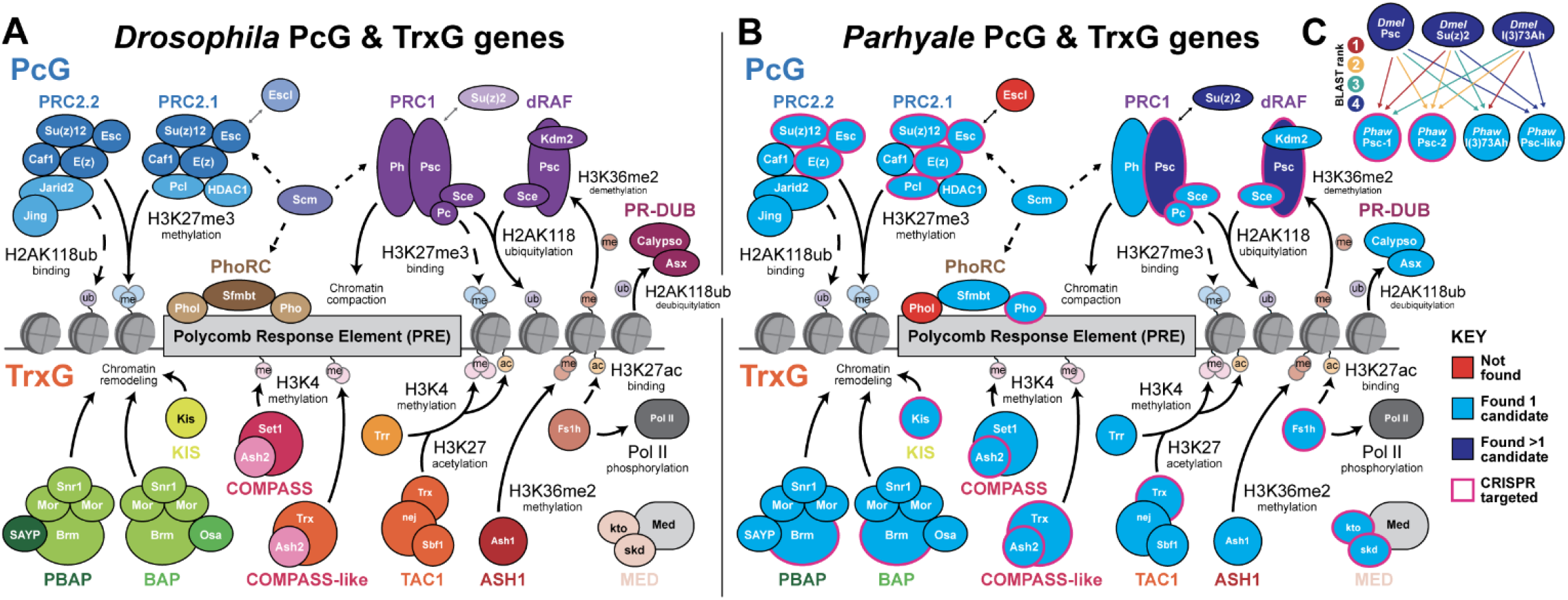
The Parhyale hawaiensis genome contains all core PcG and TrxG genes. A) Schematic of the core PcG and TrxG complexes found in *Drosophila*, in addition to other genes classified as PcG/TrxG genes, alongside known gene regulatory functions. Adapted from (Erokhin et al., 2018; Geisler and Paro, 2015; Kassis et al., 2017; Schuettengruber et al., 2017). B) Summary of PcG/TrxG genes found in *Parhyale* using reciprocal BLAST. All genes except *esc-l* and *pho-l* were found in the *Parhyale* genome. C) *Drosophila Psc* and *Su(z)2* in *Parhyale* map ambiguously to two transcripts, which we have named *Psc-1* and *Psc-2*. A third potential *Psc* paralog was also identified. We named this gene *Psc-like*.

In this paper, we reveal the role of PcG and TrxG genes in the regulation of *Hox* genes in the amphipod crustacean *Parhyale hawaiensis*. We demonstrate that all core members of these complexes are present in the *Parhyale* genome and expressed during embryonic development. Using CRISPR-Cas9 mutagenesis, we ablate PcG and TrxG function and identify homeotic transformations induced by PcG knockout, indicating that these genes are essential for *Parhyale Hox* regulation. In addition, we examine the effects of PcG knockout on the expression of *Parhyale Hox* genes using *in situ* hybridization chain reaction and immunohistochemistry. These data reveal that the *Parhyale Hox* genes appear to have idiosyncratic regulatory mechanisms with respect to developmental timing, tissue type, and specific PcG complex components. Our results provide the first mechanistic insights into the upstream regulatory architecture of the *Parhyale Hox* complex, and suggest that *Hox* genes in crustaceans may have evolved distinct regulation. Such distinct regulation offers a mechanistic explanation for how different crustacean groups were able to evolve distinct patterns of *Hox* expression to generate their dramatic body plan diversity.

## Results

### The *Parhyale* genome contains all core PcG/TrxG components

To identify candidate genes of interest for CRISPR-Cas9 mutagenesis, we examined the current literature on *Drosophila melanogaster* and *Mus musculus* PcG/TrxG function. We cataloged phenotypes of interest for a number of genes (Supp. Tables 1 and 2), and used reciprocal BLAST to search for all core PcG/TrxG complex components in several *Parhyale hawaiensis* transcriptomes and genome annotations (see Methods). The data sources for each transcript, along with the gene ID in that transcript source, are listed in Supp. Tables 3 and 4.

We were able to identify all of the core PcG/TrxG genes we examined in the *Parhyale* genome. We observed that *Parhyale* has a single copy of *extra sex combs* and *pleiohomeotic*, in contrast to *Drosophila*, where additional gene duplications with redundant functionality (*extra sex combs-like* (*esc-l*) and *pleiohomeotic-like* (*pho-l*) appear to have evolved (Figure 1B)(Brown et al., 2003; Kurzhals et al., 2008; Ohno et al., 2008). In addition, we found four *Parhyale* genes that appeared to be related to the *Drosophila* genes *Psc*, *Su(z)2*, and *l(3)73Ah*.

*Psc* and *Su(z)2* are two genes in *Drosophila* that are located adjacent to each other in the genome and which have semi-redundant functionality (Brunk et al., 1991; Lo Stanley M. et al., 2009). In *Parhyale*, we found four potential candidates for *Psc* and *Su(z)2* (Fig. 1C, Supp. Fig. 2.1F, G). One of these candidates, which we named *Phaw Psc-1*, appeared to be the top BLAST hit for both *Dmel Psc* and *Dmel Su(z)2*. A second candidate, which we named *Phaw Psc-2*, appeared to be the second-best BLAST hit for both *Dmel Psc* and *Dmel Su(z)2*. For both *Dmel Psc* and *Su(z)2*, we also found a third BLAST hit, which we identified as *Phaw l3(73)Ah* based on BLAST to *Dmel l3(73)Ah*. Finally, the fourth-best BLAST hit to both *Dmel Psc* and *Dmel Su(z)2* appeared to be another protein with a RING-HC PCGF and RAWUL PCGF2-like domain, similar to *Psc* (Supp. Fig. 2.1F, G). We named this gene *Phaw Psc-like*.

To determine whether each of these PcG/TrxG genes were expressed during embryonic development in *Parhyale*, we leveraged the recently described Mikado transcriptome, which also serves as an improved genome annotation (Sun et al., 2021). We identified the best Mikado transcript for each PcG/TrxG gene, and examined the expression of that transcript at four developmental stages (S13, S19, S21, S23; three replicates per stage) from a previously described transcriptome (Supp. Fig. 1.2)(data from Sun et al. 2021). Based on these data, all PcG/TrxG genes we examined appeared to be expressed at these four stages of development.

Based on the potential to induce homeotic phenotypes previously described in *Drosophila*, we selected a number of PcG and TrxG genes to perform CRISPR-Cas9 mutagenesis. For PcG, we designed CRISPR guide RNAs to the PRC1 genes *Sce*, *Pc*, *Psc-1*, *Psc-2*; the PRC2 genes *Su(z)12*, *E(z)*, *Esc*, and *Pcl*; and the PhoRC gene *pho*. For TrxG, we designed CRISPR guide RNAs to the BAP/PBAP gene *Brm*; *Kis*; the COMPASS and COMPASS-like genes *Ash2* and *Trx*; and three additional genes categorized as TrxG members: *Fs1h*, *kto*, and *skd*. For each gene, we designed guide RNAs targeting the 5’-most region of the RNA sequence for which BLAST evidence suggested homology to either *Drosophila* or another arthropod. We injected pairs of guide RNAs into 1- or 2-cell embryos and screened embryos for phenotypes using confocal and widefield microscopy.

### PcG KO induces homeotic transformations of anterior segments to T4/T5 walking leg identity

When we examined knockouts of PcG genes, we consistently observed striking homeotic transformations of anterior body segments towards T4/5 walking leg identity. Representative hatchlings from select CRISPR experiments are shown in Fig. 2, illustrating the homeotic transformations we observed. We observed this phenotype in hatchlings for 7/9 of the genes we targeted using CRISPR (*Sce*, *Pc*, *Su(z)12*, *E(z)*, *Esc, Pcl*, and *pho*, see Table 1). We most frequently observed transformations of T2/3 claw appendages to T4/5 walking leg identity, but we also observed for some genes, such as *Pcl* and *Su(z)12*, transformations of antennae towards T4/5 walking leg identity. We also observed another class of apparent transformations of A1-3 swimming leg identities to A4-6 anchoring leg identities in *Pc* mutants, but at frequencies much lower than transformations to T4/5 walking leg identities. Finally, among all mutants exhibiting transformations towards T4/5 identity, we also observed defects in mouthpart morphology, where mouthparts appeared to exhibit “stubby” morphology not obviously analogous to any known wild type morphology. A summary of the phenotypes we observed can be found in Table 1.

**Fig. 2.**
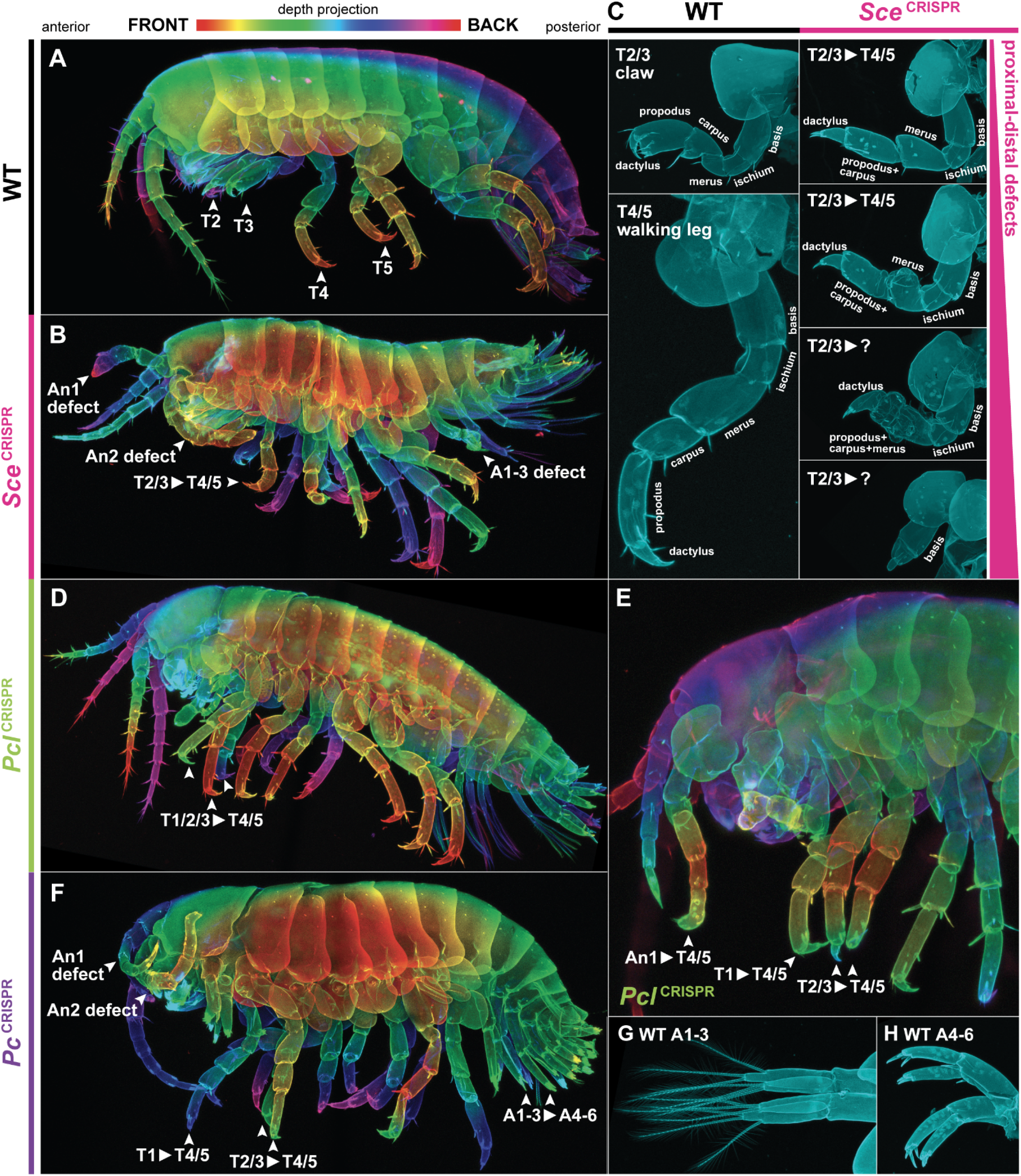
PcG knockout hatchlings exhibit homeotic transformations. A) Representative wildtype *Parhyale* hatchling. Of note, the T2 and T3 appendages are claws (chelipeds) and the T4 and T5 appendages are forward walking legs (forward pereopods). B) Representative *Sce* mutant hatchling. *Sce* mutants show various limb defects, in addition to homeotic transformation of anterior thoracic appendages (T1-3) toward T4/5 identity. C) Proximal-distal phenotypes concurrent with homeotic phenotypes in *Sce* mutants. Left column shows dissected WT T2/3 and T4/5 appendages. Right column shows a range of different proximal-distal defects in *Sce* T2/3 appendages. Despite these proximal-distal defects, comparison of the merus of *Sce* mutant T2/3 appendages to the merus of WT T2/3 and WT T4/5 shows a clear homeosis. D) Representative *Pcl* mutant hatchling, showing prominent T1/2/3 -> T4/5 homeotic transformations. *Pcl* mutants did not often appear to have proximal-distal defects. E) Representative *Pcl* mutant hatchling showing T1/2/3 -> T4/5 homeotic transformations, as well as An2 -> T4/5 transformation. Notably, the formation of an ectopic coxal plate suggests T4/5 transformation and phenocopies previous *Ubx* heat shock overexpression experiments (Pavlopoulos et al., 2009). F) Representative *Pc* mutant hatchling showing T1/2/3 -> T4/5 homeosis, as well as A1-3 -> A4-6 homeosis and antennal defects. G) Dissected WT A1-3 swimming leg (pleopod). H) Dissected WT A4-6 anchoring leg (uropod).

**Table 1:** Summary of PcG and TrxG CRISPR phenotypes

| Gene | Family | Complex | Homeosis | Plate Defects | Limb Defects | Other Phenotypes |
| --- | --- | --- | --- | --- | --- | --- |
| *Polycomb (Pc)* | PcG | PRC1 | T1 -> T4/5<br>T2/3 -> T4/5<br>A1-3 -> A4-6 | Missing T2 coxal and tergal plates, unfused plates, asymmetric plate fusion | Deformed antenna, deformed mouthparts, deformed limbs, proximal-distal fused limbs | – |
| *Sex combs extra (Sce)* | PcG | PRC1, dRAF | T1 -> T4/5<br>T2/3 -> T4/5 | Missing T2 coxal and tergal plates, reduced basal plates, unfused tergal plates | Deformed antenna, deformed mouthparts, deformed limbs, proximal-distal fused limbs | – |
| *Posterior sex combs-1 (Psc-1)* | PcG | ? | – | – | – | – |
| *Posterior sex combs-2 (Psc-2)* | PcG | PRC1, dRAF | Not observable (no limbs) | Loss of plates | Loss or major reduction of limbs | High embryonic lethality |
| *Suppressor of zeste 12 (Su(z)12)* | PcG | PRC2 | An -> T4/5<br>Mp -> T4/5<br>T1 -> T4/5<br>T2/3 -> T4/5 | Missing T2 coxal and tergal plates, reduced basal plates, unfused tergal plates | Deformed antenna, deformed mouthparts, deformed limbs, proximal-distal fused limbs, twisted limbs | – |
| *Enhancer of zeste (E(z))* | PcG | PRC2 | T2/3 -> T4/5 | Reduced basal and coxal plates | Deformed antenna, deformed mouthparts, deformed limbs, proximal-distal fused limbs, twisted limbs | Major malformations, possible body segment fusion |
| *Extra sex combs (Esc)* | PcG | PRC2 | T2/3 -> T4/5 | Reduced basal and coxal plates | Deformed antenna, deformed mouthparts, deformed limbs, proximal-distal fused limbs, twisted limbs | Major malformations |
| *Polycomb-like (Pcl)* | PcG | PRC2.1 | An -> T4/5<br>Mp -> T4/5<br>T1 -> T4/5<br>T2/3 -> T4/5 | Missing T2 coxal and tergal plates, reduced basal and coxal plates, unfused tergal plates | Deformed antenna, deformed mouthparts, deformed limbs, proximal-distal fused limbs, twisted limbs | – |
| *Pleiohomeotic (pho)* | PcG | PhoRC | T1 -> T4/5<br>T2/3 -> T4/5<br>Coxal plate<br>T2-5 -> posterior plates | Reduced basal and coxal plates | Deformed antenna, deformed mouthparts, deformed limbs, proximal-distal fused limbs, twisted limbs | – |
| *Trithorax (Trx)* | TrxG | TAC1 | – | – | – | – |
| *Brahma (Brm)* | TrxG | PBAP, BAP | – | Missing plates | Missing limbs | Embryonic lethality, major malformations |
| *Kismet (Kis)* | TrxG | – | – | – | – | – |
| *Absent, small, or homeotic discs 2 (Ash2)* | TrxG | COMPASS, COMPASS-like | – | Reduced coxal, tergal, basal plates; unfused tergal plates | Rounded limbs | Major malformations |
| *Female sterile (1) homeotic (Fs1h)* | TrxG | – | – | – | – | Embryonic lethality |
| *Kohtalo (kto) aka Med12* | TrxG (?) | Mediator | – | Unfused tergal plates | Deformed limbs | Embryonic lethality, major malformations |
| *Skuld (skd) aka Med13* | TrxG (?) | Mediator | – | Unfused tergal plates | Deformed limbs | Embryonic lethality, major malformations |
| *CTCF* | Other | – | – | – | – | – |

The transformations of anterior segments to T4/5 identity are notable in that they appear identical to the homeotic transformations observed by heat shock misexpression of *Parhyale Ultrabithorax* (Pavlopoulos et al., 2009). In that previous study, anterior segments were frequently transformed towards T4/5 walking leg identities in heat-treated embryos and hatchlings. Based on our understanding of the role of each *Hox* gene in patterning the *Parhyale* body plan, we hypothesized that the homeotic phenotypes we observed in our PcG knockouts would be a result of misexpression of *Ubx*. These results suggest that PcG genes are necessary for proper *Hox* regulation in *Parhyale*.

### PcG genes are essential for proper limb proximal-distal axis development and plate formation

In addition to the homeotic phenotypes we observed, we also observed other types of developmental defects in CRISPR-targeted embryos. In particular, for 8/9 genes we examined (excluding *Psc-1*), we observed various non-homeotic defects in limb morphogenesis (see Table 1). For example, in addition to homeosis of T2/3 -> T4/5 in *Sce* mutant embryos, we also observed fusions or loss of leg segments (Fig. 2C, Fig. 3B, B’). This proximal-distal defect phenotype varied from hatchling to hatchling, and from limb to limb across the body plan. To assess the potential for off-target effects of our knockouts, we performed thorough phenotype analyses of hatchlings from CRISPR knockouts using single guides of three genes:*Sce*, *Pcl*, *Su(z)12*. For all three genes, we were able to recover all of our cataloged phenotypes for each guide RNA (Supp. Figs. 2.2, 2.3, 2.4).

**Fig. 3:**
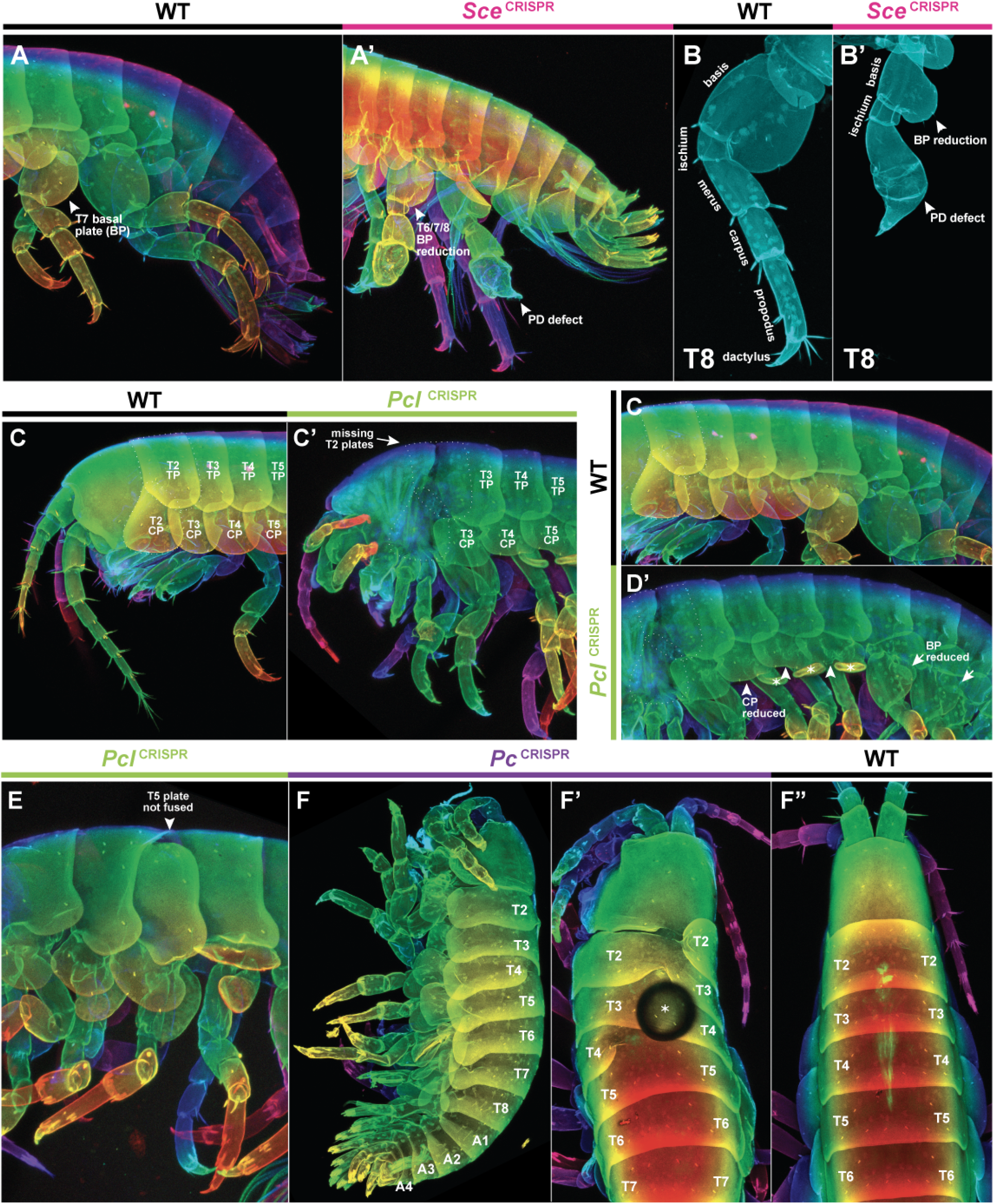
Proximal-distal and plate defects of PcG mutants. A) Wildtype *Parhyale* posterior showing T6-8 jumping legs and abdomen. A’) *Sce* mutant hatchling showing proximal-distal defects in T6/8 appendages and deformed basal plates. B) WT T8 appendage. B’) *Sce* mutant T8 appendage showing proximal-distal leg segment fusion and reduced basal plate. C) WT anterior, with tergal and coxal plates labeled. C’) *Pcl* mutant showing loss, with dotted line indicating position of missing T2 tergal and coxal plates, as evidenced by the lack of the triangular T2 coxal plate. D) View of tergal and coxal plates along WT thorax. D’) *Pcl* mutant showing reduced coxal and basal plates. Marked with asterisks are the gills, normally not visible from a lateral view in a WT hatchling, but revealed by coxal plate reduction. E) *Pcl* mutant showing unfused tergal plate (labeled). This sample also shows reduced coxal plates and visible gills. F-F ’) Lateral and dorsal views of the same *Pc* mutant hatchling, illustrating asymmetrical fusion of plates between left and right halves of the body. Asterisk marks a bubble in the mounting medium. F’’) Dorsal view of a WT hatchling.

In addition, we observed seemingly improper cuticle deposition, resulting in wrinkled-looking appendages, bulbous limbs, twisted limbs, and other general morphological defects (Fig. 3, Supp. Figs. 2.1, 2.2, 2.3, 2.4, 2.5). These results indicate that PcG genes are necessary for proper development of limbs along the proximal-distal axis, potentially interfacing with the leg gap gene system described in (Bruce and Patel, 2020; Clark-Hachtel and Tomoyasu, 2020), and that loss of PcG function can induce general morphogenic irregularities.

We also observed defects in the development of tergal, coxal, and basal plates in a number of PcG knockout hatchlings. For example, for all genes other than *Psc-1* and *Psc-2*, we observed reduced coxal, tergal, and basal plates (Fig. 3B, B’, D, D’). These phenotypes were reminiscent of previous phenotypes induced by knockout of wing developmental genes such as *vestigial* and *scalloped* (Clark-Hachtel and Tomoyasu, 2020). For many PcG genes, we also observed total loss of plate morphogenesis, such as loss of the entire T2 tergal and coxal plates, which usually present with a stereotypical triangular shape (Fig. 3C, C’). In other cases, we observed “floating” tergal plates, where the left and right halves of the hatchling failed to properly close on the dorsal side of the embryo (Fig. 3E). Some of these “floating” plates appeared to be caused by improper pairing of the left and right halves of the embryo (Fig. 3F, F’, F’’). For example, Figure 3F’ shows a hatchling where the left T2 plate has joined with the right T3 plate. These results suggest that PcG genes may also play a role in regulating the process of dorsal closure.

The phenotypes we observed appear to affect body segments in more restricted domains than those governed by *Hox* regionalization alone. For example, the T2 segment, in which we frequently observed total loss of the tergal and coxal plates, normally develops into a clawed appendage. The T3 segment also develops a clawed appendage, but we did not observe total loss of T3 tergal and coxal plates in our mutants. These results reveal potential segmental identities unique to individual numerical body segments, regulated by a more specific mechanism than that of *Hox* regional identity, in which PcG genes play some important role.

### *Parhyale Psc-2*, but not *Psc-1*, is essential for proper embryonic development

*Parhyale Psc-1* and *Psc-2* were the only ambiguously identified orthologs of PcG or TrxG genes found in our examination of the *Parhyale* genome. CRISPR-Cas9 mutagenesis of *Psc-2* caused major developmental defects, including apparent fusion of body segments and dramatic truncation of limbs (Supp. Fig. 2.1). Most *Psc-2* hatchlings did not hatch on their own, and required dissection from the egg at late developmental stages. CRISPR treatment with guides targeting *Psc-1* produced no observable phenotypes. Thus, *Psc-2* is indispensable for embryonic development, and likely serves a role in PcG complex function, while *Psc-1* does not appear to be essential for development.

### TrxG KO induces embryonic lethality, major developmental defects, or no phenotype; but not homeosis

In contrast to the consistent phenotypes observed for PcG knockouts, TrxG knockout hatchlings displayed a wider variety of phenotypes including embryonic lethality, major developmental defects, or no visible phenotype. We also did not observe any obvious homeotic transformations in TrxG knockout hatchlings.

Knockouts of *Brm* and *Fs1h* in *Parhyale* appeared to result in embryonic lethality (Supp. Fig. 2.6D-D’, E). None of the *Fs1h* knockout embryos survived long enough to hatch, and only a small number of *Brm* embryos survived to late developmental stages. In the few *Brm* hatchlings we recovered, it appeared that approximately half of the embryo was deleted (Supp. Fig. 2.6D). This phenotype is consistent with the loss of one-half of the embryo starting at the 2-cell stage, based on previous lineage tracing experiments (Gerberding et al., 2002; Price et al., 2010), and suggests that *Brm* is essential for cell viability during *Parhyale* development.

Knockouts of *kto* and *skd*, members of the Mediator complex, showed major morphological defects such as compaction along the anterior-posterior axis, including fusion between adjacent body segments and defects in dorsal closure (Supp. Fig. 2.6A-A’’, B-B’’). Careful analysis of *kto* and *skd* mutant limbs did not reveal any obvious homeotic transformations. *Ash2* knockout hatchlings showed occasional defects in plate development, with reduced plates reminiscent of PcG mutants, as well as twisted limbs, but did not appear to show any obvious homeotic transformations (Supp. Fig. 2.6C-C’’).

*Trx* and *Kis* knockout embryos did not show any obvious phenotypic differences from WT embryos with the first pair of guide RNAs we used for each gene (Supp. Fig. 2.7). We designed additional pairs of guide RNAs targeted towards the 5’ end of conserved domains of these genes that have previously been characterized as being important for gene function. In the case of *Trx*, careful analysis of gene models from several different transcriptomes revealed that the *Trx* gene may contain two separate, non-contiguous ORFs. The 5’ ORF appeared to contain the conserved PHD domain and FYRN domains found in the 5’ end of *Drosophila Trx*, while the 3’ ORF appeared to contain FYRC and SET domains found in the 3’ end of *Drosophila Trx* (Supp. Fig. 2.7C). When we identified *Trx* in the Mikado transcriptome, the 5’ and 3’ ORFs were annotated as two separate genes, which we referred to as *Trx (DBD)* and *Trx (SET)*. Quantification of the expression of each of these two transcripts across a developmental RNA-Seq timecourse (from Sun et al., 2021) revealed that the two ORFs appeared to have differential expression over time (Supp. Fig. 2.7B). To account for the possibility of the *Trx (SET)* ORF performing a function separate from the *Trx (DBD)* ORF, we also designed a pair of guides specifically targeting the *Trx (SET)* ORF. In total, we designed 8 guides targeting *Trx* and 4 guides targeting *Kis*. We did not observe any obvious homeotic phenotypes for any of the guides we tested for these genes. Hatchlings overall showed WT morphology or mild morphological defects (such as twisted body axis) which have previously been observed in sham injection experiments (Schmid, 2011).

Our experiments were unable to reveal any obvious role of TrxG in *Hox* regulation in *Parhyale*. It is possible that the homeotic effects of TrxG knockout are more subtle than the limb transformations observed in PcG knockouts; deeper analysis will be necessary to define the morphological landmarks of such phenotypes, such as the exact pattern of setae (sensory hairs) and other fine structures which may be specified via *Hox* mechanisms (such as was performed in Almazán et al., 2021). Additional experiments, such as using RNAi to induce gene knockdown for essential genes, or CRISPR mutagenesis of multiple genes simultaneously, could also help more clearly reveal whether TrxG genes play any role in *Hox* regulation in this organism.

### PcG gene knockout induces *Hox* misexpression during limb developmental stages

Based on the homeotic phenotypes we observed, we hypothesized that *Ubx* would be misexpressed in anterior segments in PcG mutants. To test this hypothesis, we performed *in situ* hybridization chain reaction (Bruce et al., 2021; Choi et al., 2018) on CRISPR-treated embryos for *Parhyale Ubx*, along with other *Hox* genes, including *abd-A* and *Abd-B*. We performed HCR on dissected embryos treated with CRISPR guides targeting *Sce* and *Pcl* at developmental stage 23, during which limb morphogenesis is underway. For both genes, we observed strong misexpression of *Ubx* in anterior body segments (Fig. 4A, B, C; *Ubx* panel), as predicted by the observed homeotic transformations. Thus, PcG knockout induces ectopic expression of *Ubx*, resulting in homeotic transformations of anterior body segments to T4/T5 identity.

**Fig. 4.**
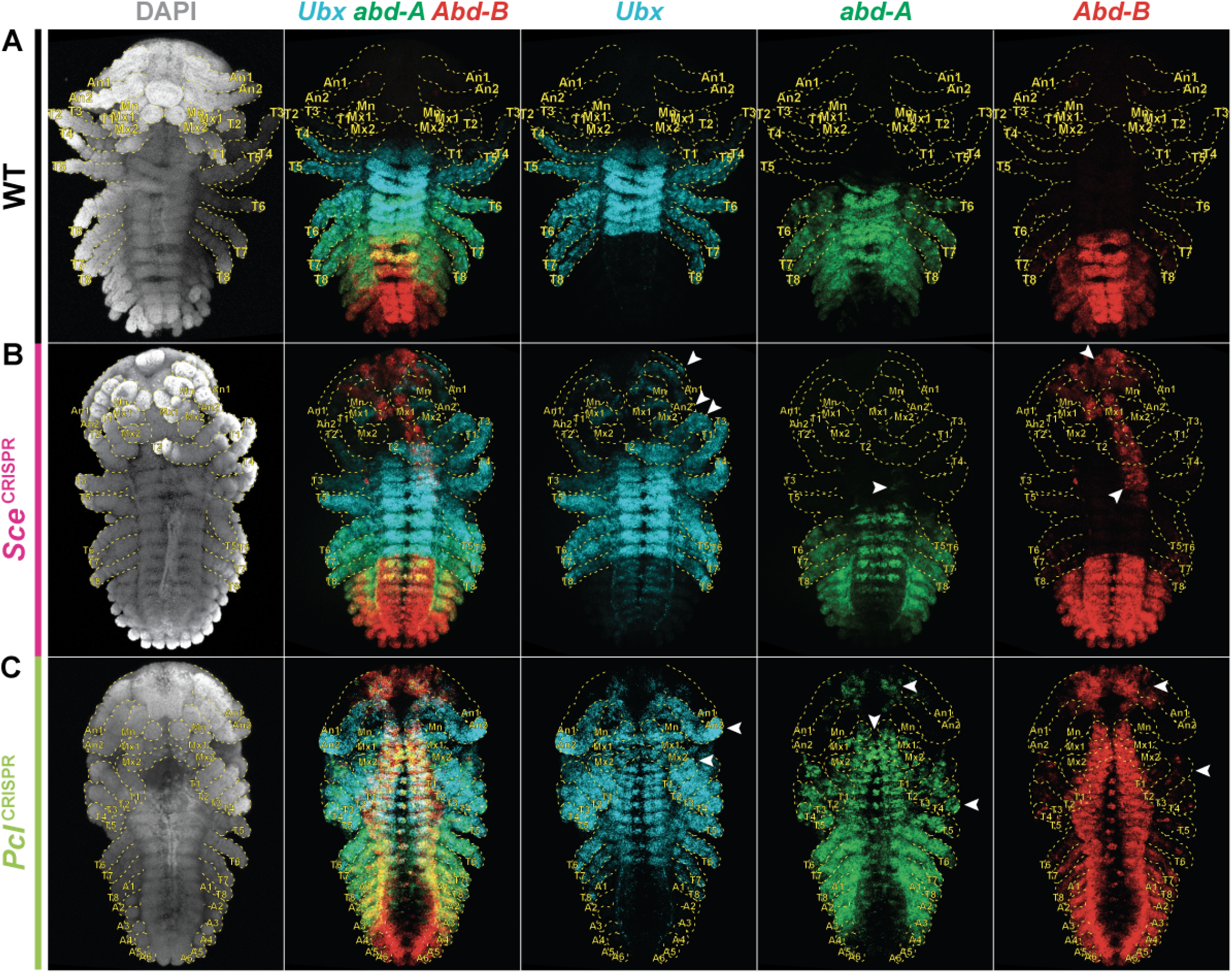
CRISPR mutagenesis of PcG induces *Hox* misexpression. A) *In situ* HCR of *Ubx, abd-A,* and *Abd-B* in a representative WT embryo at stage 23. The anterior boundary of *Ubx* expression is in the T2 appendage, with weak *Ubx* expression in T2-3 and stronger *Ubx* expression in T4-8. The anterior boundary of *abd-A* expression is the T6 appendage. Strong *abd-A* expression is found in T6-A3, with weaker, gradated expression found in A4 and A5. The anterior boundary of *Abd-B* expression is A1, and *Abd-B* expression is found throughout the abdomen (A1-6). B) *In situ* HCR of *Ubx, abd-A,* and *Abd-B* in a representative *Sce* mutant embryo at stage 23. The embryo appears to have mosaic loss of function, with the head and anterior-left half of the embryo showing *Ubx* misexpression beyond the normal WT anterior boundary (the embryo is mounted ventral side up; in the image, the left half of the embryo is on the right side). *abd-A* misexpression is slight, restricted to a few cells of the nervous system. *Abd-B* is misexpressed strongly in the nervous system and not in the limbs. C) *In situ* HCR of *Ubx, abd-A,* and *Abd-B* in a representative *Pcl* mutant embryo at stage 23. The embryo appears to have bilateral loss of function. *Ubx* misexpression expands throughout the anterior of the embryo. *abdA* misexpression appears strongly throughout the nervous system and in strong patches in the limbs, in stark contrast to weak misexpression in *Sce* mutant embryos. *Abd-B* appears strongly misexpressed in the nervous system, including long filaments of *Abd-B* misexpression in the limbs, which resembles the pattern of limb enervation.

We also examined the expression of *abd-A* and *Abd-B* in mutant embryos (Fig. 4A, B; *abd-A* and *Abd-B* panels). At late developmental stages, the tissue location and degree of misexpression of *abd-A* and *Abd-B* differed when compared to *Ubx*. For example, in *Sce* mutant embryos, whereas *Ubx* misexpression appear to expand anteriorly broadly across all tissues, *Abd-B* expression expanded anteriorly, but appeared to be restricted to the medial region of the body, where the neurogenic ectoderm is located. *abd-A* in *Sce* mutant embryos appeared to be only misexpressed in the nervous system in a few cells per body segment. These results suggest that *Sce* is necessary for repression of *Abd-B* in the nervous system, but not in the limbs, and that *Sce* weakly represses *abd-A*.

We also examined the expression of two other *Hox* genes, *Scr* and *Antp*, in mutant embryos at S23 (Fig. 5A, B). *Scr* in WT embryos is expressed in the Mx1, Mx2, and T1 segments. In mutant embryos, *Scr* appeared to be expressed within the same regions, but with a notable reduction in expression in the T1 appendage. In the Mn segment, we observed patches of ectopic *Scr* expression. *Scr* expression also appeared to exhibit trace misexpression in further anterior structures, with a few transcriptional spots in the brain and antennae (Fig. 5B, *Scr* channel, arrows). When we compared the domain of *Scr* expression to *Ubx* expression, we observed that *Ubx* expression appeared to expand into the T1 appendage, suggesting a mechanism for the reduction of *Scr* expression in that limb. In a previous study (Pavlopoulos et al., 2009), heat shock overexpression of *Ubx* was associated with decreased or nearly abolished *Scr* expression within its normal domain, providing an explanation for the reduction in *Scr* expression in T1 as *Ubx* expression expands into that segment, as well as a lack of strong expression in more anterior segments, where *Ubx* is strongly misexpressed.

**Fig. 5.**
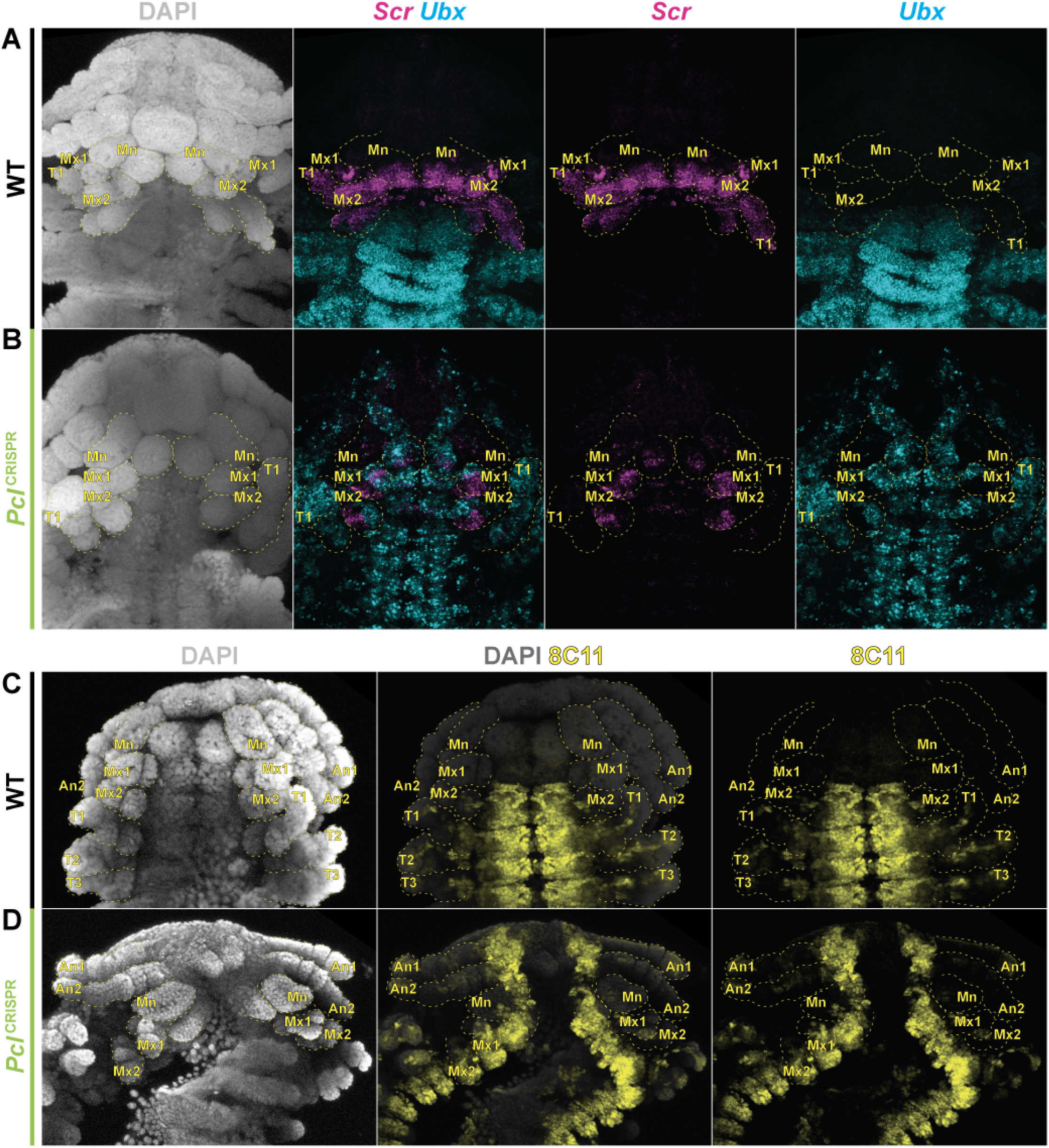
Misexpression of *Scr* and *Antp* in PcG mutants. A) *In situ* HCR of *Scr* and *Ubx* in the head of a WT embryo at stage 23. *Scr* is expressed in Mx1, Mx2, and T1. *Ubx* is expressed in T2 and T3. B) *In situ* HCR of *Scr* and *Ubx* in the head of a *Pcl* mutant embryo at stage 23. *Scr* expression expands anteriorly into Mn, as well as a few transcriptional spots (arrowheads) in the antennal segments. *Scr* expression is largely abolished from T1, and is retained in patches of Mx1 and Mx2. *Ubx* expression expands anteriorly into all anterior segments. *Ubx* and *Scr* appear to be expressed in mutually-exclusive domains in Mx1 and Mx2. C) Antibody stain using ms anti-Antp (8C11) in WT embryo at stage 21. The anterior boundary of Antp protein is the Mx2 segment. D) Antibody stain using 8C11 in a *Pcl* mutant embryo at stage 21. Antp protein is found more anterior than the normal boundary, expanding into the nervous system of the An1, An2, Mn, and Mx1 segments.

However, in PcG knockout embryos, *Scr* expression is retained in Mx1 and Mx2 (Fig. 5B). Closer examination of the patterns of *Scr* and *Ubx* expression within the same embryo revealed that *Scr* expression in PcG mutants is maintained in the posterior half of the Mx1 and Mx2, and missing in the anterior half of the limbs. In these appendages, *Ubx* is expressed in a nearly mutually-exclusive domain when compared to *Scr*. This phenotype can be explained if *Scr* loss in the anterior half of the Mx1 and Mx2 limbs is also mediated by *Ubx* repression of *Scr*.

In addition, these results suggest that the posterior half of the Mx1 and Mx2 appendages may lack some activating factor necessary for PcG derepression-mediated *Ubx* misexpression. These findings of mosaic *Hox* expression in mouthparts may also explain the relatively infrequent occurrence of mouthpart -> T4/5 transformations we observed in our knockout experiments, as well as the stubby or malformed morphology of mouthparts in most PcG mutants, which did not clearly resemble any known limb type.

To examine *Antp* expression, we used the cross-reactive mouse anti-Antp antibody (8C11) rather than *in situ* HCR. A challenge of *in situ* hybridization for detecting *Antp* in crustaceans is that crustaceans express a hybrid *Antp-Ubx* transcript in limbs across the broad region overlapping both *Antp* and *Ubx* expression, in addition to broad expression of *Antp* in the nervous system (Serano et al., 2016; Shiga et al., 2006). Antibody staining has previously revealed that functional Antp protein is only produced in the T2/3 appendages and in the nervous system – thus, examination of *Antp* expression requires direct detection of Antp protein. Using 8C11, we observed that the anterior boundary of *Antp* expression was perturbed in mutant embryos (Fig. 5C, D). In WT embryos, Antp protein in the nervous system is bounded in the anterior at the Mx2 segment. However, we observed strong Antp expression in the nervous system throughout the entire body axis in *Sce* and *Pcl* mutant embryos.

Together, these results indicate that PcG genes are essential for proper repression of *Hox* expression in *Parhyale*, as loss of PcG expression induces ectopic *Hox* expression for all of the five *Hox* genes we examined. In addition, these results reveal that each *Hox* gene in *Parhyale* appears to have a differential requirement for PcG function in different tissues and regions of the body.

### PcG knockout-induced misexpression begins after wild-type *Ubx* boundary establishment

To examine the role of PcG genes in the establishment of *Hox* expression patterns, we performed HCR on a developmental time course of *Sce* and *Pcl* mutant embryos at developmental stages S13 (early germband), S19 (posterior flexure/ late germband), S21 (early limb morphogenesis) and S23 (late limb morphogenesis) and compared the expression of *Ubx* and *Abd-B* to that of wild-type embryos.

We observed that the anterior boundary of *Ubx* expression appeared unchanged in CRISPR-treated versus WT embryos (Fig. 6A, Supp. Fig. 6.1A, B). In WT embryos, *Ubx* expression begins at the very earliest stages in parasegment 5, and then expands anteriorly into parasegment 4 by around S13 (Liubicich et al., 2009). At S13, for both *Sce* and *Pcl* knockout embryos, we observed *Ubx* expression with an anterior boundary of parasegment 4 (5/5 *Sce* embryos, 9/9 *Pcl* embryos; Fig. 6A, Supp Fig. 6.1B). In the case of *Sce* we observed several embryos where one side of the embryo appeared developmentally delayed relative to the other, suggesting that one half of the embryo had been affected by CRISPR-Cas9 mutagenesis, while the other half had not (Fig. 6A). In these embryos, we also observed that the anterior boundary of *Ubx* expression was unchanged.

**Fig. 6:**
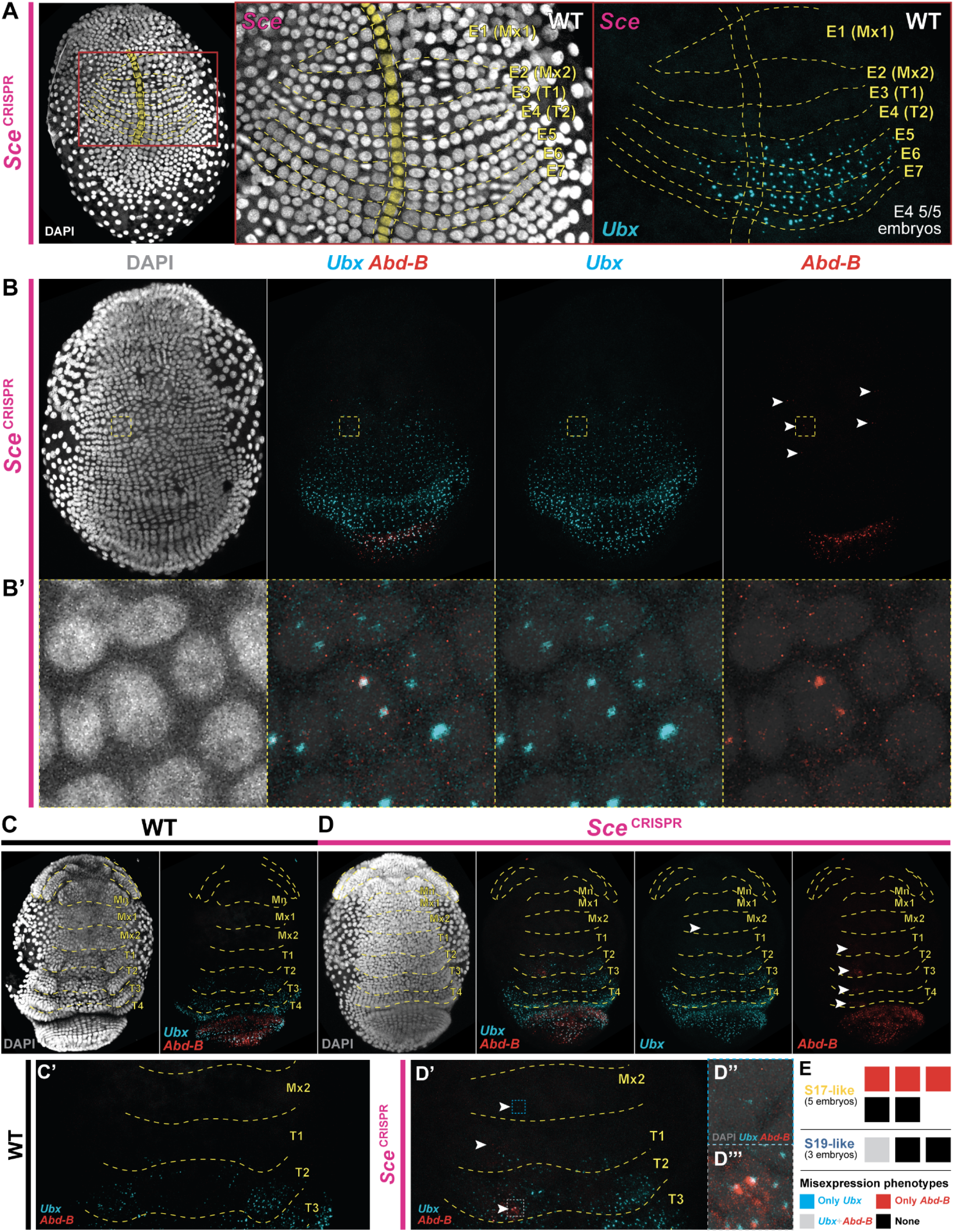
Differential timing of PcG knockout-mediated misexpression for *Ubx* and *Abd-B*. A) *In situ* HCR of *Ubx* in a unilateral *Sce* mutant embryo. Based on a delayed pattern of cell divisions found in the right half of the embryo (embryo mounted ventrally; right half of embryo is on the left of the image), we infer that this half has lost *Sce* expression. The anterior boundary of *Ubx* expression between the *Sce* mutant and wildtype halves is the same, at parasegment E4, which primarily contributes to the T2 appendage. We observed a parasegment E4 anterior boundary of *Ubx* expression in 5/5 embryos examined at this stage. B) *In situ* HCR of *Ubx* and *Abd-B* in an S17-like embryo. The very early stages of *Abd-B* expression have begun in the posterior of the embryo. Concurrently, *Abd-B* misexpression is found in the neurogenic ectoderm of anterior segments. B’) Inset region of interest showing clear *Ubx* and *Abd-B* transcription spots in the nucleus of an *Sce* S17-like mutant embryo. Previous work has shown that the *Parhyale Hox* cluster is a single cluster. *Ubx* and *Abd-B* transcription spots should be nearly co-localized within the nucleus. C) *In situ* HCR of *Ubx* and *Abd-B* in a WT embryo at S19. C’) Inset region of Mx2, T1, T2, and T3 segments in embryo from panel C. D) *In situ* HCR of *Ubx* and *Abd-B* in a *Sce* S19-like mutant embryo. Arrows in *Ubx* and *Abd-B* panels mark regions of misexpression. D’) Inset region of Mx2, T1, T2, and T3 segments in embryo from panel D. D’’) Inset region from panel D’ showing *Ubx* misexpression spots in a cell found in the Mx2 segment. D’’’) Inset region from panel D’ showing *Ubx* and *Abd-B* misexpression in cells found in the T2 segment. E) Schematic representation of *Ubx* and *Abd-B* misexpression patterns in S17-like and S19-like *Sce* mutant embryos. All S17-like mutants showing misexpression appeared to show only *Abd-B* misexpression. The single S19-like mutant embryo showed both *Ubx* and *Abd-B* misexpression.

When we examined the expression of *Ubx* at S19 in knockout embryos, we began to see misexpression anterior to the WT expression boundary. Among the embryos we dissected at the S19 developmental stage, some embryos appeared to have developmental delay such that they had not yet formed the ventral furrow characteristic of S19 embryos. We labeled these embryos “S17-like” based on their morphology, and embryos with a more typical morphology “S19-like”. Among S19-like embryos, we observed spots of *Ubx* misexpression anterior to the WT boundary (*Sce*: 1/3 embryos; *Pcl*: 3/3 embryos; Fig. 6C, D, Supp. Fig. 6D-D’’). The misexpression seemed to be localized primarily to the neurogenic ectoderm. These results suggest that PcG genes are dispensable for wildtype *Ubx* anterior boundary establishment, but are essential for anterior boundary maintenance.

We observed a difference in the timing of misexpression relative to anterior boundary establishment when we examined *Abd-B* expression in knockout embryos at S19, in both S17-like and S19-like embryos. In S17-like embryos, we observed that *Abd-B* misexpression appeared to begin nearly concurrently with the early stages of *Abd-B* expression at the wildtype boundary (*Sce*: 3/5 embryos; *Pcl* 3/3 embryos; Fig. 6B, B’; Supp. Fig. 6.1C, C’). Moreover, S17-like embryos displayed *Abd-B* misexpression, but not *Ubx* misexpression. All S19-like embryos expressing *Ubx* also misexpressed *Abd-B* (Fig. 6E, Supp. Fig. 6.1E). These results suggest that the timing of *Abd-B* dependence on PcG function may differ from that of *Ubx*; namely, that *Abd-B* dependence on PcG function precedes *Ubx* dependence, and that *Abd-B* PcG-dependent boundary maintenance occurs simultaneously with wildtype boundary establishment.

### Tissue- and PcG knockout gene-specific effects on *Hox* expression

As development progresses to S21, we observed misexpression of *Ubx, abd-A,* and *Abd-B* in mutant embryos (Supp. Fig. 6.2). For *Sce* mutant embryos, we observed broad misexpression of *Ubx* in both limbs and the nervous system, weak misexpression of *Abd-B* in the nervous system, and very weak misexpression of *abd-A* in a few cells of the nervous system. The expression of these genes appeared to differ relative to that of *Pcl* mutant embryos, which displayed strong and broad misexpression of *Ubx* in both limbs and the nervous system, strong misexpression of *Abd-B* in the nervous system, and weak misexpression of *abd-A* in the nervous system. Thus, by the S21 stage, we began to observe differences in the effects of individual PcG gene knockouts on the expression of individual *Hox* genes.

At S23, the differences between *Sce* and *Pcl* mutant embryos became more dramatic (Fig. 4). While both knockouts displayed strong and broad *Ubx* misexpression, as well as strong *Abd-B* expression in the nervous system, we observed striking differences in the degree of *abd-A* misexpression between mutants. In *Sce* mutants, a small number of cells, restricted to the neurogenic ectoderm, appeared to show strong *abd-A* misexpression. In contrast, in *Pcl* mutants, we observed strong misexpression of *abd-A* all across the neurogenic ectoderm, as well as patchy misexpression in the limbs. The difference in these phenotypes suggests that *Pcl* is indispensable for proper *abd-A* repression, while *Sce* more weakly represses *abd-A*, indicating that individual PcG genes may have different roles in regulating individual *Hox* genes.

To assess the tissue localization of the apparent *Hox* misexpression, we performed *in situ* HCR on *abd-A* and *Abd-B* along with the pan-neural marker *elav* in *Sce* and *Pcl* knockout embryos (Fig. 7A, B; Supp. Fig. 7.1). We observed restriction of the *Abd-B* misexpression region to the neurogenic ectoderm, as we had imputed based on morphological evidence. Moreover, we confirmed that *abd-A* misexpression in *Sce* mutants is primarily restricted to small patches of cells in the nervous system, while *abd-A* misexpression in *Pcl* mutants expands into both neural and limb tissue.

**Fig. 7:**
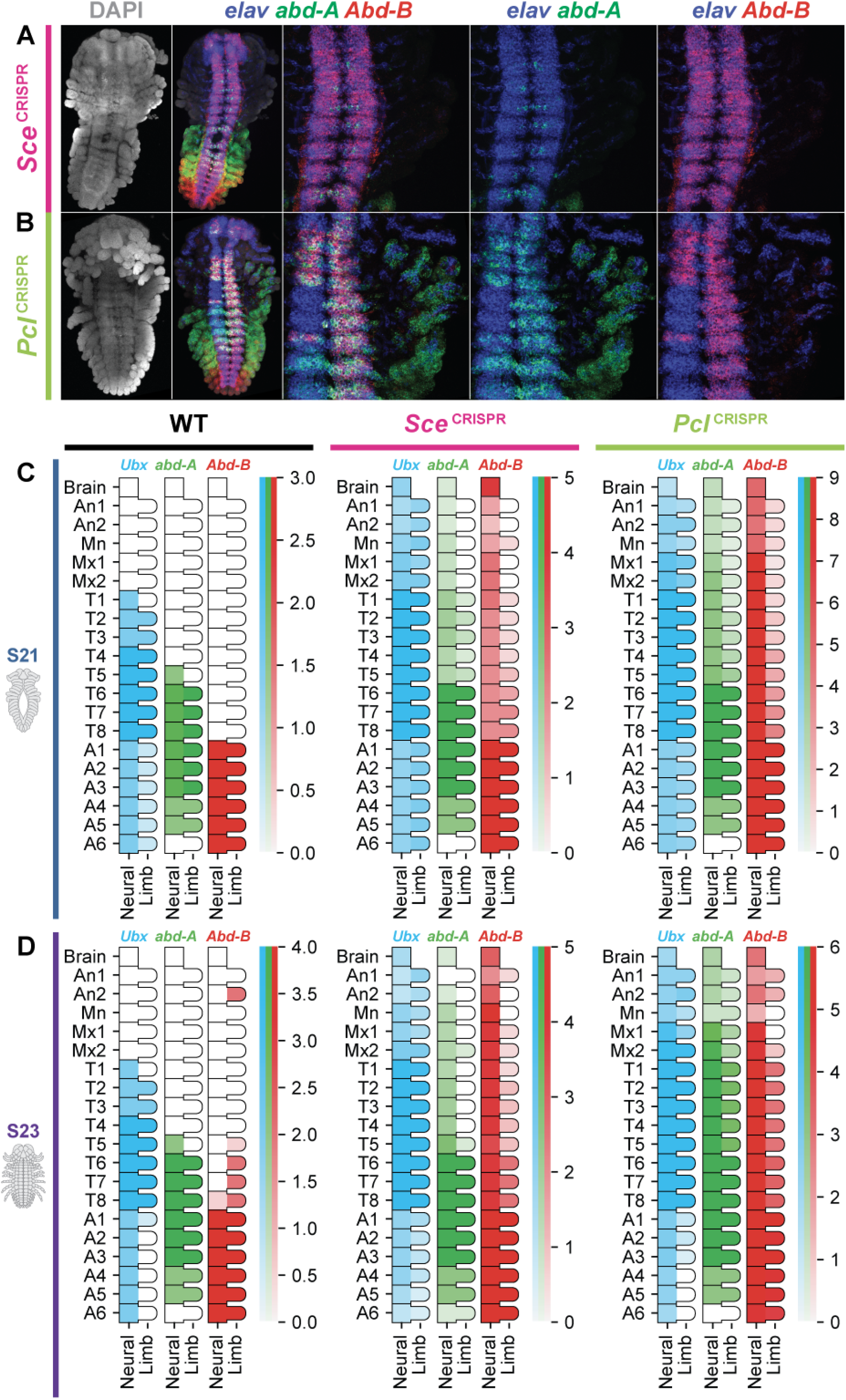
Differential tissue-specific misexpression of *Hox* genes between PcG mutants. A) *In situ* HCR of *abd-A* and *Abd-B* as well as the neural marker *elav* in a *Sce* mutant embryo at stage 23. *Abd-B* misexpression appears to be restricted to the nervous system. *abd-A* misexpression is found in a small number of nervous system cells. B) *In situ* HCR of *abd-A* and *Abd-B* as well as the neural marker *elav* in a *Pcl* mutant embryo at stage 23. *Abd-B* misexpression appears to be primarily found in the nervous system. *abd-A* misexpression, in contrast to *Sce* mutants, is stronger in the nervous system and also occurs in patches of the limbs. C) Quantification of *Ubx*, *abd-A*, and *Abd-B* expression in the limbs and nervous system of each segment of the body in WT, *Sce*, and *Pcl* embryos at stage 21. A region containing strong expression equivalent to the strongest WT expression was assigned a value of 1; a region containing patchy expression or expression less than the strongest WT expression was assigned a value of 0.5. 3 WT embryos, 5 *Sce* mutants, and 9 *Pcl* mutants were examined. D) Quantification of *Ubx*, *abd-A*, and *Abd-B* expression in the limbs and nervous system of each segment of the body in WT, *Sce*, and *Pcl* embryos at stage 21. Analysis performed identically to panel C. 4 WT embryos, 5 *Sce* mutants, and 6 *Pcl* mutants were examined. Differences were observed between *abd-A* and *Abd-B* expression in *Sce* and *Pcl* mutants. *abd-A* expression in *Sce* mutants is weaker in the nervous system and largely absent from the limbs, as compared to in *Pcl* mutants. *Abd-B* misexpression was stronger in the limbs of *Sce* mutants, but was largely found in strips reminiscent of limb enervation.

**Fig. 8:**
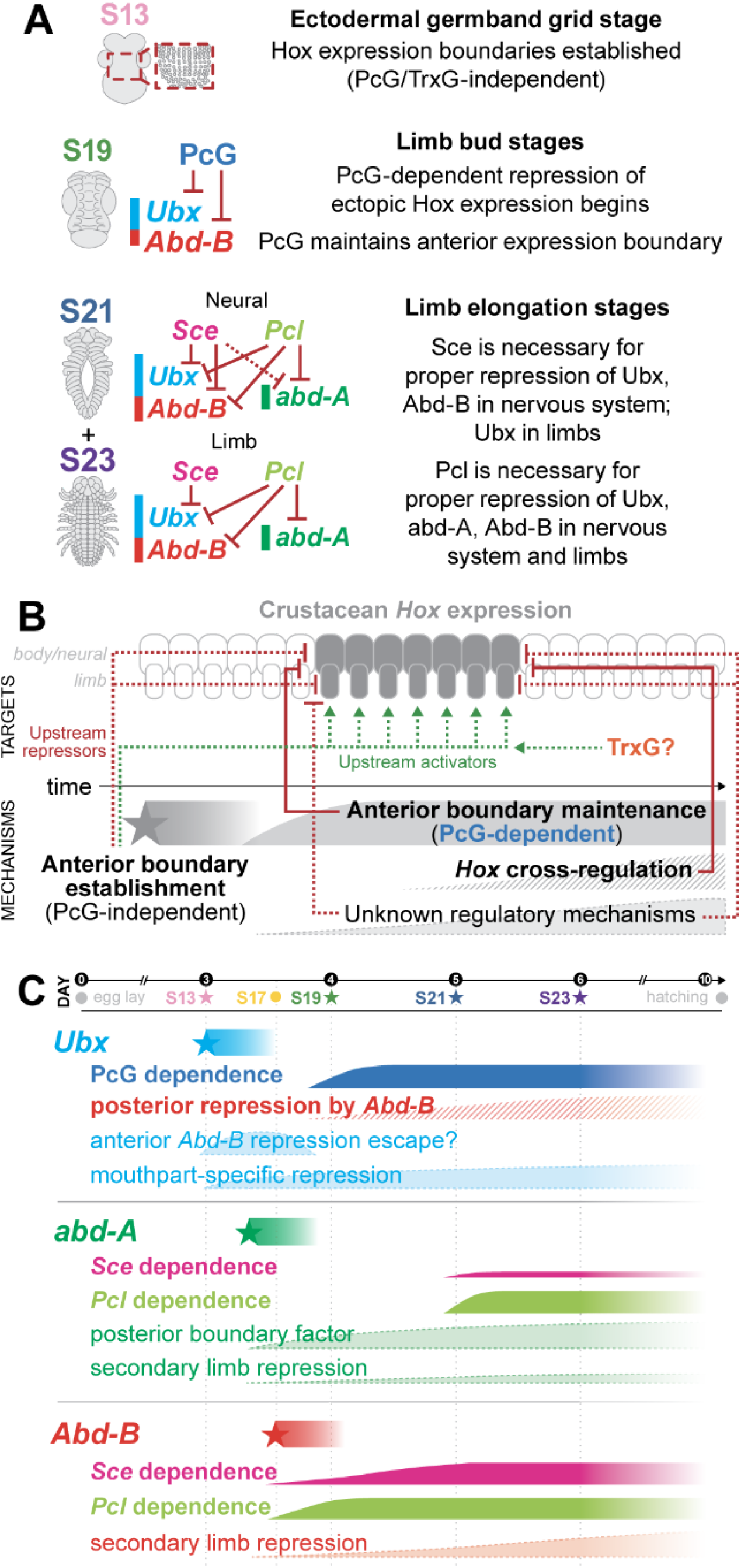
Summary of findings and implications for *Hox* regulatory evolution. A) Summary of the phenotypes and inferred genetic relationships of *Hox* genes and PcG genes at S13, S19, S21, and S23. B) Model of *Hox* regulation in crustaceans consisting of at least three mechanisms: anterior boundary establishment, anterior boundary maintenance (PcG-dependent), and *Hox* cross-regulation. Comparison of the results of PcG knockout to *Hox* gene knockout suggests that additional unknown regulatory mechanisms may also be involved in crustacean *Hox* regulation. C) Summary of the *Hox* regulatory mechanisms for each of the three posterior *Hox* genes in *Parhyale*. *Ubx* anterior boundary establishment occurs at around S13, and the PcG-dependent phase of *Ubx* expression begins around S19. The inability of ectopic *Abd-B* to repress *Ubx* in anterior tissues suggests that anterior *Ubx* expression may achieve an autoregulatory state, preventing repression by *Abd-B*. The mutually exclusive expression domains of *Ubx* and *Scr* in the mouthparts suggest differential competence of mouthpart tissue to express *Ubx*. *abd-A* establishes its *Hox* expression between S13 and S17, but does not appear to become dependent on PcG repression until around S21. *abd-A* is differentially dependent on *Sce* and *Pcl* function for proper repression. An unknown factor may be responsible for the posterior boundary of *abd-A* expression, which is not perturbed by either *Abd-B* knockout or PcG loss of function. *Abd-B* establishes its anterior boundary and becomes dependent on PcG function at S19. *Abd-B* misexpression induced by PcG knockout appears to be primarily restricted to neural tissue. An unknown mechanism may repress *Abd-B* expression in the anterior limbs.

Based on these data, we further quantified the degree of *Hox* misexpression for *Ubx*, *abd-A*, and *Abd-B* across our HCR-stained wildtype and knockout embryos. For each segment of the body, we evaluated the expression of each of the three posterior *Hox* genes in both the medial portion of the embryo containing the neurogenic ectoderm, and the lateral regions containing limb tissue. We assigned a value of 1 to a wildtype-like high level of *Hox* expression, a value of 0.5 to expression below the maximum wildtype expression or for patchy or partial expression, and a value of 0 to absence of expression. The results of this quantification are summarized in Fig. 7C for S21 embryos and Fig. 7D for S23 embryos. Notably, we observed that *abd-A* expression appeared to be lower in the nervous system in *Sce* knockout embryos at both stages S21 and S23 when compared to *Pcl* knockout embryos. We also observed that *abd-A* appeared largely absent from the limbs in *Sce* knockout embryos, but showed strong expression in *Pcl* knockout embryos, further confirming our observation that *Sce* and *Pcl* appear to have different roles in regulating *abd-A*.

These results together reveal that individual *Hox* genes in *Parhyale* appear to be regulated by distinct genetic mechanisms. For *Ubx*, both *Sce* and *Pcl* mutants showed similarly strong misexpression in the nervous system and limbs. *Abd-B* misexpression appeared to be concentrated in the nervous system in *Sce* knockout embryos, with slight misexpression in the limbs – for *Pcl* knockout embryos, we also observed strong misexpression in the nervous system and slightly higher *Abd-B* misexpression in the limbs. Among the three posterior *Hox* genes, *abd-A* showed the strongest difference in effect between *Pcl* and *Sce* knockout embryos, with weak, neurally-restricted misexpression induced by *Sce* knockout and broad, strong misexpression induced by *Pcl* knockout. These results suggest that each of the three posterior *Hox* genes have differential dependence on the function of individual PcG genes, and that the role of PcG regulation in the limbs and nervous system also differs between *Hox* genes.

## Discussion

In this paper, we reveal the essential roles of PcG genes in maintaining proper *Hox* expression in the amphipod crustacean *Parhyale hawaiensis*. Loss of PcG function induces *Hox* misexpression and homeotic transformations in a manner similar to that observed in other systems. However, the effects of PcG loss of function appeared to manifest differently for individual *Hox* genes, in different regions of the embryo, and for individual PcG genes. The striking differences in the effects of PcG knockout suggest idiosyncratic regulation of *Hox* genes by PcG group genes. Moreover, the phenotypes we observed across tissues and timepoints also provided additional insights into the mechanisms of *Hox* regulation in this organism.

### Proper *Hox* regulation in *Parhyale* requires at least three distinct mechanisms

The phenotypes we observed, when considered in relation to previous experiments, suggest that there are at least three distinct regulatory mechanisms involved in proper *Hox* regulation in *Parhyale*: boundary establishment, boundary maintenance, and *Hox* cross-regulation. These results are consistent with previous work on *Hox* regulation in *Drosophila* (Akam, 1987; Mallo and Alonso, 2013).

#### 1. A PcG-independent mechanism governs early *Hox* anterior boundary establishment

First, our data suggest that an early boundary establishment mechanism precedes the PcG-dependent phase of *Hox* regulation in *Parhyale*. In PcG knockout embryos at the early germband stage (S13), we consistently observed proper establishment of the anterior expression boundary of *Ubx*. This suggests that PcG genes are dispensable for the proper establishment of the anterior expression boundary, and that the early regionalization of the *Parhyale* embryo is governed by other mechanisms.

In insects, gap and pair rule transcription factors are the upstream regulators of *Hox* genes, and are responsible for anterior boundary establishment (Maeda and Karch, 2006, 2015). While the precise determinants of anterior boundary establishment in *Parhyale* remain unknown, a similar phase of regulation prior to PcG dependence must also proceed during early development. We hypothesize that a family of transcription factors may act in a similar function to the gap and pair-rule genes in insects to establish the early anterior expression boundaries of *Hox* genes in *Parhyale*. Future experiments using techniques such as single-cell RNA sequencing may provide opportunities to identify such factors.

#### 2. A PcG-dependent mechanism governs *Hox* anterior expression boundary maintenance

Second, our data clearly demonstrate that, following the boundary establishment phase of *Hox* expression, PcG genes become essential for proper anterior boundary maintenance. For the five *Hox* genes we examined (*Scr*, *Antp*, *Ubx*, *abd-A*, and *Abd-B*), we observed evidence of *Hox* misexpression anterior to the wildtype anterior expression boundary. This result demonstrates a second phase of *Hox* regulation, wherein PcG function is required to maintain proper *Hox* repression. The degree of PcG dependence appeared to vary at the level of tissue location and PcG gene for each of the genes we examined.

The observation of two phases of early *Hox* regulation in *Parhyale* is notable when compared to previous experiments in insects. In the long-germ insect *Drosophila*, the boundary establishment and boundary maintenance phases of *Hox* regulation appear to occur in an ordered sequence, wherein gap and pair rule genes serve as upstream inputs to *Hox* regulation, and PcG genes later become essential for *Hox* repression (Jaeger, 2011; Simon et al., 1992; Struhl and Akam, 1985). In contrast, in the short-germ insect *Gryllus bimaculatus*, RNAi knockdown of PRC2 complex components *E(z)* and *Esc* appeared to suggest that the *Hox* genes *abd-A* and *Abd-B* lacked the gap gene-based establishment phase of *Hox* expression (Matsuoka et al., 2015). The authors made this assertion based on data indicating that anterior *Hox* genes (*Scr*, *Antp*, *Ubx*) in PRC2-RNAi embryos displayed wild-type anterior expression boundaries early in development, followed by expansion of expression later in development. In contrast, for the posterior *Hox* genes *abd-A* and *Abd-B*, PRC2-RNAi embryos exhibited much earlier misexpression, which appeared at the same time that the wildtype expression boundary would be observed in WT embryos.

We observed a similar contrast in the potential timing of boundary establishment and boundary maintenance between *Ubx* and *Abd-B* in *Parhyale*. Namely, the *Ubx* anterior expression boundary in PcG knockouts appeared normal at early stages (∼S13), while misexpression began later in development (∼S19). In contrast, for *Abd-B*, misexpression appeared to begin concurrently with the wildtype expression boundary (∼S17). These data in *Parhyale* could indicate a similar mechanism to that observed in *Gryllus*: that anterior and posterior *Hox* genes have different boundary maintenance and boundary establishment requirements. However, in *Parhyale*, while *Abd-B* misexpression appeared to occur simultaneously with normal boundary establishment, the tissue location and intensity of expression differed. *Abd-B* misexpression appeared primarily in the neurogenic ectoderm at that early stage, and the broad normal anterior boundary was also clearly visible. Thus, it is possible that, despite the simultaneous appearance of the normal anterior boundary and ectopic expression, both mechanisms could still be required for *Parhyale Abd-B*.

Our results suggest that the previous data in *Gryllus* could benefit from reinvestigation. For *Gryllus abd-A* and *Abd-B*, the colorimetric *in situ* hybridization results presented show strong misexpression at developmental stages concurrent with strong wildtype expression; however, it is possible that at slightly earlier stages, one might still observe a stronger and clearer normal expression boundary, as we observed in *Parhyale*. Using a more sensitive technique such as *in situ* HCR at earlier stages could reveal the potential for a simultaneous biphasic regulatory mechanism, as our results in *Parhyale* suggest.

#### 3. PcG knockouts reveal conditional cross-regulation of *Hox* genes

Third, our results provide further insight into the complexities of *Hox* cross-regulatory mechanisms in *Parhyale*. Previous studies have revealed that *Hox* genes in *Parhyale* exhibit cross-regulation. For example, CRISPR-Cas9 knockout of *Abd-B* results in posterior expansion of *Ubx* expression, and homeotic transformations of abdominal segments to walking and jumping legs (Martin 2016, Jarvis 2018). This indicates that *Abd-B* represses *Ubx*. In a different set of studies, heat shock-induced misexpression of *Ubx* resulted in decreased *Scr* expression, indicating that *Ubx* represses *Scr* (Pavlopoulos et al., 2009). In our study, we observed contrasting effects of PcG knockout on the *AbdB* repression of *Ubx* and the *Ubx* repression of *Scr*.

In PcG knockout embryos, *Abd-B* appeared to properly repress *Ubx* in the abdomen of the embryo, while in regions where *Abd-B* was ectopically expressed, *Ubx* did not appear to be ectopically repressed. In wildtype embryos, *Ubx* expression is strongly observed in cells that begin expressing *Abd-B* at S17, and gradually grows weaker after S19, such that *Ubx* expression in the abdominal limbs and nervous system is weak but still observable by S21. By S23, *Ubx* expression in wildtype embryos is not observable in abdominal limbs, but weakly persists in some cells of the nervous system.

In PcG knockouts, the same trend is observed: strong *Ubx* expression in *Abd-B*-expressing cells of the abdomen, which gradually tapers into weak expression restricted to the nervous system by S23. In the same PcG knockouts, *Abd-B* misexpression begins with a few cells of the neurogenic ectoderm at early stages (S17), expanding to encompass much of the nervous system of all anterior segments by S23. Despite this strong expression of *Abd-B*, *Ubx* expression does not appear to be repressed in the nervous system. Cells expressing both *Ubx* and *Abd-B* can be found even at S23 in the nervous system of PcG knockout embryos. Thus, despite the fact that both anterior neural cells and posterior abdominal tissue contain cells that simultaneously express *Ubx* and *Abd-B* beginning at S17, *Abd-B* only represses *Ubx* in the abdomen, and not in the anterior neural tissue where it is ectopically expressed.

One explanation for this observation may be that *Abd-B* repression of *Ubx* depends on some third factor expressed in abdominal tissue, and that *Abd-B* repression by *Ubx* in the abdomen is not dependent on PcG function. Alternatively, anteriorly expressed *Ubx* could undergo auto-regulation, thus escaping later ectopic *Abd-B* expression. Heat shock misexpression of *Abd-B* could more precisely reveal whether *Abd-B* repression of *Ubx* in anterior tissue is dependent on PcG function.

PcG knockout embryos also revealed further complexity in the repression of *Scr* by *Ubx*. In PcG mutants, *Scr* expression in T1 is substantially reduced, likely due to strong misexpression of *Ubx*. Moreover, *Scr* misexpression in the antennae is observed in only a few cells, potentially also due to strong *Ubx* misexpression in those tissues. These results indicate that, in the antennae and T1 appendages, *Ubx* repression of *Scr* is not dependent on PcG function.

A more complicated mechanism appears to take place in the Mn, Mx1, and Mx2 appendages, where mutually exclusive *Scr* and *Ubx* expression domains are observed. In anterior regions of these limbs that ectopically express *Ubx*, *Scr* expression is almost completely eliminated. However, the posterior regions of these limbs appear to retain *Scr* expression. In these tissues, it is possible that either *Ubx* requires PcG function to repress *Scr*, or that some other mechanism prevents ectopic *Ubx* expression, such as the presence of a *Ubx* repressor or absence of *Ubx* activators. Thus, these results indicate that *Hox* cross-regulation may depend on factors found only in particular regions of an individual body segment.

Together, these data indicate that *Hox* cross-regulation occurs in *Parhyale* via layered mechanisms, potentially dependent on tissue type, expression timing, and body region. Future experiments using knockouts of *Hox* genes and PcG genes, or heat shock induced misexpression combined with PcG knockout, could clarify the role of PcG function in *Hox* cross-regulation.

### Each posterior *Hox* gene in *Parhyale* exhibits idiosyncratic regulation

In addition to revealing insights into broader *Hox* regulatory mechanisms, our results also suggest that individual *Hox* genes in *Parhyale* follow idiosyncratic regulation. The three posterior *Hox* genes showed differences in the timing of their dependence on PcG function, the tissues in which ectopic expression was observed, and their dependence on specific PcG complex genes.

#### Differences in relative *Hox* misexpression timing

We were able to characterize the timing of misexpression for *Ubx* and *Abd-B*, and observed differences in the relative timing of *Hox* misexpression in relation to the normal anterior boundary establishment. *Ubx* showed no misexpression at the time of *Hox* initiation (S13), and did not exhibit observable misexpression in segments more anterior than the normal anterior expression boundary until around S19. In contrast, *Abd-B* showed simultaneous misexpression and normal anterior boundary establishment at S17. These results suggest that *Ubx* and *Abd-B* become dependent on PcG function at different times during development.

#### Differences in tissues competent for ectopic expression

While *Ubx* became strongly misexpressed upon knockout of PcG genes, *Abd-B* misexpression appeared to exhibit tissue dependence. For both *Pcl* and *Sce* knockout, *Abd-B* became strongly misexpressed in the nervous system, but did not appear to exhibit strong misexpression in the limbs. This result suggests that PcG may not be the primary method of repression for *Abd-B* in the limbs, and that additional mechanisms may be needed to explain how *Abd-B* is expressed within the limbs.

The presence of differential expression of *Hox* genes along the proximal-distal axis (in this case, the medial neurogenic ectoderm and the more distal limbs) is not without precedent in *Parhyale*. Indeed, more anterior *Hox* genes such as *Scr* appear to have differential regulation along the proximal-distal axis. *Scr* expression in the T1/Mxp appendage is restricted to the limb, and is absent from the nervous system. This *Scr* expression is critical to T1/Mxp identity, as revealed in knockout experiments. Our results indicate that a similar partitioning of limb and nervous system expression potential also exists for *Abd-B*. The precise mechanisms of this proximal-distal regulation remain to be investigated.

Together, these data suggest that proximal-distal regulation of *Hox* expression may be a general feature of the crustacean *Hox* regulatory architecture. Moreover, the existence of such mechanisms provide another regulatory parameter for the fine tuning of *Hox* expression, which may have played a role in the evolution of different expression patterns across crustaceans.

#### Differences in PcG complex gene dependence

Among the three genes we examined, *abd-A* appeared to show a striking difference in its dependence on different PcG genes for proper regulation. The phenotypes for PcG knockout-induced *Ubx* and *Abd-B* misexpression appeared similar in *Sce* and *Pcl* mutants; however, they differed substantially in the case of *abd-A*. *Sce* knockout induced misexpression in a few cells of the nervous system, whereas *Pcl* knockout induced strong and broad *abd-A* misexpression in the nervous system and strong but patchy misexpression in the limbs.

These results indicate that individual PcG genes may have gene-specific regulatory functions for individual *Hox* genes. In the case of *Sce* and *Pcl*, members of PRC1 and PRC2.1 respectively, complex-specific functions may explain the differences in phenotypes. For example, PRC1 is responsible for H2AK188 ubiquitylation, whereas PRC2 is responsible for H3K4 trimethylation. Thus, it is possible that H2AK188ub is largely dispensable for *abd-A* regulation, whereas H3K4me3 is essential. Immunofluorescence experiments on PcG knockout embryos could reveal such differences, while ChIP-Seq or CUT&RUN experiments could reveal the precise cis-regulatory targets of these genes.

The presence of differential regulation of *Hox* genes by PcG genes reveals that the upstream factors governing *Hox* regulation in *Parhyale* may have become distinct. Alterations to the function of *Sce*, for example, could preferentially result in changes to *Ubx* and *Abd-B* expression, without altering *abd-A* expression. While we only examined two PcG genes in detail in our study, it is possible that other PcG genes may also have regulatory functions that preferentially affect or exclude individual *Hox* genes. Such regulatory mechanisms would provide a direct method to alter the expression of a single *Hox* gene while leaving the expression of other *Hox* genes intact, providing a potential mechanism for the evolution of new *Hox* expression patterns.

#### PcG genes are crucial for diverse developmental processes in *Parhyale*

In addition to homeotic phenotypes and effects on *Hox* regulation, we characterized a number of additional roles for PcG genes during *Parhyale* development. In particular, we observed defects in the development of body and limb plates, as well as defects in proximal-distal patterning, suggesting that PcG genes may also play important roles in these developmental processes.

PcG knockout embryos displayed reduced plates reminiscent of those observed in knockouts of genes crucial for the development of insect wings, such as *vestigial*, *nubbin*, and *apterous*, which resulted in coxal, tergal, and basal plates of reduced size. This suggests that PcG genes may regulate such plate developmental genes. For one gene, *pho*, we observed potential homeotic transformations of plates, suggesting that PcG genes may also play a role in plate identity. We also observed phenotypes indicating that PcG genes play a role in the closing of the dorsal side of the embryo, as we observed tergal plates that failed to fuse dorsally, fused asymmetrically between left and right halves of the embryo, or fused adjacent segments along the anterior-posterior axis. The genes involved in dorsal closure in *Parhyale* remain uninvestigated, but we expect such genes are likely also regulated by PcG function.

Moreover, the consistent loss of the T2 coxal and tergal plates across PcG genes reveals subdivisions of regions of *Hox*-specified identity in the *Parhyale* embryo. Both T2 and T3 limbs in *Parhyale* develop into a claw, or cheliped. However, knockout of PcG frequently ablates the entire T2 tergal and coxal plates, in addition to inducing T2 -> T4/5 transformations. In the T3 segment, which also consistently had homeotic transformations of T3 -> T4/5 identity, we sometimes observed failure of fusion of the tergal plate, but we did not observe total loss of the tergal and coxal plates. Thus, PcG knockout reveals that distinct genetic mechanisms may separate body segments with similar homeotic identity, revealing additional layers of distinction that subdivide body segments with similar *Hox* expression.

Finally, we observed defects in proximal-distal development for most of the PcG knockouts we examined, including truncation of limbs and fusion of limb segments. For some of these phenotypes, it was possible to infer the specific limb segments that were lost or fused. For example, *Sce* knockout appears to induce variable fusion of the propodus, carpus, and merus. Such fusions are not identical to any of the phenotypes previously described for the leg gap gene or *Hox* gene knockouts in *Parhyale*. These results indicate that PcG genes may work in concert with leg gap genes, and other unknown genes, to specify proximal-distal identity along the legs in *Parhyale*.

#### TrxG proteins show no obvious role in *Parhyale Hox* regulation

We were unable to recover any homeotic phenotypes for the TrxG genes we targeted using CRISPR-Cas9 mutagenesis. Some of the genes we analyzed displayed their roles in *Hox* regulation in suppressor screens of PcG genes in *Drosophila* – performing combined knockouts of these genes and PcG genes in *Parhyale* could clarify their roles in *Hox* regulation.

The absence of any obvious phenotypes for *Trx* is somewhat surprising, given the phenotypes observed in *Drosophila* knockouts, which show homeotic transformations of haltere to wing structures (Breen and Harte, 1991, 1993). It is possible that the function of *Trx* could be replaced in *Parhyale* by *Trithorax-related* (*Trr*), which we also found in the *Parhyale* genome, which would be an interesting question for a future study. More careful analysis of the *Trx* locus, including long-read RNA-sequencing, RACE, or other experiments that examine the exact sequence content of *Trx* transcripts, could also clarify whether *Trx* in *Parhyale* is indeed expressed as two separate transcripts, and lead to clearer understanding of the function and structure of this gene.

## Ideas and Speculation

When considered alongside previous data in this organism, our results provide a number of interesting testable hypotheses and avenues for future investigation. In particular, we propose that currently undiscovered mechanisms, both ancestral to arthropods and evolved in crustaceans, may play a role in crustacean *Hox* regulation. We also examine the implications of our data in relationship to previous studies characterizing the functions of PcG genes in other organisms.

### PcG knockout phenotypes suggest a limb-specific repression mechanism for *Abd-B*

While we observed strong and broad misexpression of *Ubx* in both limbs and the nervous system in response to PcG knockout at late stages (S21-S23), *Abd-B* misexpression was mostly restricted to the nervous system for both *Sce* and *Pcl* knockouts. This result contrasts with PcG knockout phenotypes observed in *Drosophila*, where broad *Abd-B* misexpression and homeotic transformations of body segments towards the *Abd-B*-specified A8 identity have been observed. The absence of strong *Abd-B* misexpression in the limbs of PcG knockout *Parhyale* suggests the presence of a secondary limb-specific repression mechanism. Such a mechanism could be either the absence of an essential *Abd-B* activator, or the presence of an additional *Abd-B* repressor in limb tissue.

This observation reveals potential insights into crustacean *Hox* regulatory evolution when compared to the results found in *Gryllus bimaculatus* PRC2 knockdown experiments (Matsuoka et al., 2015). In those studies, *Abd-B* misexpression also appeared to be restricted to the neurogenic ectoderm, although the authors do not make any special note of this phenotype. Given that *Abd-B* is not normally expressed in any limb tissue in insects, the absence of *Abd-B* expression in limbs may not be surprising for this organism. However, the phenotypes observed in *Parhyale* provide additional context, namely suggesting that an ancestral secondary mechanism for *Abd-B* repression in thoracic limbs may be shared between crustaceans and short-germ insects. This secondary repression mechanism may then have been lost in long-germ insects, such as *Drosophila*, enabling broader derepression of *Abd-B* in response to loss of PcG function (Supp. Fig. 8.1A).

### PcG knockouts reveal gene-specific roles in *abd-A* regulation and an unknown posterior boundary maintenance mechanism

In addition to the differences in effect for individual PcG gene knockouts on *abd-A*, it is also notable that the posterior boundary of *abd-A* appears to remain intact in PcG mutants. In *Drosophila*, while the anterior boundaries of *Hox* expression are maintained by PcG function, the posterior boundaries of *Hox* expression are thought to be maintained by *Hox* cross-regulation. In *Parhyale*, this posterior boundary maintenance appears to be true for *Abd-B* repression of *Ubx* and *Ubx* repression of *Scr*. However, previous knockout experiments have shown that neither *Ubx* nor *Abd-B* is necessary for the proper posterior boundary maintenance of *abd-A*, suggesting no role of *Hox* cross-regulation in this process (Jarvis et al., 2022). Our data indicate that PcG knockout also does not disrupt the posterior boundary of *abd-A* expression. Thus, an additional, unknown mechanism governs the position of the posterior boundary of *abd-A* expression which is independent of known *Hox* maintenance mechanisms in arthropods.

The existence of such a mechanism could provide an explanation for the divergent expression patterns of *abd-A* in different groups of crustaceans. For example, in the decapod crustacean *Procambarus fallax* and the mysid *Mysidium columbiae*, *abd-A* is strongly expressed in segments A4 and A5, whereas its expression is weaker in *Parhyale*. The evolution of a novel, gradated *abd-A* repression mechanism could explain the shift in *abd-A* expression in *Parhyale* and other amphipods, providing a mechanistic explanation for crustacean body plan evolution. We propose that a Posterior Boundary Factor may govern the regulation of the posterior boundary of *abd-A* in crustaceans, and that this factor operates independently of *Hox* cross-regulatory mechanisms (Supp. Fig. 8.1B). This factor may be one of the many additional PcG genes for which phenotypes have not been thoroughly investigated, or it may be another mechanism entirely. Such a Posterior Boundary Factor, if identified, could represent a novel regulatory mechanisms evolved within crustaceans.

### Implications for the cis-regulatory architecture of the *Parhyale Hox* cluster

Our experiments demonstrate a clear role for PcG genes in *Parhyale Hox* regulation. Given the current understanding of PcG function in *Drosophila* and its relationship to cis-regulatory elements, we can make certain hypotheses about the cis-regulatory architecture of the *Parhyale Hox* cluster. In particular, our data imply the existence of Polycomb Response Elements (PREs) in the *Parhyale Hox* cluster. PREs are specialized cis-regulatory elements that recruit *pho* binding and later PcG complex activity to a locus in *Drosophila*.

Future studies using ChIP-Seq or CUT&RUN, combined with newly available functional genomic data for *Parhyale*, such as ATAC-Seq (Sun et al., 2021), could reveal the precise location of PREs and other important cis-regulatory elements in the *Parhyale Hox* cluster. Such approaches could then enable the precise dissection of cis- and trans-regulatory mechanisms of *Hox* regulation, and enable direct comparisons of the *Parhyale Hox* cluster to the more well-studied *Drosophila Hox* cluster.

### Implications for the ancestral roles of PcG in *Hox* regulation

Previous work in flies and mice has revealed two different potential roles for *PcG* regulation of *Hox* genes. In *Drosophila*, *Hox* boundary establishment occurs independently of PcG function, and PcG genes are required for proper boundary maintenance (Maeda and Karch, 2006, 2015). In vertebrates, boundary establishment and maintenance both appear to depend on PcG/TrxG-mediated chromatin states (Chambeyron and Bickmore, 2004; Deschamps and van Nes, 2005; Kmita and Duboule, 2003; Rastegar Mojgan et al., 2004; Soshnikova and Duboule, 2009; Tschopp and Duboule, 2011).

The previous study in the cricket *Gryllus bimaculatus* suggested that *Gryllus* may represent an intermediate between the mouse and fly literature, wherein anterior *Hox* genes depend on PcG for boundary maintenance, but not boundary establishment, whereas *abd-A* and *Abd-B* have a vertebrate-like dependence on PcG for both boundary establishment and maintenance (Matsuoka et al., 2015).

Our data suggest that more careful examination of the timing and intensity of misexpression could reveal that *Hox* genes require PcG only for boundary maintenance, rather than both maintenance and establishment, in short-germ arthropods such as *Gryllus* and *Parhyale*. Notably, short-germ arthropods have progressive segmentation of the body, where anterior regions segments develop prior to more posterior segments. Thus, within an embryo, anterior segments are “older” than posterior segments, and often further along in developmental processes such as limb bud formation and *Hox* expression. In PcG loss of function experiments in both *Gryllus* and *Parhyale*, ectopic *Hox* expression in anterior tissues occurs concurrently with appearance of the wildtype boundary. The previous study in *Gryllus* argued that this result suggested that *Gryllus abd-A* and *Abd-B* did not utilize a PcG-independent boundary establishment phase, and instead used a vertebrate-like PcG-dependent boundary establishment mechanism. We favor the interpretation that the PcG-dependent boundary maintenance phase begins in anterior tissue at the same time that more posterior tissue enters the PcG-independent boundary establishment phase in short germ insects such as *Gryllus* and *Parhyale*, and that the apparent concurrent timing is a result of the different “ages” of the anterior and posterior regions.

As our data do not directly identify upstream PcG-independent boundary establishment factors for *Parhyale Abd-B*, we provide two potential interpretations of our findings in the context of broader bilaterian *Hox* regulation by PcG (Supp. Fig. 8.1C). In the first hypothesis, *Parhyale Abd-B* behaves analogously to the previous interpretation of *Gryllus abd-A* and *Abd-B* regulation, wherein the more posterior *Hox* genes utilize a PcG-independent mechanism of boundary establishment. The second hypothesis, which we favor based on our interpretation of the early *Abd-B* misexpression pattern in *Parhyale*, suggests that all arthropods may deploy two phases of boundary regulation: PcG-independent boundary establishment, and PcG-dependent boundary maintenance.

In the case of the second hypothesis, a striking asymmetry is observed between arthropod and vertebrate *Hox* regulation by PcG genes, wherein arthropod PcG function is primarily necessary for *Hox* expression maintenance, rather than initial boundary establishment, while vertebrate PcG function is important for both establishing and maintaining *Hox* patterns. These results refute the assertion that a transition in *Hox* regulation occurred during arthropod evolution from an ancestral PcG-based *Hox* boundary establishment mechanism towards a gap-gene mechanism during arthropod evolution. Instead, our results suggest that understanding the ancestral role of PcG in regulating the urbilaterian *Hox* cluster will require further sampling at more early-branching protostome and deuterostome lineages.

## Conclusion

### Distinct regulatory strategies provide possible mechanisms for *Hox* regulatory evolution

Together, our PcG knockout and *Hox* expression data indicate that *Parhyale Hox* genes have meaningful differences in their upstream regulatory mechanisms. These differences can occur with respect to the timing of PcG dependence, tissue-specific regulatory mechanisms, and PcG gene-specific effects. The presence of distinct regulatory mechanisms for each *Hox* gene in *Parhyale* suggest that crustacean *Hox* genes more broadly could have evolved distinct *Hox* regulatory mechanisms, wherein perturbations to canonical regulation could have effects on individual *Hox* genes, at specific times in development, and in individual tissues, rather than affecting all *Hox* genes uniformly across the body.

In addition, the phenotypes we have observed, when considered in conjunction with previous work examining *Hox* cross-regulatory mechanisms, suggest that additional layers and mechanisms of regulation remain to be discovered within crustacean *Hox* clusters. Such unknown mechanisms may include: the affectors responsible for *Hox* anterior boundary establishment prior to PcG dependence; the activators and repressors governing *Ubx* expression potential in mouthparts; the secondary repression mechanism for *Abd-B* in the limbs; and the affectors maintaining the posterior boundary of *abd-A*. Identifying such factors and examining their function across crustaceans could lead to a deeper mechanistic understanding of the ancient processes underlying crustacean body plan evolution.

## Methods

### BLAST identification of PcG/TrxG genes

General strategy outlined in Supp Fig. 1.2b. We identified core PcG/TrxG genes from the *Drosophila* and *Mus* literature, with special attention to those genes that had previously been shown to induce homeotic phenotypes (Supp Tables 1 and 2). We downloaded peptide sequences for these genes from UNIPROT and used tblastn to identify potential best hits to the *Parhyale* genome. For each peptide, we used BLAST against 5 different *Parhyale* transcript sources: the *Parhyale* MAKER genome annotation (acquired from Leo Blondel), the Kao *et al*. Mikado transcriptome, the Sun *et al*. Mikado transcriptome, the Sun *et al*. StringTie2 short + long read embryonic transcriptome, and the Trinity-Limb transcriptome from the Sun *et al*. manuscript (generated by Heather Bruce and Jessen Bredeson). Several groups of genes appeared to have shared BLAST hits, including *Trx*, *Setd1*, and *Trr*. We disambiguated any inconclusive results using reciprocal best BLAST searches using the online BLAST portal. The full list of the *Parhyale* PcG/TrxG genes we identified is found in Supp. Tables 3 and 4.

### RNA-Seq expression quantification

We used RNA-Seq data from the Sun et al. 2022 manuscript for developmental stages S13, S19, S21, and S23 (three replicates for each stage). We quantified the expression of all transcripts using Kallisto alignment-free transcript abundance estimation to the Sun-Mikado transcriptome. For each PcG/TrxG gene, we used the best Mikado transcript BLAST hit.

### CRISPR-Cas9 guide design

We used the CRISPR-Cas9 targeting tool in Benchling to choose guide RNA sequences. We designed guides targeting either a 5’ portion of the ORF of a gene of interest, determined based on BLAST to all organisms to assess the true START codon, or to a specifically identifiable BLAST homology domain with suggested importance in gene function.

### CRISPR-Cas9 mutagenesis

Cas9 ribonucleoprotein complexes were assembled using Cas9-NLS enzyme purchased from the QB3 MacroLab at Berkeley and guide RNAs purchased from Synthego. Cas9 enzyme was used at a final concentration of 333 ng/µL and guide RNAs were used at a final concentration of 100ng/µL for each guide RNA in multi-guide knockout experiments, and 200ng/µL for single-guide knockout experiments. Final guide injection mix also included 0.05% phenol red; all injection components were mixed in nuclease-free water. For most experiments, we used two guide RNAs targeting each gene of interest. To demonstrate the reproducibility of single guide RNAs to generate the same phenotypes, thus controlling for potential off-target effects, we performed single-guide knockouts of *Sce*, *Pcl*, and *Su(Z)12* (see Supp. Figs. 2.2, 2.3, and 2.4) and quantified the proportion of phenotypes we observed between single and double-guide knockout experiments. We recovered the same classes of phenotypes between the single-guide and double-guide knockout experiments, although the relative proportion of each phenotype and total mutagenic efficacy varied.

### Embryo culture and dissection

Embryos were raised as previously described. After injection, embryos were transferred to filter-sterilized artificial sea water (FASW) made using Tropic Marin sea salt to a final salinity of ∼31-35%. Embryos were raised at 27°C in a humidified chamber in an incubator. We performed regular water changes on embryos every 1-2 days, removing any dead embryos. Wildtype embryos usually hatch by 10-11dpf. For our experiments, if any embryos remained unhatched but alive at 14dpf, we sacrificed and dissected those embryos using sharpened tungsten needles. For *in situ* hybridization and immunofluorescence experiments, we dissected embryos in 9 parts FASW, 1 part PEM buffer, and 1 part 32% paraformaldehyde and fixed for 25-35 minutes at room temperature (∼21°C) using previously described techniques.

### Microscopy and phenotype analyses

Hatchlings were fixed in 9 parts FASW, 1 part PEM buffer, and 1 part 32% paraformaldehyde. After removing FASW and applying the 9:1:1 fix, we used a tungsten needle to pierce the gut tube, usually through the ventral side of the hatchling around the juncture between the head and thorax. This “throat punch” technique largely prevented gross morphological distortions often observed in fixed hatchlings. Hatchlings were fixed for 1hr at room temperature in 9:1:1 fix, or overnight at 4°C. After fixation, hatchlings were washed 3X with PT and transferred to 50% glycerol/ 50% 1XPBS, then 70% glycerol/ 30% 1X PBS for mounting. For limb dissection panels, limbs were dissected in the 70% glycerol/ 30% 1X PBS mounting medium. Hatchlings were imaged on a Zeiss LSM 780 confocal microscope using the 10X objective and 405 laser; hatchling cuticle is highly autofluorescent in the DAPI channel. Images were processed using FIJI. First, the Gamma value was adjusted to create a more visually uniform cuticle, and levels were adjusted to bring the background fluorescence to 0. A depth projection was then generated using the Stacks > Hyperstacks > Temporal Color Code command and the “Spectrum” LUT. Images were adjusted for additional brightness using the Levels command in Adobe Photoshop Creative Cloud.

### *In situ* HCR probes

We ordered *in situ* hybridization chain reaction probes for *Ph-Ubx* isoform 2, *Ph-abd-A* isoform 2, *Ph-Abd-B* isoform 2, *Ph-Scr*, and *Ph-elav* through Molecular Instruments (Lot #PRG221-225).

### *In situ* hybridization chain reaction

We performed *in situ* hybridization chain reaction on dissected embryos using the Bruce et al. modifications to the Choi et al. *in situ* hybridization chain reaction 3.0 method. Embryos were mounted as previously described and imaged using a Zeiss LSM 780 confocal microscope using a 10X or 20X objective. We substantially narrowed the detection range for the fluorophores for each probe to avoid bleedthrough between channels, using wildtype single-probe stains as a benchmark for expected expression boundaries. We utilized the same detection ranges for imaging our mutant and wildtype embryos. Z-stack maximum intensity projection images were generated using FIJI, and background (non-embryo tissue) fluorescence was adjusted to 0 using the Window/Level tool.

### *Hox* expression analysis

For each embryo used for analysis in Fig. 7, we examined the expression of each *Hox* gene compared to the DAPI-stained morphology of each embryo. For each embryo, we evaluated whether any expression was observed in the medial region or the limbs. We assigned a 1 to expression equivalent to the highest wildtype expression (evaluated by eye), and a 0.5 to lower levels of expression or patchy expression. As embryos are symmetrical, but we observed mosaicism in our knockouts, if either half of the embryo showed any expression, we used that value as our measure for that region of the embryo. This allowed us to gauge the maximum possible extent of misexpression in mutants.

## Acknowledgments

We would like to thank members of the Nipam Patel, Hernan Garcia, David Weisblat, and Iswar Hariharan labs for feedback on the experiments and analyses performed for this project. RJC and YT were supervised by Hernan Garcia during their undergraduate thesis work that contributed to this project.

## Contributions

DAS, YT, and RJC conceived of the project. YT, RJC, and DAS performed literature research to identify candidate PcG/TrxG genes. YT compiled literature summary tables. YT, DAS, and RJC performed bioinformatic analyses to identify PcG/TrxG genes, and DAS performed bioinformatic analyses to evaluate gene expression levels. DAS and RJC performed CRISPR injections. RJC, YT, and DAS raised and phenotyped mutant hatchlings. DAS and YT performed confocal microscopy on hatchlings. YT wrote scripts to visualize phenotype data with input from DAS. DAS performed embryo dissections and *in situ* hybridization and antibody stains. DAS wrote and revised the manuscript with input from YT, RJC, and NHP. NHP supervised project.

## References

1. Akam, M. (1987). The molecular basis for metameric pattern in the Drosophila embryo. Development 101, 1–22.

2. Akasaka, T., van Lohuizen, M., van der Lugt, N., Mizutani-Koseki, Y., Kanno, M., Taniguchi, M., Vidal, M., Alkema, M., Berns, A., and Koseki, H. (2001). Mice doubly deficient for the Polycomb Group genes Mel18 and Bmi1 reveal synergy and requirement for maintenance but not initiation of Hox gene expression. Development 128, 1587–1597.

3. Almazán, A., Çevrim, Ç., Musser, J.M., Averof, M., and Paris, M. (2021). Regenerated crustacean limbs are precise replicas. BioRxiv 2021.12.13.472338.

4. Angelini, D.R., and Kaufman, T.C. (2005). Comparative Developmental Genetics and the Evolution of Arthropod Body Plans. Annu. Rev. Genet. 39, 95–119.

5. Averof, M., and Akam, M. (1995). Hox genes and the diversification of insect and crustacean body plans. Nature 376, 420–423.

6. Averof, M., and Patel, N.H. (1997). Crustacean appendage evolution associated with changes in Hox gene expression. Nature 388, 682.

7. Breen, T.R., and Duncan, I.M. (1986). Maternal expression of genes that regulate the bithorax complex of Drosophila melanogaster. Dev. Biol. 118, 442–456.

8. Breen, T.R., and Harte, P.J. (1991). Molecular characterization of the trithorax gene, a positive regulator of homeotic gene expression in Drosophila. Mech. Dev. 35, 113–127.

9. Breen, T.R., and Harte, P.J. (1993). Trithorax regulates multiple homeotic genes in the bithorax and Antennapedia complexes and exerts different tissue-specific, parasegment-specific and promoter-specific effects on each. Development 117, 119–134.

10. Brinkmeier, M.L., Geister, K.A., Jones, M., Waqas, M., Maillard, I., and Camper, S.A. (2015). The Histone Methyltransferase Gene Absent, Small, or Homeotic Discs-1 Like Is Required for Normal Hox Gene Expression and Fertility in Mice1. Biol. Reprod. 93, 121, 1–12.

11. Brown, J.L., Fritsch, C., Mueller, J., and Kassis, J.A. (2003). The Drosophila pho-like gene encodes a YY1-related DNA binding protein that is redundant with pleiohomeotic in homeotic gene silencing. Development 130, 285–294.

12. Bruce, H.S., and Patel, N.H. (2020). Knockout of crustacean leg patterning genes suggests that insect wings and body walls evolved from ancient leg segments. Nat. Ecol. Evol. 4, 1703–1712.

13. Bruce, H., Jerz, G., Kelly, S., McCarthy, J., Pomerantz, A., Senevirathne, G., Sherrard, A., Sun, D., Wolff, C., and Patel, N. (2021). Hybridization Chain Reaction (HCR) In Situ Protocol. Protocols.Io.

14. Brunk, B.P., Martin, E.C., and Adler, P.N. (1991). Molecular genetics of the Posterior sex combs/Suppressor 2 of zeste region of Drosophila: aberrant expression of the Suppressor 2 of zeste gene results in abnormal bristle development. Genetics 128, 119–132.

15. Chambeyron, S., and Bickmore, W.A. (2004). Chromatin decondensation and nuclear reorganization of the HoxB locus upon induction of transcription. Genes Dev. 18, 1119–1130.

16. Choi, H.M.T., Schwarzkopf, M., Fornace, M.E., Acharya, A., Artavanis, G., Stegmaier, J., Cunha, A., and Pierce, N.A. (2018). Third-generation in situ hybridization chain reaction: multiplexed, quantitative, sensitive, versatile, robust. Development 145, dev165753.

17. Clark-Hachtel, C.M., and Tomoyasu, Y. (2020). Two sets of candidate crustacean wing homologues and their implication for the origin of insect wings. Nat. Ecol. Evol. 4, 1694–1702.

18. de Graaff Wim, Tomotsune Daihachiro, Oosterveen Tony, Takihara Yoshihiro, Koseki Haruhiko, and Deschamps Jacqueline (2003). Randomly inserted and targeted Hox/reporter fusions transcriptionally silenced in Polycomb mutants. Proc. Natl. Acad. Sci. 100, 13362–13367.

19. Denell, R.E. (1978). Homoeosis in Drosophila. II. A genetic analysis of Polycomb. Genetics 90, 277.

20. Deschamps, J., and van Nes, J. (2005). Developmental regulation of the Hox genes during axial morphogenesis in the mouse. Development 132, 2931–2942.

21. Deschamps, J., and Wijgerde, M. (1993). Two Phases in the Establishment of HOX Expression Domains. Dev. Biol. 156, 473–480.

22. Deutsch, J.S., and Mouchel-Vielh, E. (2003). Hox genes and the crustacean body plan. BioEssays 25, 878–887.

23. Erokhin, M., Georgiev, P., and Chetverina, D. (2018). Drosophila DNA-Binding Proteins in Polycomb Repression. Epigenomes 2.

24. Fauvarque, M.-O., Laurenti, P., Boivin, A., Bloyer, S., Griffin-Shea, R., Bourbon, H.-M., and Dura, J.-M. (2001). Dominant modifiers of the polyhomeotic extra-sex-combs phenotype induced by marked P element insertional mutagenesis in Drosophila. Genet. Res. 78, 137–148.

25. Geisler, S.J., and Paro, R. (2015). Trithorax and Polycomb group-dependent regulation: a tale of opposing activities. Development 142, 2876–2887.

26. Gerberding, M., Browne, W., and Patel, N. (2002). Cell lineage analysis of the amphipod crustacean Parhyale hawaiensis reveals an early restriction of cell fates. Development 129, 5789–5801.

27. Guenther, M., Jenner, R., and Chevalier, B. (2005). Global and Hox-specific roles for the MLL1 methyltransferase. Proc. ….

28. Hrycaj, S.M., and Wellik, D.M. (2016). Hox genes and evolution. F1000Res 5, 859.

29. Hughes, C.L., and Kaufman, T.C. (2002). Hox genes and the evolution of the arthropod body plan. Evol. Dev. 4, 459–499.

30. Irish, V.F., Martinez-Arias, A., and Akam, M. (1989). Spatial regulation of the Antennapedia and Ultrabithorax homeotic genes during Drosophila early development. EMBO J. 8, 1527–1537.

31. Isono Kyo-ichi, Fujimura Yu-ichi, Shinga Jun, Yamaki Makoto, O-Wang Jiyang, Takihara Yoshihiro, Murahashi Yasuaki, Takada Yuki, Mizutani-Koseki Yoko, and Koseki Haruhiko (2005). Mammalian Polyhomeotic Homologues Phc2 and Phc1 Act in Synergy To Mediate Polycomb Repression of Hox Genes. Mol. Cell. Biol. 25, 6694–6706.

32. Jaeger, J. (2011). The gap gene network. Cell. Mol. Life Sci. 68, 243–274.

33. Kassis, J.A., Kennison, J.A., and Tamkun, J.W. (2017). Polycomb and Trithorax Group Genes in Drosophila. Genetics 206, 1699–1725.

34. Kennison, J.A., and Tamkun, J.W. (1988). Dosage-dependent modifiers of polycomb and antennapedia mutations in Drosophila. Proc. Natl. Acad. Sci. 85, 8136–8140.

35. Kmita, M., and Duboule, D. (2003). Organizing axes in time and space; 25 years of colinear tinkering. Science 301, 331–333.

36. Krumlauf, R. (1994). Hox genes in vertebrate development. Cell 78, 191–201.

37. Kurzhals, R.L., Tie, F., Stratton, C.A., and Harte, P.J. (2008). Drosophila ESC-like can substitute for ESC and becomes required for Polycomb silencing if ESC is absent. Dev. Biol. 313, 293–306.

38. Liubicich, D.M., Serano, J.M., Pavlopoulos, A., Kontarakis, Z., Protas, M.E., Kwan, E., Chatterjee, S., Tran, K.D., Averof, M., and Patel, N.H. (2009). Knockdown of Parhyale Ultrabithorax recapitulates evolutionary changes in crustacean appendage morphology. Proc Natl Acad Sci USA 106, 13892–13896.

39. Lo Stanley M., Ahuja Nitin K., and Francis Nicole J. (2009). Polycomb Group Protein Suppressor 2 of Zeste Is a Functional Homolog of Posterior Sex Combs. Mol. Cell. Biol. 29, 515–525.

40. Maeda, R.K., and Karch, F. (2006). The ABC of the BX-C: the bithorax complex explained. Development 133, 1413–1422.

41. Maeda, R.K., and Karch, F. (2015). The open for business model of the bithorax complex in Drosophila. Chromosoma 124, 293–307.

42. Mallo, M., and Alonso, C.R. (2013). The regulation of Hox gene expression during animal development. Development 140, 3951–3963.

43. del Mar Lorente, M., Marcos-Gutierrez, C., Perez, C., Schoorlemmer, J., Ramirez, A., Magin, T., and Vidal, M. (2000). Loss- and gain-of-function mutations show a polycomb group function for Ring1A in mice. Development 127, 5093–5100.

44. Martin, E.C., and Adler, P.N. (1993). The Polycomb group gene Posterior Sex Combs encodes a chromosomal protein. Development 117, 641–655.

45. Martin, A., Serano, J.M., Jarvis, E., Bruce, H.S., Wang, J., Ray, S., Barker, C.A., C O., Liam, and Patel, N.H. (2015). CRISPR/Cas9 Mutagenesis Reveals Versatile Roles of Hox Genes in Crustacean Limb Specification and Evolution. Curr Biol 26, 14–26.

46. Matsuoka, Y., Bando, T., Watanabe, T., Ishimaru, Y., Noji, S., Popadić, A., and Mito, T. (2015). Short germ insects utilize both the ancestral and derived mode of Polycomb group-mediated epigenetic silencing of Hox genes. Biol. Open 4, 702–709.

47. Ohno, K., McCabe, D., Czermin, B., Imhof, A., and Pirrotta, V. (2008). ESC, ESCL and their roles in Polycomb Group mechanisms. Mech. Dev. 125, 527–541.

48. Panganiban, G., Sebring, A., Nagy, L., and Carroll, S. (1995). The Development of Crustacean Limbs and the Evolution of Arthropods. Science 270, 1363–1366.

49. Pavlopoulos, A., Kontarakis, Z., Liubicich, D.M., Serano, J.M., Akam, M., Patel, N.H., and Averof, M. (2009). Probing the evolution of appendage specialization by Hox gene misexpression in an emerging model crustacean. Proc Natl Acad Sci USA 106, 13897–13902.

50. Peifer, M., Karch, F., and Bender, W. (1987). The bithorax complex: control of segmental identity. Genes Dev. 1, 891–898.

51. Price, A.L., Modrell, M.S., Hannibal, R.L., and Patel, N.H. (2010). Mesoderm and ectoderm lineages in the crustacean Parhyale hawaiensis display intra-germ layer compensation. Dev Biol 341, 256–266.

52. Puro, J., and Nygrén, T. (1975). Mode of action of a homoeotic gene in Drosophila melanogaster: Localization and dosage effects of Polycomb. Hereditas 81, 237–247.

53. Rastegar Mojgan, Kobrossy Laila, Kovacs Erzsebet Nagy, Rambaldi Isabel, and Featherstone Mark (2004). Sequential Histone Modifications at Hoxd4 Regulatory Regions Distinguish Anterior from Posterior Embryonic Compartments. Mol. Cell. Biol. 24, 8090–8103.

54. Schmid, B. (2011). Molecular Studies on Head Development of the Amphipod Crustacean Parhyale hawaiensis (Georg-August-University Göttingen).

55. Schuettengruber, B., Bourbon, H.-M., Di Croce, L., and Cavalli, G. (2017). Genome Regulation by Polycomb and Trithorax: 70 Years and Counting. Cell 171, 34–57.

56. Schwentner, M., Combosch, D.J., Pakes Nelson, J., and Giribet, G. (2017). A Phylogenomic Solution to the Origin of Insects by Resolving Crustacean-Hexapod Relationships. Curr. Biol. 27, 1818–1824.e5.

57. Schwentner, M., Richter, S., Rogers, D.C., and Giribet, G. (2018). Tetraconatan phylogeny with special focus on Malacostraca and Branchiopoda: highlighting the strength of taxon-specific matrices in phylogenomics. Proc. R. Soc. B Biol. Sci. 285.

58. Serano, J.M., Martin, A., Liubicich, D.M., Jarvis, E., Bruce, H.S., La, K., Browne, W.E., Grimwood, J., and Patel, N.H. (2016). Comprehensive analysis of Hox gene expression in the amphipod crustacean Parhyale hawaiensis. Dev Biol 409, 297–309.

59. Shiga, Y., Sagawa, K., Takai, R., Sakaguchi, H., Yamagata, H., and Hayashi, S. (2006). Transcriptional readthrough of Hox genes Ubx and Antp and their divergent post-transcriptional control during crustacean evolution. Evol. Dev. 8, 407–414.

60. Simon, J., Chiang, A., and Bender, W. (1992). Ten different Polycomb group genes are required for spatial control of the abdA and AbdB homeotic products. Development 114, 493–505.

61. Soshnikova, N., and Duboule, D. (2009). Epigenetic temporal control of mouse Hox genes in vivo. Science 324, 1320–1323.

62. Struhl, G., and Akam, M. (1985). Altered distributions of Ultrabithorax transcripts in extra sex combs mutant embryos of Drosophila. EMBO J. 4, 3259–3264.

63. Struhl, G., and White, R.A.H. (1985). Regulation of the Ultrabithorax gene of drosophila by other bithorax complex genes. Cell 43, 507–519.

64. Sun, D.A., Bredeson, J.V., Bruce, H.S., and Patel, N.H. (2021). Identification and classification of cis-regulatory elements in the amphipod crustacean *Parhyale hawaiensis*. BioRxiv 2021.09.16.460328.

65. Tschopp, P., and Duboule, D. (2011). A genetic approach to the transcriptional regulation of Hox gene clusters. Annu. Rev. Genet. 45, 145–166.

66. Wang, H., Wang, L., Erdjument-Bromage, H., Vidal, M., Tempst, P., Jones, R.S., and Zhang, Y. (2004). Role of histone H2A ubiquitination in Polycomb silencing. Nature 431, 873–878.

67. Wang, S., He, F., Xiong, W., Gu, S., Liu, H., Zhang, T., Yu, X., and Chen, Y. (2007). Polycomblike-2-deficient mice exhibit normal left–right asymmetry. Dev. Dyn. 236, 853–861.

68. Wedeen, C., Harding, K., and Levine, M. (1986). Spatial regulation of Antennapedia and bithorax gene expression by the Polycomb locus in Drosophila. Cell 44, 739–748.

69. Wolfe, J.M., Breinholt, J.W., Crandall, K.A., Lemmon, A.R., Lemmon, E.M., Timm, L.E., Siddall, M.E., and Bracken-Grissom, H.D. (2019). A phylogenomic framework, evolutionary timeline and genomic resources for comparative studies of decapod crustaceans. Proc. R. Soc. B Biol. Sci. 286, 20190079.

70. Yu Benjamin D., Hanson Robin D., Hess Jay L., Horning Susan E., and Korsmeyer Stanley J. (1998). MLL, a mammalian trithorax-group gene, functions as a transcriptional maintenance factor in morphogenesis. Proc. Natl. Acad. Sci. 95, 10632–10636.

