## Supplemental Figures for "Distinct regulation of *Hox* genes by Polycomb Group genes in a crustacean"

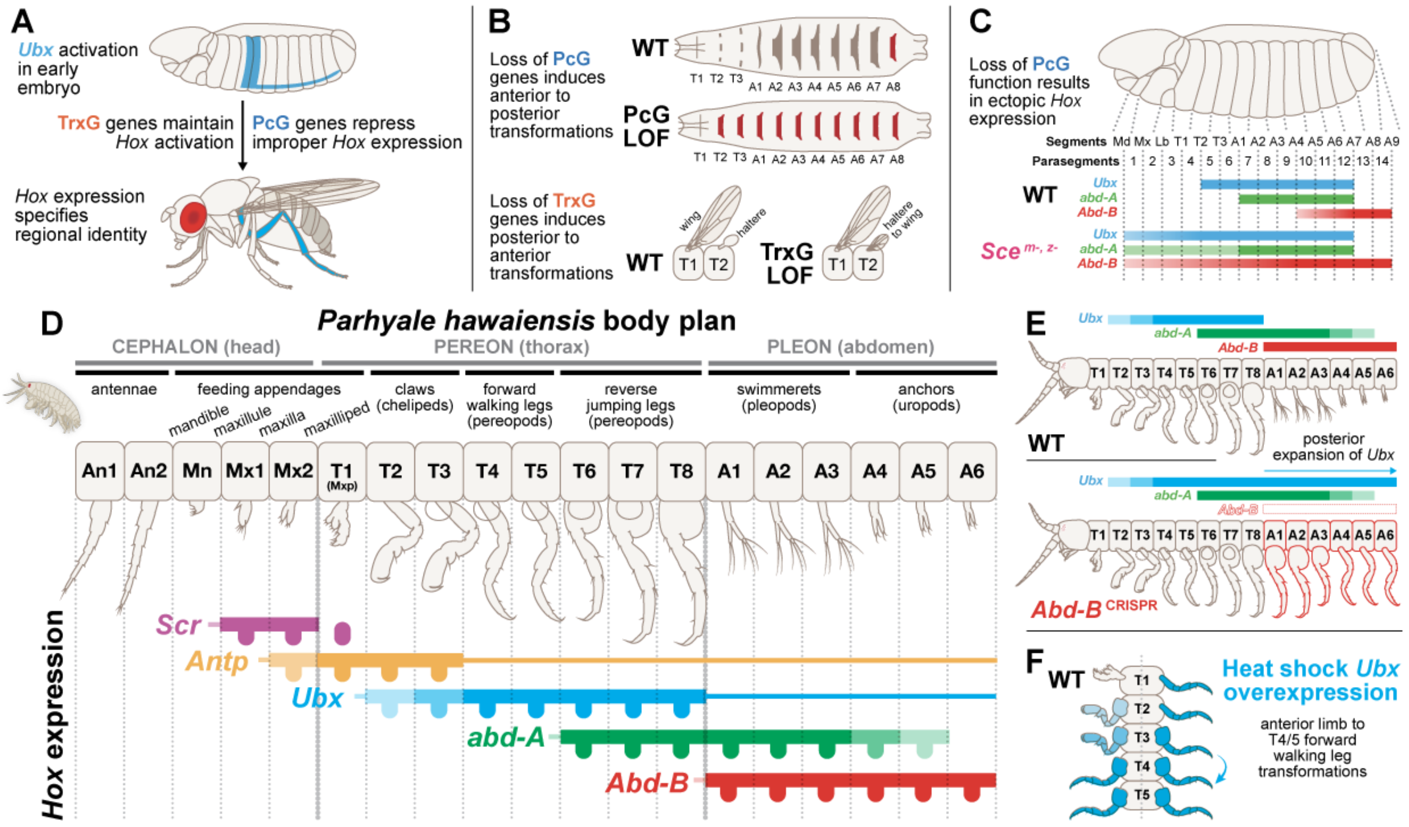


#### Supp. Fig. 1.1: Background

A) Summary model of phases of *Hox* regulation as described in *Drosophila*. *Hox* expression boundaries are specified in the early embryo via transcription factor-based mechanisms. As development progresses, PcG and TrxG genes are required to maintain proper *Hox* expression through repressive and activating mechanisms, respectively. The presence of *Hox* effectors in given body regions throughout development helps specify regional identity.

B) Summary of PcG and TrxG loss of function phenotypes. Loss of PcG genes frequently results in all body segments transformed to the most posterior body segment. Loss of TrxG genes can induce posterior to anterior transformations, such as haltere to wing transformation.

C) Example of *Hox* misexpression in PcG mutants. Loss of function of *Sce* results in broad misexpression of multiple *Hox* genes, with *Abd-B* overriding the other *Hox* identities to cause transformations of all thoracic segments to the A8 identity.

D) Limb identities and expression of five *Hox* genes (*Scr*, *Antp*, *Ubx*, *abd-A*, and *Abd-B*) in the wildtype *Parhyale* body plan.

E) Summary of *Abd-B* CRISPR knockout experiments from M(Martin et al., 2015) and Jarvis et al., 2022. Loss of *Abd-B* results in expansion of *Ubx* to the posterior of the embryo, and subsequent homeotic transformations. This result indicates that *Abd-B* represses *Ubx* to set the posterior boundary of *Ubx* expression.

F) Summary of *Ubx* heat shock overexpression results from Pavlopoulos et al., 2009. Heat shock results in transformation of anterior appendages to T4/5 forward walking leg identity.

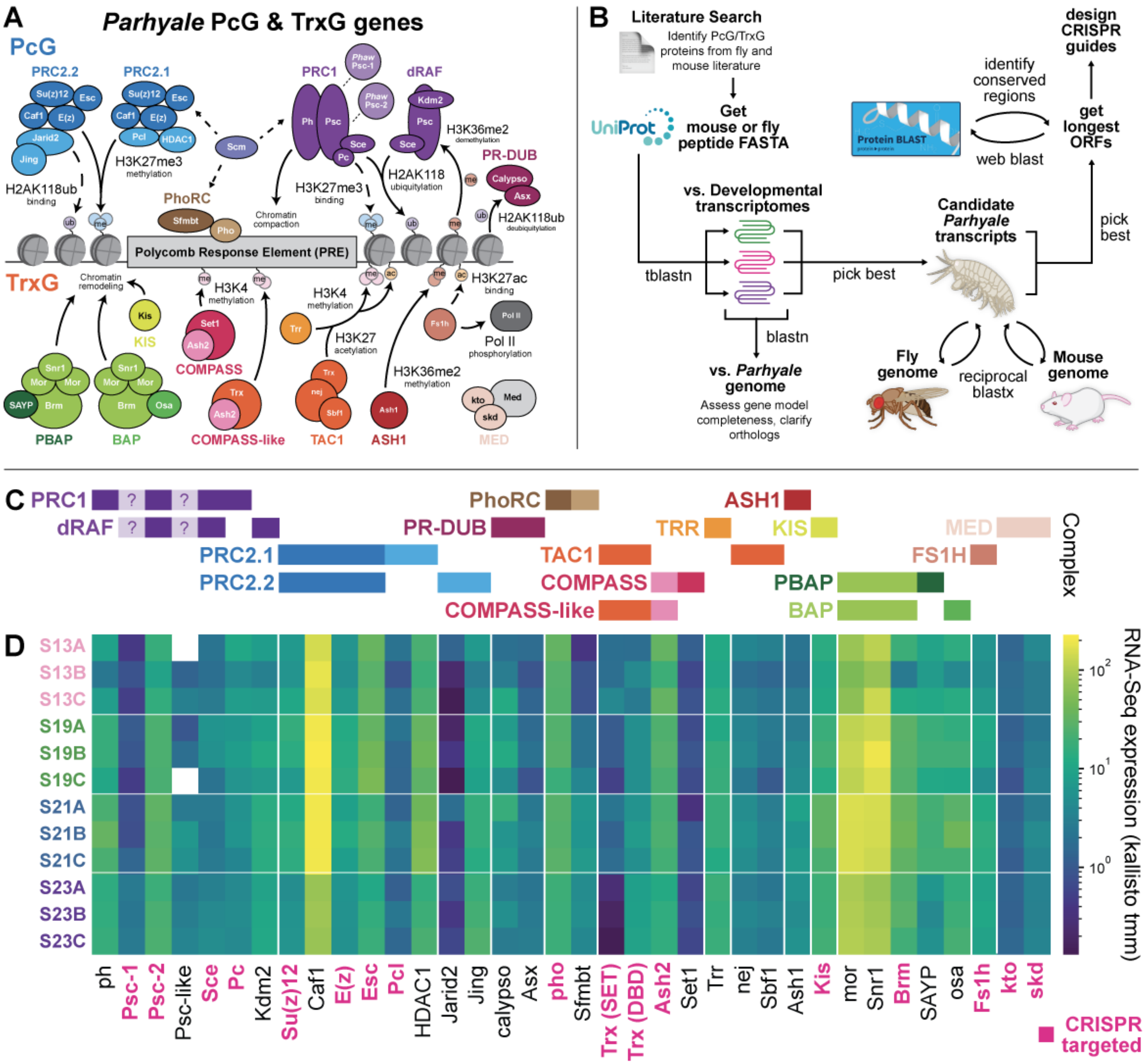


#### Supp. Fig. 1.2: Identification and expression of PcG/TrxG genes in *Parhyale hawaiensis*

A) Full diagram of PcG/TrxG genes found in the *Parhyale* genome.

B) Schematic of bioinformatic methods used to identify PcG/TrxG genes. A literature review was performed to identify the core components of PcG/TrxG genes in both *Mus musculus* and *Drosophila melanogaster*. Mouse and fly peptide sequences were obtained for each gene from UNIPROT and tblastn was used to identify the top BLAST hits in several transcript sources, including the phaw_5.0 MAKER genome annotation, the Kao et al. Mikado annotation, the Trinity-limb transcriptome from Sun et al., and the Sun et al. Mikado annotation. For each top candidate, the gene models were manually examined within the *Parhyale* genome to identify gene model completeness and clarify orthologs based on relative genomic location. We also identified the location of possible splice junctions at this step to avoid designing CRISPR guides that overlapped with slice junctions. After determining the best candidate locus and best gene model from across the transcript sources, we performed reciprocal BLAST to both the *Drosophila* and mouse genomes to confirm our top hits, and clarify potential paralog identities. Upon identifying the best candidate for each gene and paralog in the genome, we used online web BLAST to identify potential conserved protein domains and peptide sequences in order to design CRISPR guide RNAs.

C) Summary of PcG and TrxG complexes in the *Parhyale* genome.

D) Expression of each PcG/TrxG gene in the *Parhyale* genome across three replicates at four developmental stages (S13, S19, S21, S23). Transcriptome data from Sun et al. 2022 was used. All genes of interest showed expression at all timepoints.

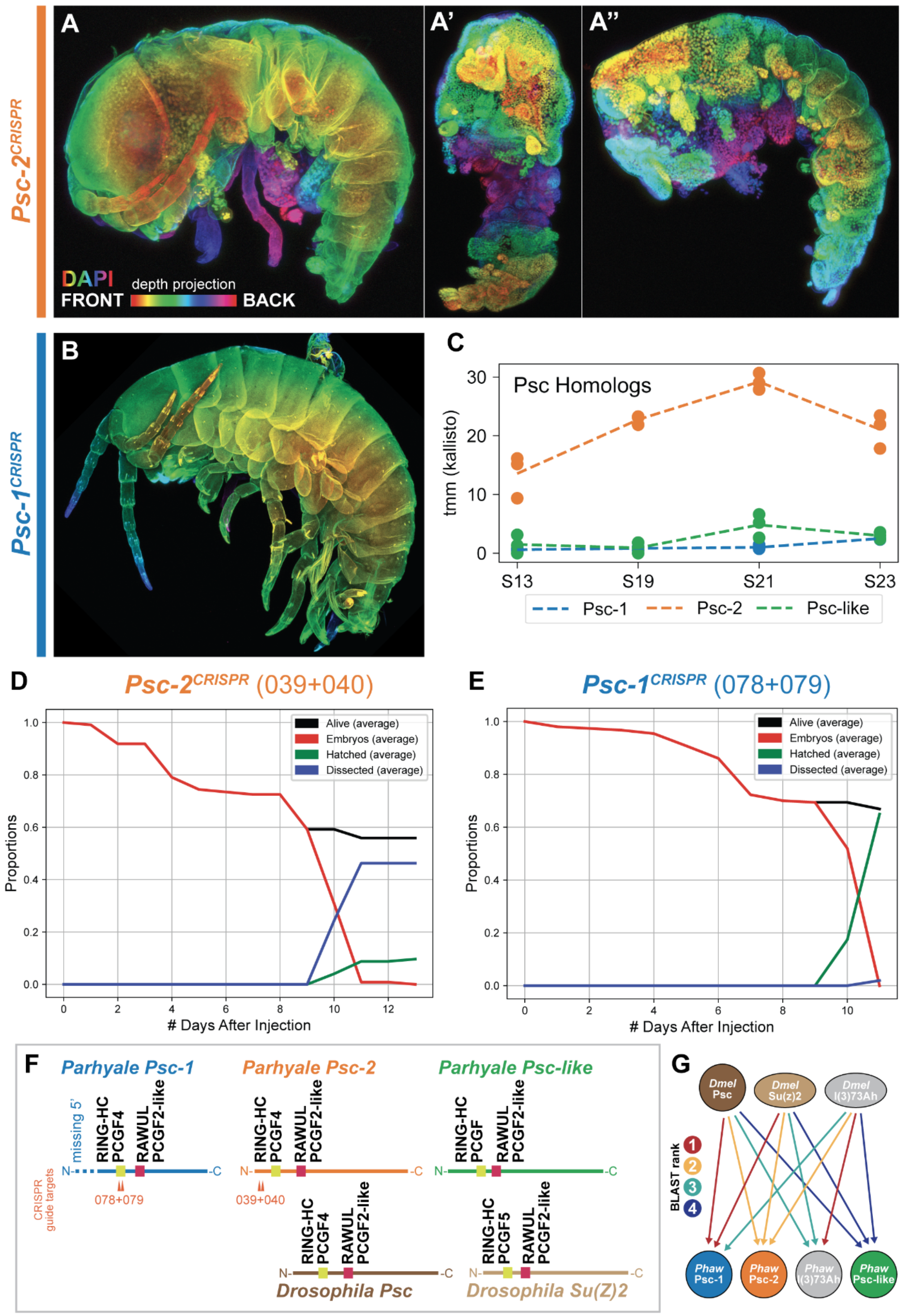

Psc-2CRISPR
A
A'
A''
DAPI
depth projection
FRONT
BACK
Psc-1CRISPR
B
C
Psc Homologs
tmm (kallisto)
S13
S19
S21
S23
Psc-1
Psc-2
Psc-like
D
Psc-2CRISPR (039+040)
Alive (average)
Embryos (average)
Hatched (average)
Dissected (average)
Proportions
### Days After Injection
E
Psc-1CRISPR (078+079)
F
Parhyale Psc-1
Parhyale Psc-2
Parhyale Psc-like
missing 5'
RING-HC
PCGF4
RAWUL
PCGF2-like
PCGF
PCGF5
CRISPR guide targets
078+079
039+040
Drosophila Psc
Drosophila Su(Z)2
G
Dmel Psc
Dmel Su(z)2
Dmel l(3)73Ah
BLAST rank
Phaw Psc-1
Phaw Psc-2
Phaw l(3)73Ah
Phaw Psc-like

#### Supp. Fig. 2.1: *Psc* family genes in *Parhyale*

A) *Psc-2* CRISPR knockout hatchlings. Hatchlings showed major morphological defects, including total loss of limbs and plates, with varying degrees of severity. A shows an apparently mosaic knockout, where some limbs and other body structures are visible. A' shows a severe phenotype from a ventral view, showing major reduction along the proximal-distal axis. A'' shows a severe phenotype from a lateral view, showing that body segments develop, but limbs do not.

*B)* *Psc-1* CRISPR knockout hatchling. We did not observe any noticeable phenotypes from dual-guide knockout of *Psc-1*.

C) Expression of *Psc-1*, *Psc-2*, and *Psc-like* in the Sun et al. 2022 embryonic transcriptome dataset. *Psc-2* is expressed at a much higher level than *Psc-1* or *Psc-like*.

D) Average survivorship curve for *Psc-2* CRISPR experiments. The vast majority of surviving *Psc-2* hatchlings required dissection, and were not able to hatch on their own.

E) Average survivorship curve for *Psc-1* CRISPR experiments. Nearly all embryos surviving to late stages of development were able to hatch successfully.

F) Diagram of *Parhyale Psc-1*, *Psc-2*, and *Psc-like* genes compared to *Drosophila Psc* and *Su(Z)2* genes. BLAST identified homology domains are placed in relative position along the peptide. All three *Parhyale Psc* homologs appear to have RING-HC PCGF domains and RAWUL PCGF2-like domains.

G) BLAST top hit mapping for *Drosophila* to *Parhyale* genes. *Parhyale Psc-1* is the top hit for both *Drosophila Psc* and *Su(Z)2*, while *Parhyale Psc-2* is the second best BLAST hit for *Drosophila Psc* and *Su(Z)2*. The third top BLAST hit for both *Drosophila Psc* and *Su(Z)2* is *Parhyale l(3)73Ah*, which was identified as being the top BLAST hit for *Drosophila l(3)73Ah*. A fourth gene, which we named *Parhyale Psc-like*, was a top blast hit for *Drosophila Psc*, *Su(Z)2*, and *l(3)73Ah*, but the reciprocal best BLAST hit for *Parhyale Psc-like* was *Drosophila Psc*. Given that *Parhyale Psc-1* and *Psc-2* are better reciprocal BLAST hits for *Drosophila Psc*, we termed this fourth gene *Parhyale Psc-like*.

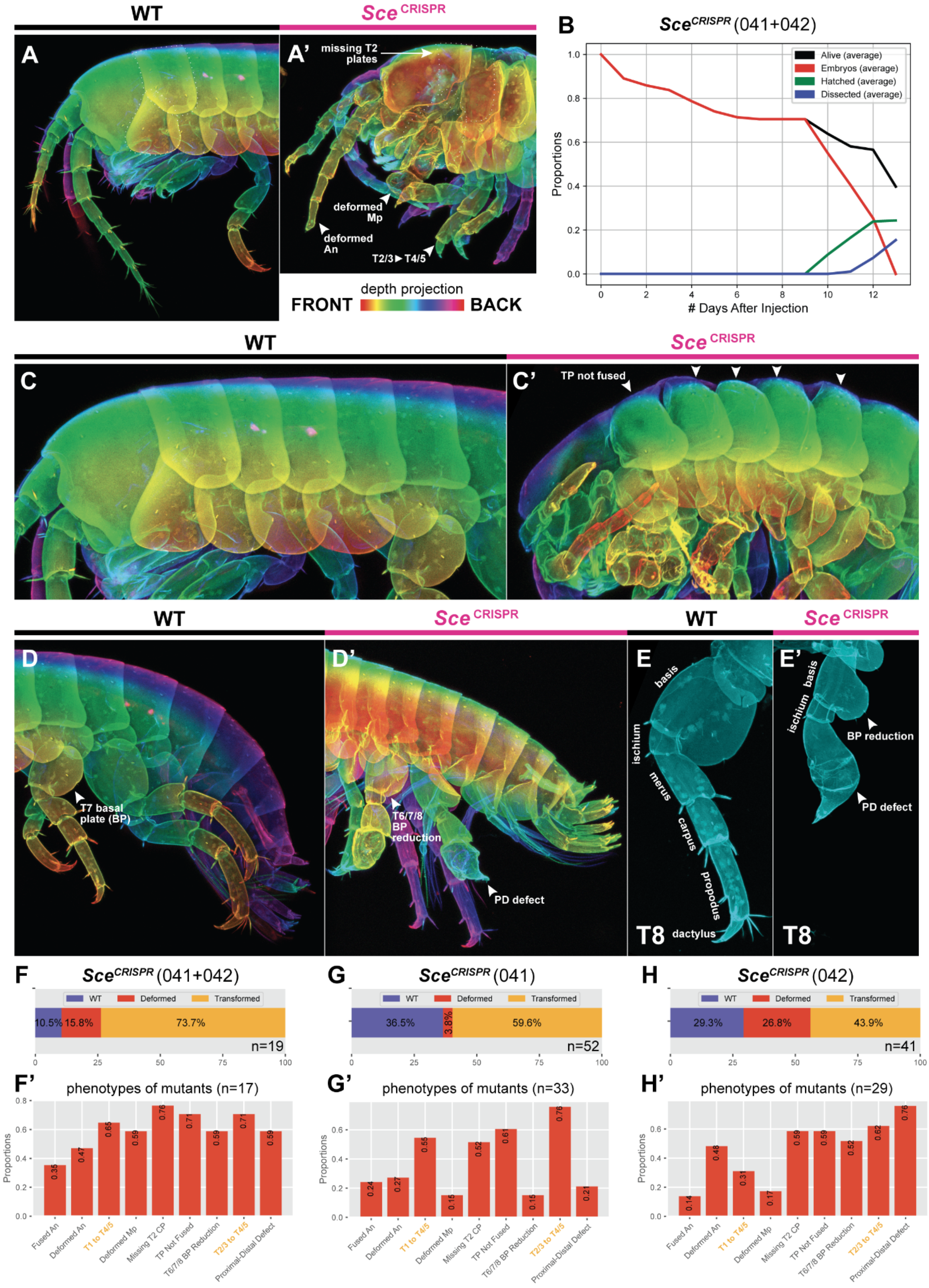

WT
Sce CRISPR
A
A'
missing T2 plates
deformed Mp
deformed An
T2/3 ► T4/5
depth projection
FRONT
BACK
B
Sce CRISPR (041+042)
Alive (average)
Embryos (average)
Hatched (average)
Dissected (average)
Proportions
### Days After Injection
C
C'
TP not fused
D
D'
T7 basal plate (BP)
T6/7/8 BP reduction
PD defect
E
E'
basis
ischium
merus
carpus
propodus
dactylus
T8
BP reduction
PD defect
F
Sce CRISPR (041+042)
WT
Deformed
Transformed
10.5%
15.8%
73.7%
n=19
G
Sce CRISPR (041)
36.5%
3.8%
59.6%
n=52
H
Sce CRISPR (042)
29.3%
26.8%
43.9%
n=41
F'
phenotypes of mutants (n=17)
G'
phenotypes of mutants (n=33)
H'
phenotypes of mutants (n=29)
Fused An
Deformed An
T1 to T4/5
Deformed Mp
Missing T2 CP
TP Not Fused
T6/7/8 BP Reduction
T2/3 to T4/5
Proximal-Distal Defect

#### Supp. Fig. 2.2: *Sce* CRISPR phenotype data

A) Wildtype *Parhyale* hatchling head and anterior thorax.

A') *Sce* CRISPR hatchling head and anterior thorax showing various homeotic transformations and missing T2 tergal and coxal plates.

B) Average survivorship curve for dual-guide *Sce* CRISPR knockout experiments.

C) Wildtype *Parhyale* hatchling view focused on tergal and coxal plates.

C') *Sce* CRISPR hatchling showing defects in tergal plate fusion marked with arrowheads.

D) Wildtype *Parhyale* hatchling posterior, with T8 tergal plate marked with arrowhead.

D') *Sce* CRISPR hatchling showing tergal plate reduction and proximal-distal defects, marked with arrowheads.

E) Dissected wildtype T8 limb with leg segments labeled.

E') Dissected *Sce* CRISPR T8 limb with leg segments and limb defects labeled.

F) Proportion of WT, deformed, and transformed hatchlings for *Sce* dual-guide knockout (guide RNAs 041 and 042) from n=19 hatchlings.

F') Proportion of phenotypes from *Sce* (041+042) dual-guide knockout for n=17 hatchlings that we did not classify as wildtype. Yellow labels indicate phenotypes we classified as homeotic mutations.

G) Proportion of WT, deformed, and transformed hatchlings for *Sce* (041) single-guide knockout for n=52 hatchlings.

G') Proportion of phenotypes from *Sce* (041) single-guide knockout for n=33 mutant hatchlings.

H) Proportion of WT, deformed, and transformed hatchlings for *Sce* (042) single-guide knockout for n=41 hatchlings.

H') Proportion of phenotypes from *Sce* (042) single-guide knockout for n=29 mutant hatchlings.

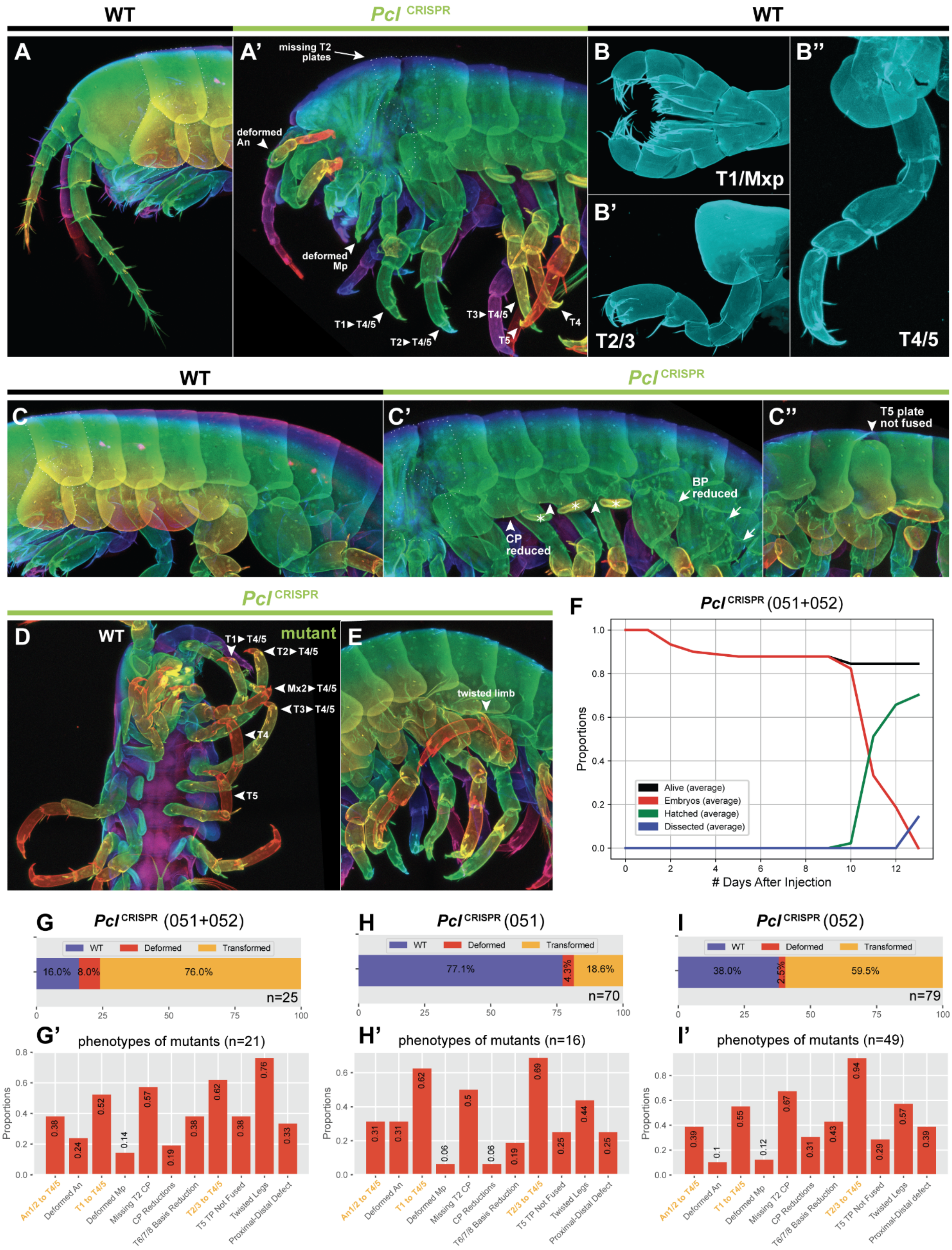

WT
Pcl CRISPR
WT
A
A'
missing T2 plates
deformed An
deformed Mp
T1►T4/5
T2►T4/5
T3►T4/5
T5
T4
B
T1/Mxp
B'
T2/3
B''
T4/5
WT
Pcl CRISPR
C
C'
CP reduced
BP reduced
C''
T5 plate not fused
Pcl CRISPR
D
WT
mutant
T1►T4/5
T2►T4/5
Mx2►T4/5
T3►T4/5
T4
T5
E
twisted limb
F
Pcl CRISPR (051+052)
Proportions
### Days After Injection
Alive (average)
Embryos (average)
Hatched (average)
Dissected (average)
G
Pcl CRISPR (051+052)
WT
Deformed
Transformed
16.0%
8.0%
76.0%
n=25
H
Pcl CRISPR (051)
77.1%
4.3%
18.6%
n=70
I
Pcl CRISPR (052)
38.0%
2.5%
59.5%
n=79
G'
phenotypes of mutants (n=21)
0.38
0.24
0.52
0.14
0.57
0.19
0.38
0.62
0.38
0.76
0.33
H'
phenotypes of mutants (n=16)
0.31
0.31
0.62
0.06
0.5
0.06
0.19
0.69
0.25
0.44
0.25
I'
phenotypes of mutants (n=49)
0.39
0.1
0.55
0.12
0.67
0.31
0.43
0.94
0.29
0.57
0.39
An1/2 to T4/5
Deformed An
T1 to T4/5
Deformed Mp
Missing T2 CP
CP Reductions
T6/7/8 Basis Reduction
T2/3 to T4/5
T5 TP Not Fused
Twisted Legs
Proximal-Distal Defect

#### Supp. Fig. 2.3: *Pcl* CRISPR phenotype data

A) Wildtype *Parhyale* hatchling head and anterior thorax.
A') *Pcl* CRISPR hatchling head and anterior thorax showing various homeotic transformations and missing T2 tergal and coxal plates.
B) Wildtype T1 appendages.
B') Wildtype T2/3 appendage.
B'') Wildtype T4/5 appendage.
C) Wildtype *Parhyale* hatchling view focused on tergal and coxal plates.
C') *Pcl* CRISPR hatchling showing reduction in basal plates (arrows) and exposed gills (asterisks) revealed by reduction in coxal plates.
C'') *Pcl* CRISPR hatchling showing tergal plate fusion defect.
D) *Pcl* CRISPR hatchling showing unilateral transformations from a ventral view. Mx2 - T3 appendages are all transformed towards T4/5 forward walking leg identity.
E) *Pcl* CRISPR hatchling showing twisted legs.
F) Average survivorship curve of *Pcl* dual-guide knockout experiments.
G) Proportion of WT, deformed, and transformed hatchlings for *Pcl* dual-guide knockout (guide RNAs 051 and 052) from n=25 hatchlings.
G') Proportion of phenotypes from *Pcl* (051+052) dual-guide knockout for n=21 hatchlings that we did not classify as wildtype. Yellow labels indicate phenotypes we classified as homeotic mutations.
H) Proportion of WT, deformed, and transformed hatchlings for *Pcl* (051) single-guide knockout for n=70 hatchlings.
H') Proportion of phenotypes from *Pcl* (051) single-guide knockout for n=16 mutant hatchlings.
I) Proportion of WT, deformed, and transformed hatchlings for *Pcl* (052) single-guide knockout for n=79 hatchlings.
I') Proportion of phenotypes from *Pcl* (052) single-guide knockout for n=49 mutant hatchlings.

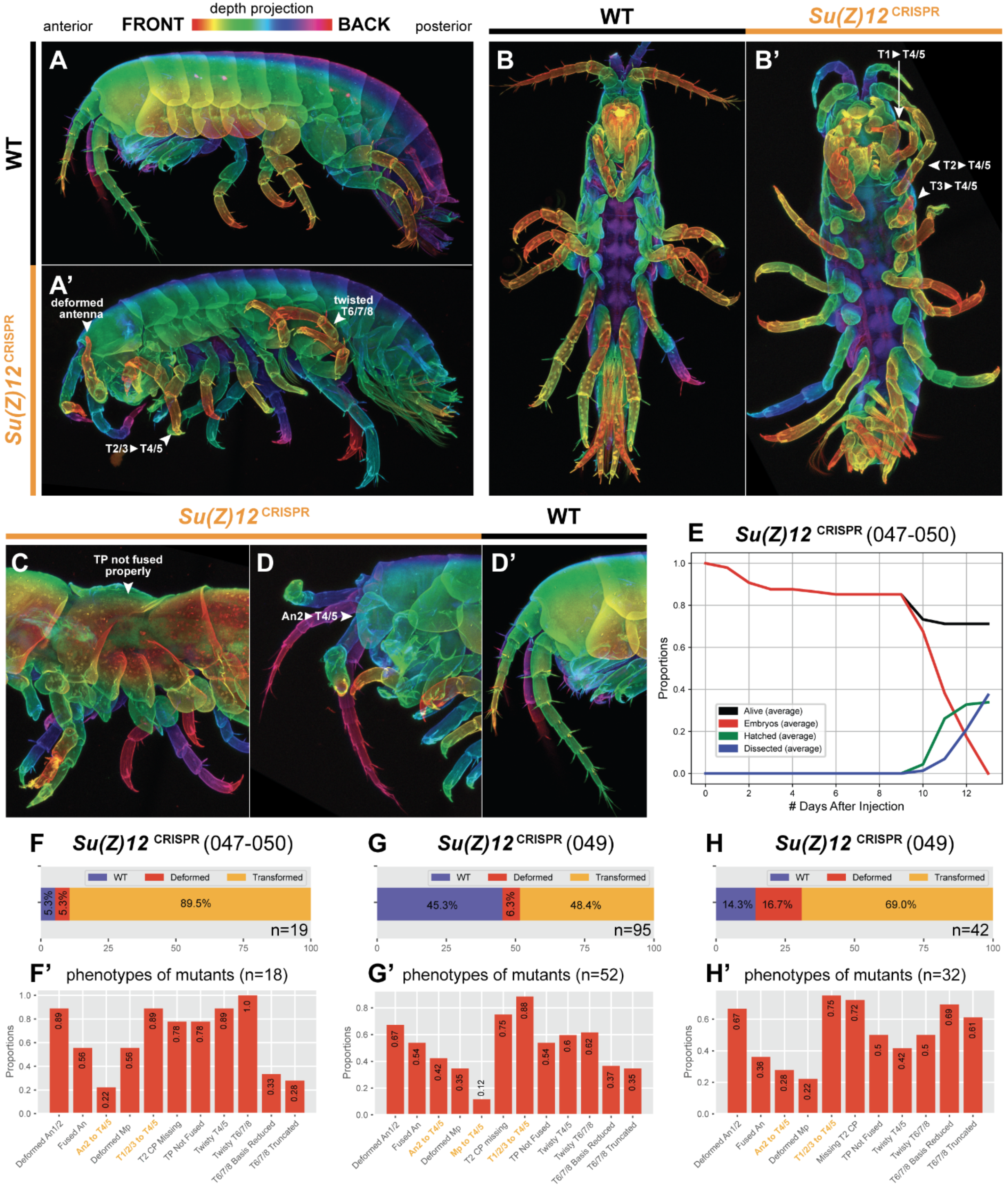
depth projection
anterior
FRONT
BACK
posterior
WT
Su(Z)12 CRISPR
A
A'
deformed antenna
twisted T6/7/8
T2/3► T4/5
B
B'
T1► T4/5
T2► T4/5
T3► T4/5
C
TP not fused properly
D
An2► T4/5
D'
E
Su(Z)12 CRISPR (047-050)
Proportions
### Days After Injection
Alive (average)
Embryos (average)
Hatched (average)
Dissected (average)
F
Su(Z)12 CRISPR (047-050)
WT
Deformed
Transformed
5.3%
5.3%
89.5%
n=19
G
Su(Z)12 CRISPR (049)
45.3%
6.3%
48.4%
n=95
H
Su(Z)12 CRISPR (049)
14.3%
16.7%
69.0%
n=42
F' phenotypes of mutants (n=18)
G' phenotypes of mutants (n=52)
H' phenotypes of mutants (n=32)
Deformed An1/2
Fused An
An2 to T4/5
Deformed Mp
Mp to T4/5
T1/2/3 to T4/5
T2 CP Missing
Missing T2 CP
TP Not Fused
Twisty T4/5
Twisty T6/7/8
T6/7/8 Basis Reduced
T6/7/8 Truncated
0.89
0.56
0.22
0.56
0.89
0.78
0.78
0.89
1.0
0.33
0.28
0.67
0.54
0.42
0.35
0.12
0.75
0.88
0.54
0.6
0.62
0.37
0.35
0.67
0.36
0.28
0.22
0.75
0.72
0.5
0.42
0.5
0.69
0.61

#### Supp. Fig. 2.4: *Su(Z)12* CRISPR phenotype data

A) Wildtype *Parhyale* hatchling, lateral view.
A') *Su(Z)12* CRISPR hatchling showing various homeotic transformations and limb defects.
B) Wildtype *Parhyale* hatchling, ventral view.
B') *Su(Z)12* CRISPR hatchling showing various unilateral homeotic transformations and limb defects, ventral view.
C) *Su(Z)12* CRISPR hatchling showing tergal plate fusion defect.
D) *Su(Z)12* CRISPR hatchling head showing antenna to T4/5 partial transformation. The hallmark of this transformation is the emergence of a coxal plate on the antennal segment, not normally observed.
D') Wildtype *Parhyale* hatchling head. Note the absence of the plate seen in the *Su(Z)12* knockout.
E) Average survivorship curve of *Su(Z)12* quadruple-guide knockout experiments.
F) Proportion of WT, deformed, and transformed hatchlings for *Pcl* quadruple-guide knockout (guide RNAs 047-050) from n=19 hatchlings. We also performed single-guide knockout experiments for guides 047 and 048, which we designed as a result of ambiguity about the true START codon for the *Su(Z)12* gene. These guides did not appear to show any obvious defects or homeotic transformations.
F') Proportion of phenotypes from *Su(Z)12* (047-050) quadruple-guide knockout for n=18 hatchlings that we did not classify as wildtype. Yellow labels indicate phenotypes we classified as homeotic mutations.
G) Proportion of WT, deformed, and transformed hatchlings for *Su(Z)12* (049) single-guide knockout for n=95 hatchlings.
G') Proportion of phenotypes from *Su(Z)12* (049) single-guide knockout for n=52 mutant hatchlings.
H) Proportion of WT, deformed, and transformed hatchlings for *Su(Z)12* (050) single-guide knockout for n=42 hatchlings.
H') Proportion of phenotypes from *Su(Z)12* (050) single-guide knockout for n=32 mutant hatchlings.

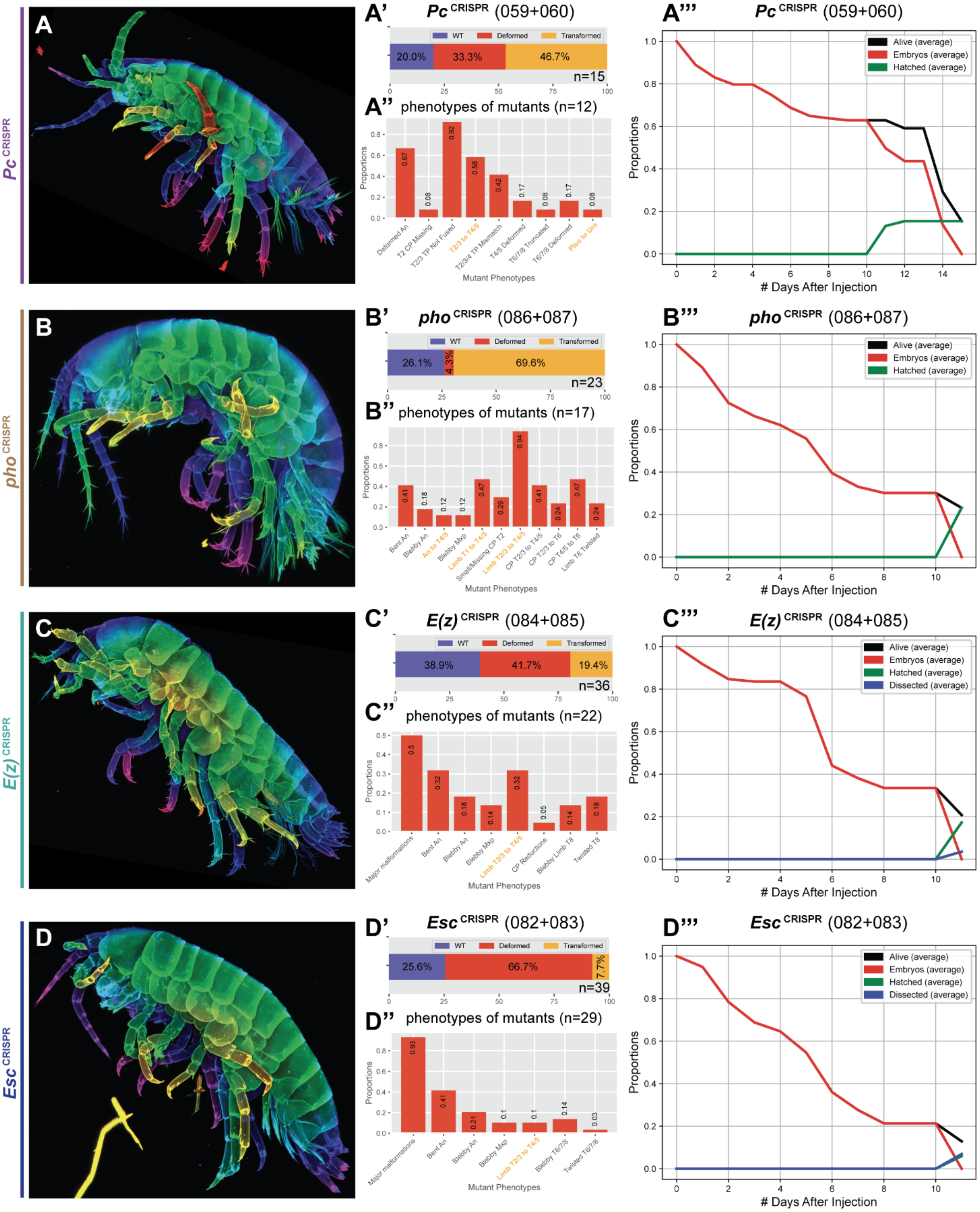

A
Pc CRISPR
A'
Pc CRISPR (059+060)
WT
Deformed
Transformed
20.0%
33.3%
46.7%
n=15
A''
phenotypes of mutants (n=12)
Proportions
Mutant Phenotypes
Deformed An
T2 CP Missing
T2/3 TP Not Fused
T2/3 to T4/5
T2/3/4 TP Mismatch
T4/5 Deformed
T6/7/8 Truncated
T6/7/8 Deformed
Pleo to Uro
0.67
0.08
0.92
0.58
0.42
0.17
0.08
0.17
0.08
A'''
Pc CRISPR (059+060)
Alive (average)
Embryos (average)
Hatched (average)
Proportions
### Days After Injection
B
pho CRISPR
B'
pho CRISPR (086+087)
26.1%
4.3%
69.6%
n=23
B''
phenotypes of mutants (n=17)
Bent An
Blebby An
An to T4/5
Blebby Mxp
Limb T1 to T4/5
Small/Missing CP T2
Limb T2/3 to T4/5
CP T2/3 to T4/5
CP T2/3 to T6
CP T4/5 to T6
Limb T8 Twisted
0.41
0.18
0.12
0.12
0.47
0.29
0.94
0.41
0.24
0.47
0.24
B'''
pho CRISPR (086+087)
C
E(z) CRISPR
C'
E(z) CRISPR (084+085)
38.9%
41.7%
19.4%
n=36
C''
phenotypes of mutants (n=22)
Major malformations
Bent An
Blebby An
Blebby Mxp
Limb T2/3 to T4/5
CP Reductions
Blebby Limb T8
Twisted T8
0.5
0.32
0.18
0.14
0.32
0.05
0.14
0.18
C'''
E(z) CRISPR (084+085)
Dissected (average)
D
Esc CRISPR
D'
Esc CRISPR (082+083)
25.6%
66.7%
7.7%
n=39
D''
phenotypes of mutants (n=29)
Major malformations
Bent An
Blebby An
Blebby Mxp
Limb T2/3 to T4/5
Blebby T6/7/8
Twisted T6/7/8
0.93
0.41
0.21
0.1
0.1
0.14
0.03
D'''
Esc CRISPR (082+083)

#### Supp. Fig. 2.5: Phenotype data for additional PcG genes

A-A''') Representative hatchling showing T2/3 to T4/5 transformations, mutant classes, phenotype proportions, and survivorship curve for *Pc* CRISPR knockout.

B-B''') Representative hatchling showing T2/3 to T4/5 transformations, mutant classes, phenotype proportions, and survivorship curve for *pho* CRISPR knockout.

C-C''') Representative hatchling showing T2/3 to T4/5 transformations, mutant classes, phenotype proportions, and survivorship curve for *E(z)* CRISPR knockout.

D-D''') Representative hatchling showing T2/3 to T4/5 transformations, mutant classes, phenotype proportions, and survivorship curve for *Esc* CRISPR knockout.

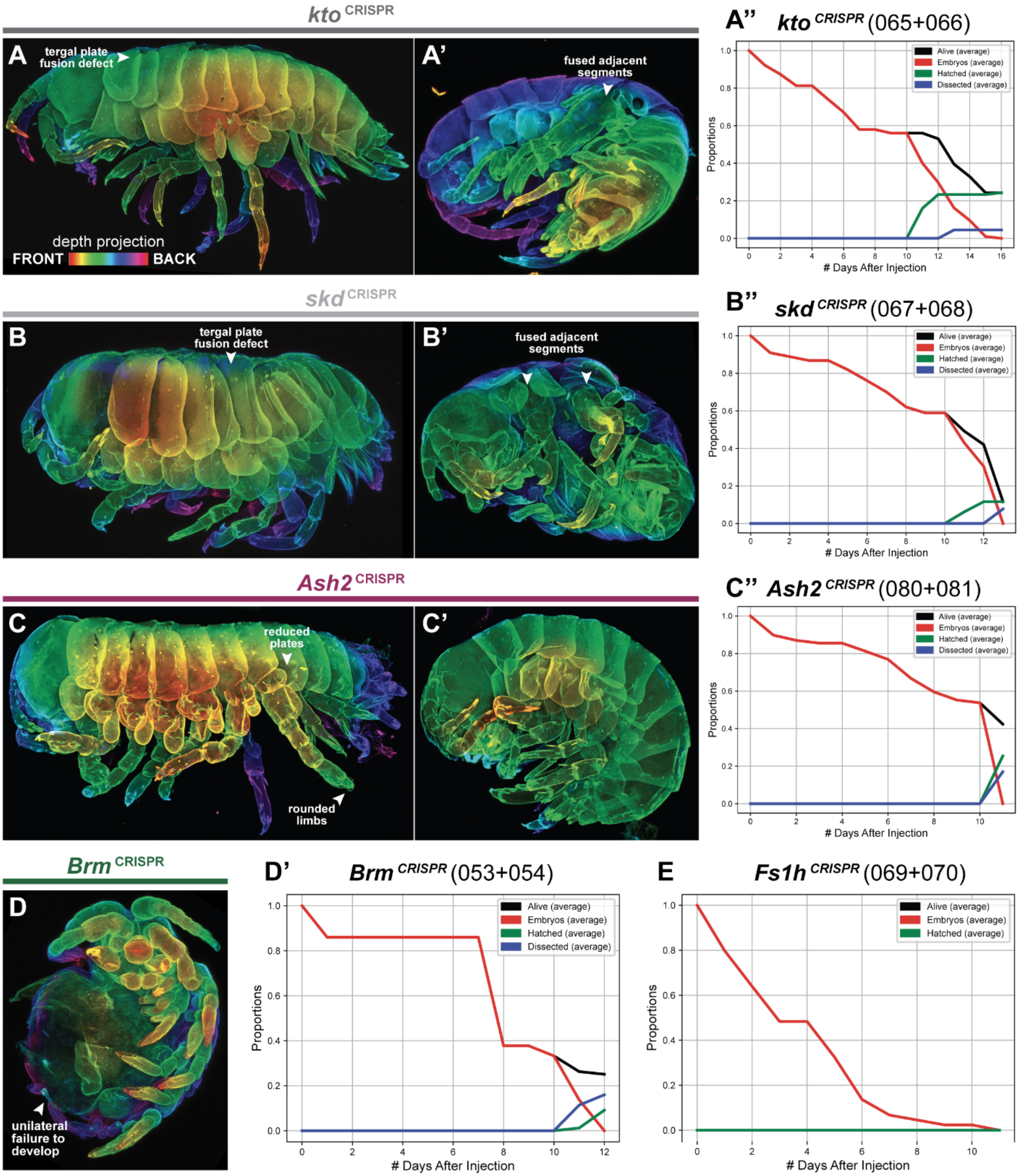

kto CRISPR
A
tergal plate fusion defect
depth projection
FRONT
BACK
A'
fused adjacent segments
A'' kto CRISPR (065+066)
Alive (average)
Embryos (average)
Hatched (average)
Dissected (average)
Proportions
### Days After Injection
skd CRISPR
B
tergal plate fusion defect
B'
fused adjacent segments
B'' skd CRISPR (067+068)
Alive (average)
Embryos (average)
Hatched (average)
Dissected (average)
Proportions
### Days After Injection
Ash2 CRISPR
C
reduced plates
rounded limbs
C'
C'' Ash2 CRISPR (080+081)
Alive (average)
Embryos (average)
Hatched (average)
Dissected (average)
Proportions
### Days After Injection
Brm CRISPR
D
unilateral failure to develop
D' Brm CRISPR (053+054)
Alive (average)
Embryos (average)
Hatched (average)
Dissected (average)
Proportions
### Days After Injection
E Fs1h CRISPR (069+070)
Alive (average)
Embryos (average)
Hatched (average)
Proportions
### Days After Injection

#### Supp. Fig. 2.6: Example phenotypes for TrxG CRISPR knockouts

A-A'') Representative hatchlings and survivorship curve of *kto* CRISPR experiments. A) shows a milder *kto* hatchling, while A') shows a more severe phenotype, with fusion between adjacent body segments marked by arrowhead.

B-B'') Representative hatchlings and survivorship curve of *skd* CRISPR experiments. B) shows a milder *skd* hatchling, while B') shows a more severe phenotype, with fusion between adjacent body segments marked by arrowhead.

C-C'') Representative hatchlings and survivorship curve of *Ash2* CRISPR experiments. C) shows an *Ash2* hatchling with rounded legs and plates, while C') shows a hatchling with minor morphological defects.

D-D') Representative hatchlings and survivorship curve of *Brm* CRISPR experiments. D) shows an example of a *Brm* mutant hatchling where one half of the embryo appears not to have developed.

E) Survivorship curve of *Fs1h* CRISPR experiments. We recovered no surviving embryos from *Fs1h* knockout experiments, suggesting that *Fs1h* knockout is embryonic lethal.

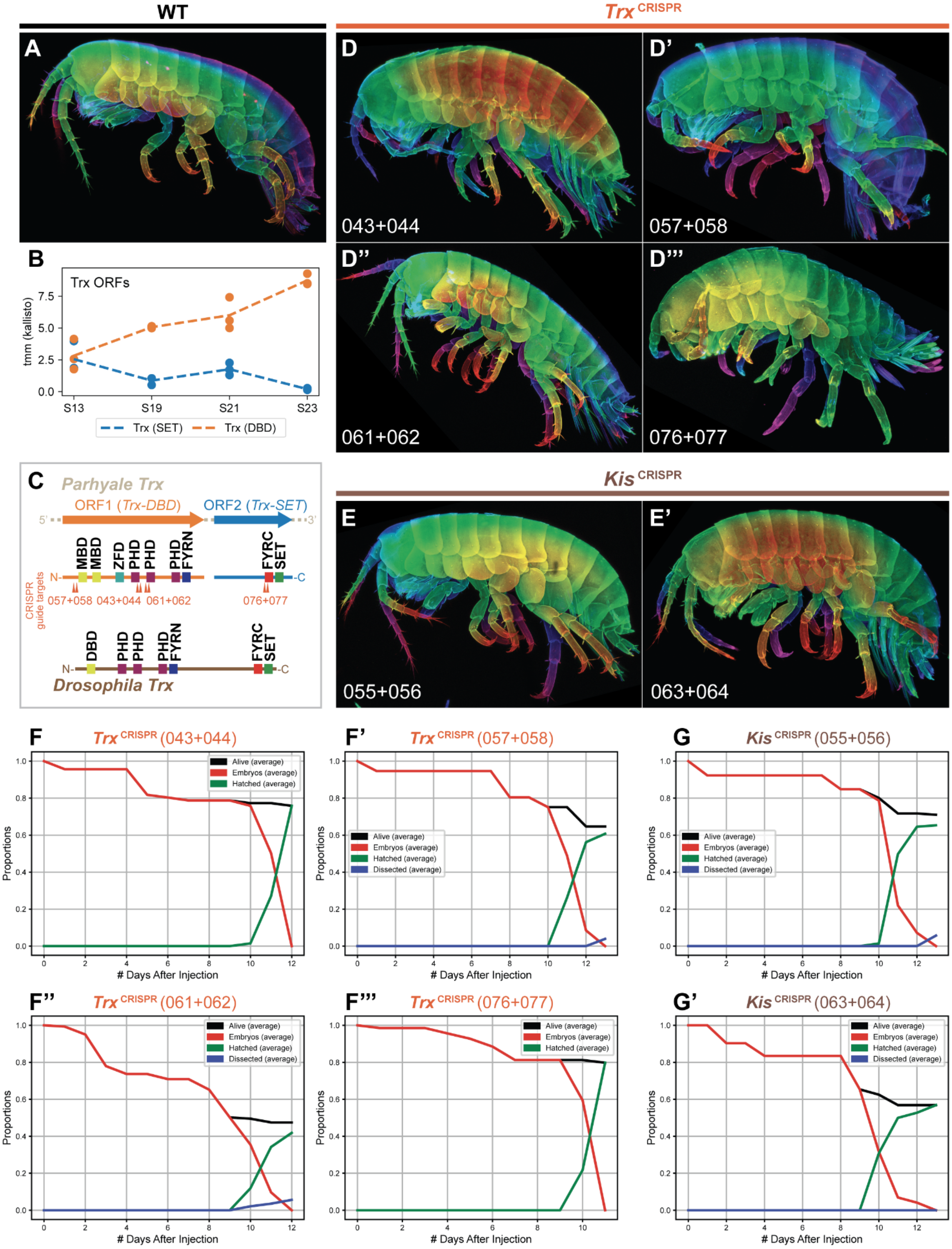

WT
Trx CRISPR
A
D
D'
043+044
057+058
D''
D'''
061+062
076+077
B
Trx ORFs
tmm (kallisto)
S13
S19
S21
S23
Trx (SET)
Trx (DBD)
C
Parhyale Trx
ORF1 (Trx-DBD)
ORF2 (Trx-SET)
MBD
MBD
ZFD
PHD
PHD
PHD
FYRN
FYRC
SET
CRISPR guide targets
057+058
043+044
061+062
076+077
DBD
PHD
PHD
PHD
FYRN
FYRC
SET
Drosophila Trx
Kis CRISPR
E
E'
055+056
063+064
F
Trx CRISPR (043+044)
F'
Trx CRISPR (057+058)
G
Kis CRISPR (055+056)
F''
Trx CRISPR (061+062)
F'''
Trx CRISPR (076+077)
G'
Kis CRISPR (063+064)
Alive (average)
Embryos (average)
Hatched (average)
Dissected (average)
Proportions
### Days After Injection

#### Supp. Fig. 2.7: Lack of obvious phenotypes for *Trx* and *Kis* knockout experiments

A) Wildtype *Parhyale* hatchling, lateral view.

B) Expression of two candidate *Parhyale Trx* ORFs, one containing DNA binding domains (Trx-DBD) and one containing the SET domain (Trx-SET). The two candidate ORFs appear to be expressed at different levels throughout development.

C) Representation of relative position of protein domains comparing *Parhyale Trx-DBD* and *Trx-SET* to *Drosophila Trx*. *Parhyale* may have a split in the two peptide domains. Relative position of CRISPR guide RNAs is illustrated for all 8 guides used to target *Trx*.

D-D''') Representative hatchlings from *Trx* CRISPR knockout experiments. Each experiment used a different pair of guide RNAs, shown in the bottom left corner. We did not observe obvious homeotic transformations for any of the guides.

E-E') Representative hatchlings from *Kis* CRISPR knockout experiments. Each experiment used a different pair of guide RNAs, shown in the bottom left corner. We did not observe obvious homeotic transformations for any of the guides.

F-F''') Survivorship curves for *Trx* CRISPR knockout experiments. We observed varying levels of survivorship for these experiments; however, during the course of the experiments, we did not observe hatchlings that showed obvious morphological defects, as was observed for PcG gene knockout experiments, or other TrxG knockouts, such as *Brm* or *Fs1h*.

G-G') Survivorship curves for *Kis* CRISPR knockout experiments. We observed a similar lack of obvious defects in *Kis* knockout embryos during development.

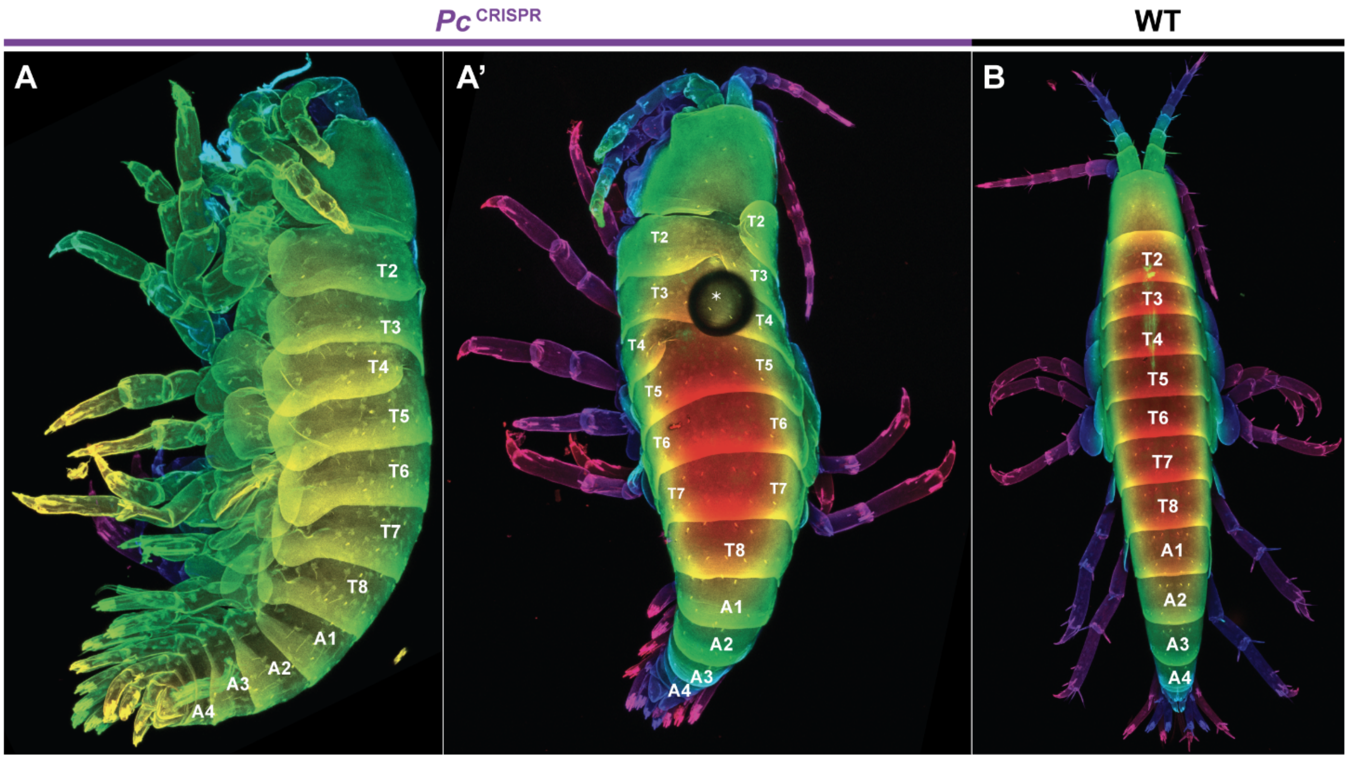


#### Supp. Fig. 3.1: Asymmetric fusion of plates in *Pc* mutant hatchlings

A-A') Lateral and dorsal views of the same *Pc* mutant hatchling, showing asymmetric fusion of plates. The same image is displayed as in Fig. 3; however, we display the entire hatchling here in A' to illustrate how we counted the number of tergal plates.

B) Dorsal view of a WT hatchling showing the counting of tergal plates.

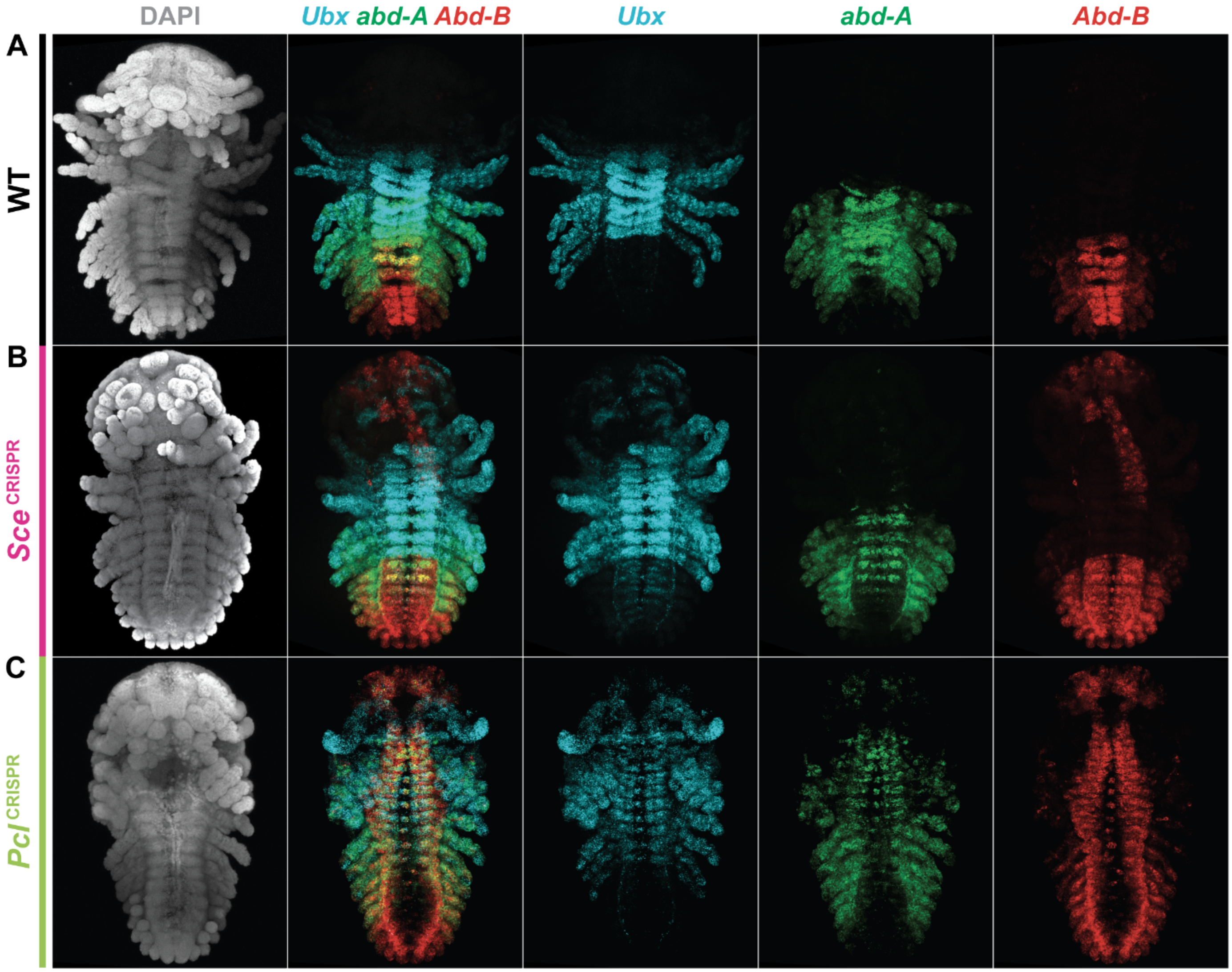


**Supp. Fig. 4.1: Expression of posterior *Hox* genes in PcG mutant embryos**

This figure is identical to Fig. 4, with the exception that we have removed limb labels.

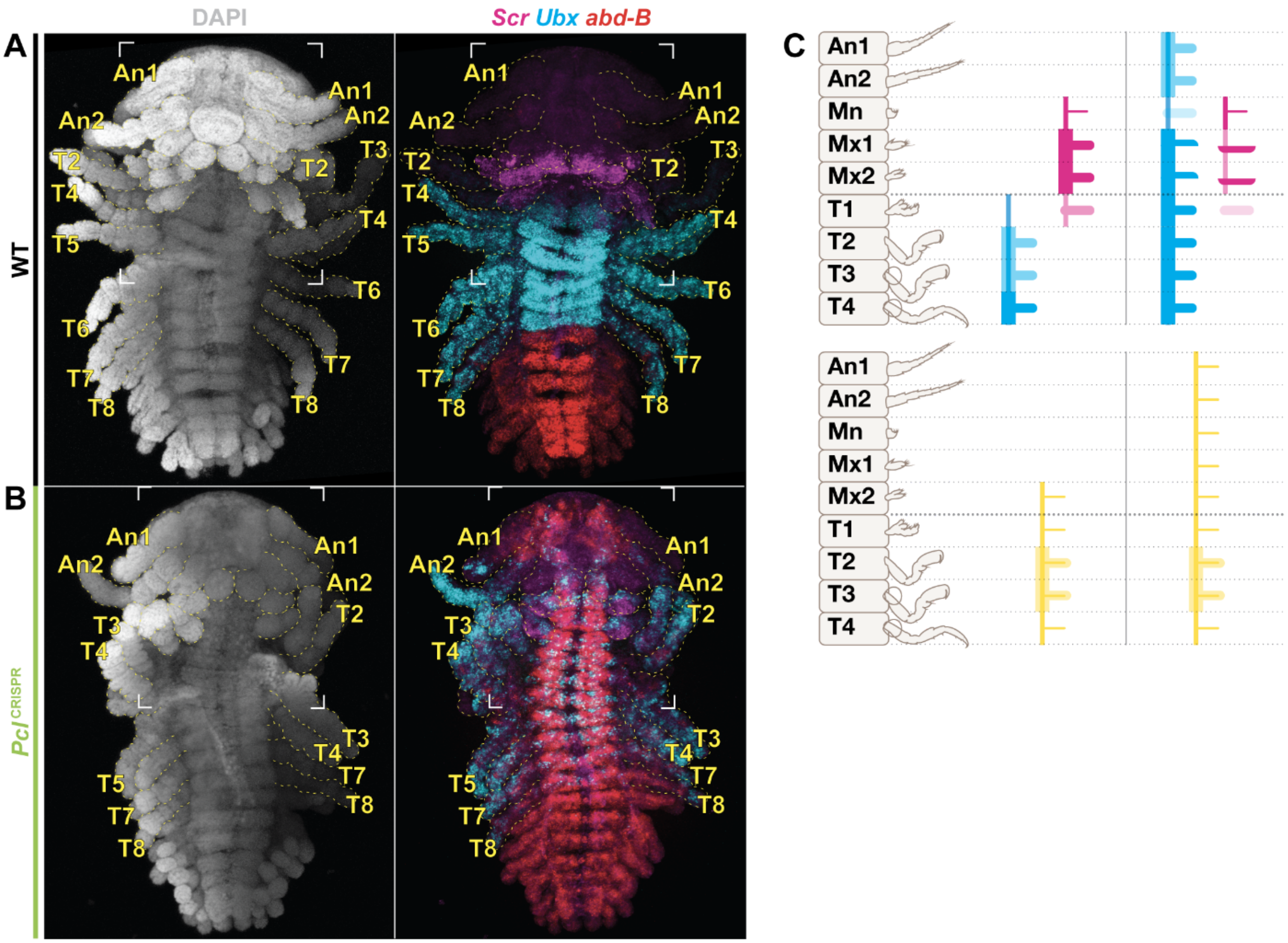


#### Supp. Fig. 5.1: Misexpression of *Scr*, *Ubx*, and *Abd-B* in *Pcl* mutant embryos

A) Wildtype *Parhyale* embryo at S23 showing expression domains of *Scr* (magenta), *Ubx* (cyan), and *Abd-B* (red).

B) *Pcl* CRISPR embryo at S23 showing expression domains of *Scr* (magenta), *Ubx* (cyan), and *Abd-B* (red). The embryo displayed appears to be bilaterally mutant throughout the body axis, as evidenced by strong *Abd-B* expression throughout the head and thoracic nervous system.

C) Schematic of patterns of *Sce*, *Ubx*, and *Antp* expression in wildtype versus PcG mutant embryos.

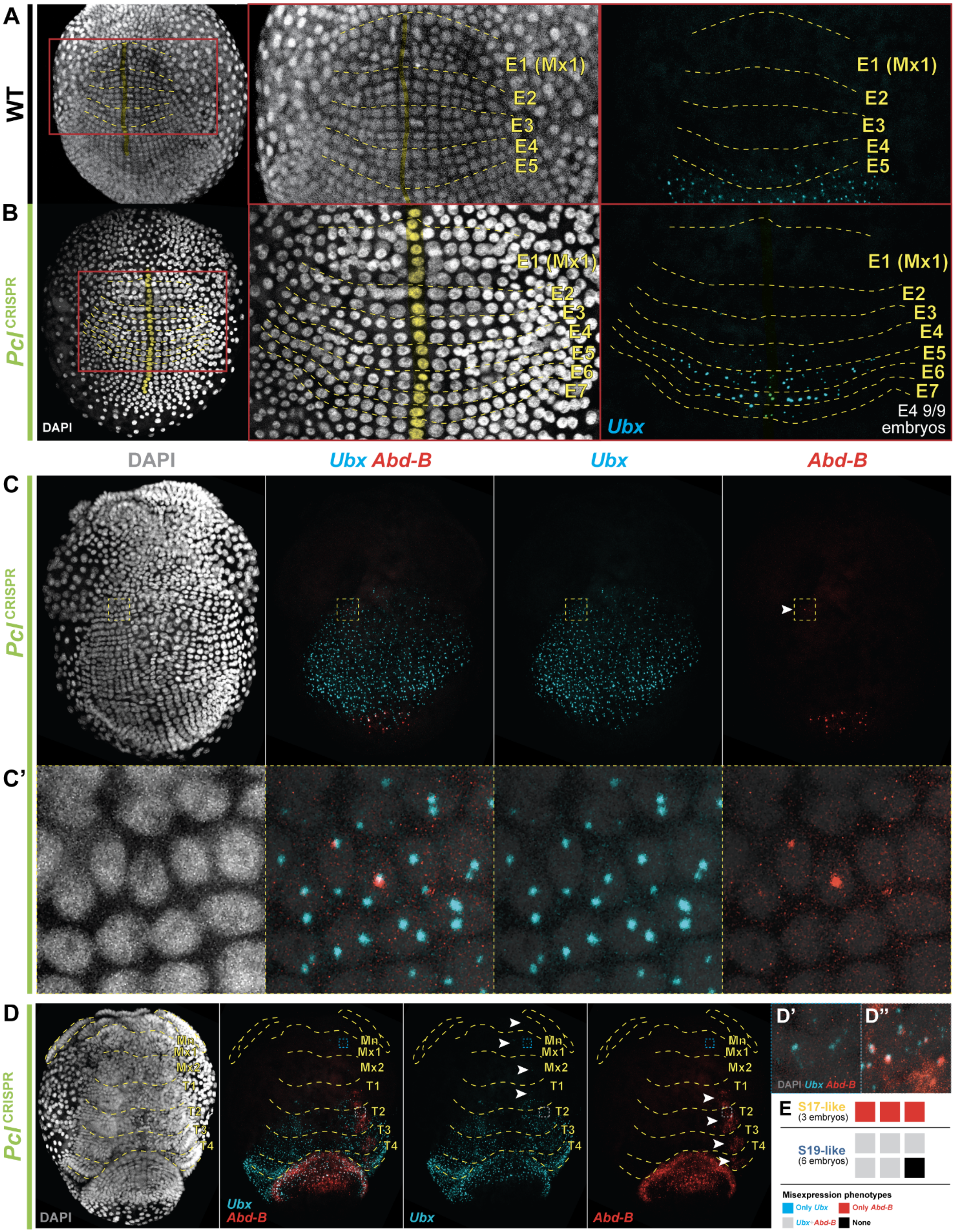

A
WT
E1 (Mx1)
E2
E3
E4
E5
B
Pcl CRISPR
DAPI
E1 (Mx1)
E2
E3
E4
E5
E6
E7
Ubx
E4 9/9 embryos
DAPI
Ubx Abd-B
Ubx
Abd-B
C
C'
D
Mn
Mx1
Mx2
T1
T2
T3
T4
D'
D''
DAPI Ubx Abd-B
E
S17-like
(3 embryos)
S19-like
(6 embryos)
Misexpression phenotypes
Only Ubx
Only Abd-B
Ubx+Abd-B
None

#### Supp. Fig. 6.1: Differences in PcG dependence timing in *Pcl* mutant embryos

A) Wildtype *Parhyale* embryo at S13 showing expression of *Ubx* (cyan) and DAPI. The anterior boundary of *Ubx* expression is the E4 parasegment.

B) *Pcl* CRISPR embryo at S13 showing expression of *Ubx* (cyan). The anterior boundary of *Ubx* expression is the E4 parasegment. We observed the same anterior boundary in 9/9 embryos imaged at this stage.

C) Expression of *Ubx* and *Abd-B* in S17-like *Pcl* mutant embryos. At the same time as *Abd-B* expression is beginning in the posterior, we observed ectopic *Abd-B* expression initiation in the anterior.

C') Inset panel from C) showing a handful of cells with strong *Abd-B* transcriptional spots, and *Ubx* expression in the same cells.

D) Expression of *Ubx* and *Abd-B* in S19-like *Pcl* mutant embryos. We observed misexpression of both *Ubx* and *Abd-B* at this stage, as marked with arrowheads in gene-specific panels; all embryos that showed *Abd-B* misexpression also showed *Ubx* misexpression.

E) Number of embryos showing different patterns of misexpression in S17-like and S19-like embryos dissected concurrently when wildtype embryos were at stage S19. We observed that 3/3 S17-like embryos showed only misexpression of *Abd-B*, while 5/6 S19-like embryos showed simultaneous misexpression of *Ubx* and *Abd-B*.

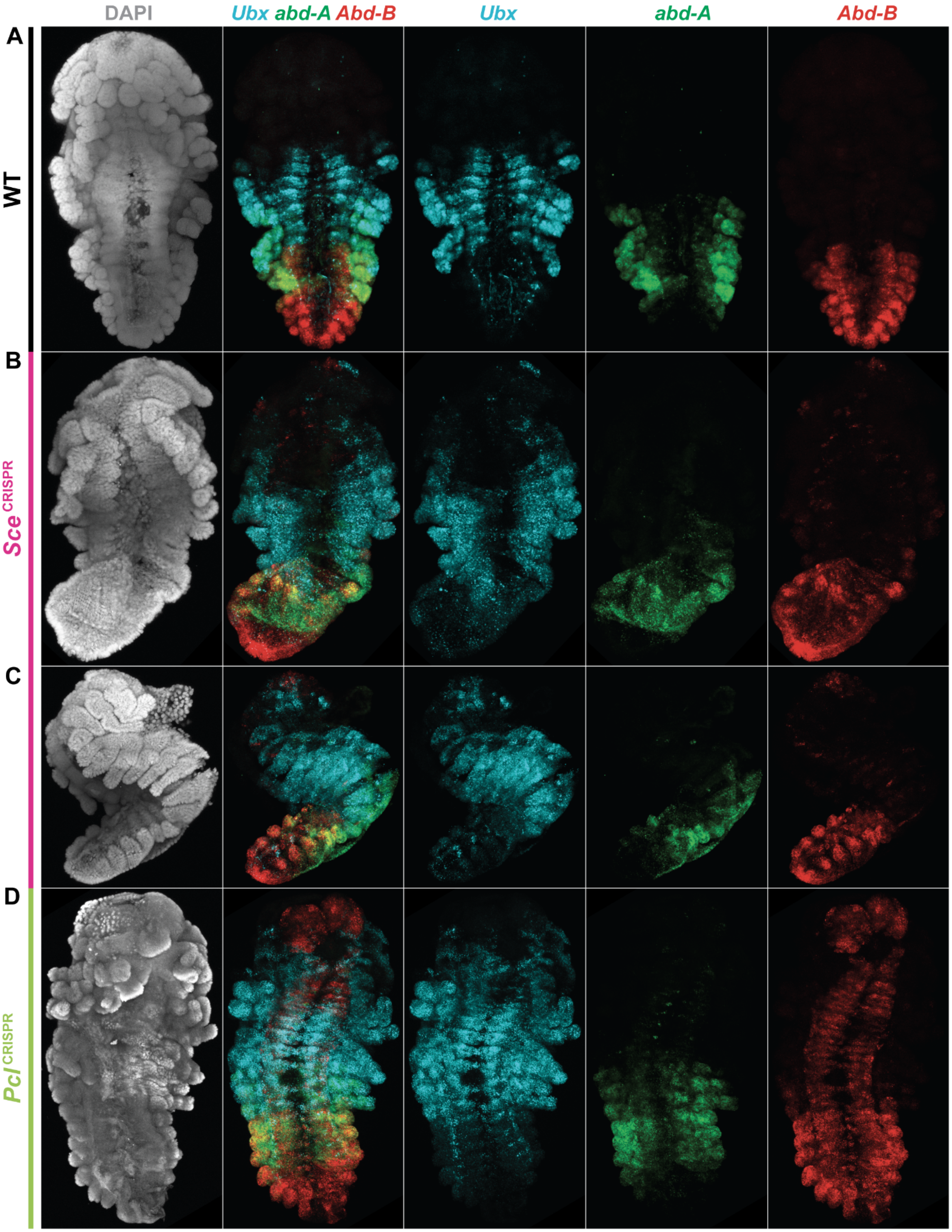

DAPI
Ubx abd-A Abd-B
Ubx
abd-A
Abd-B
A
WT
B
Sce CRISPR
C
D
Pcl CRISPR

#### Supp. Fig. 6.2: Expression of posterior *Hox* genes in S21 embryos

A) Wildtype *Parhyale* embryo at S21 showing expression of *Ubx* (cyan), *abd-A* (green), *Abd-B* (red), and DAPI (gray).

B) *Sce* CRISPR embryo at S21. Weak *Abd-B* and *Ubx* misexpression anterior of the wildtype domain is observed. Very little *abd-A* misexpression is observed.

C) *Sce* CRISPR embryo at S21. This embryo is mounted as a lateral view, and is slightly more morphologically developed than in B. *Ubx* misexpression is much stronger by this point, and small patches of *abd-A* misexpression are beginning.

D) *Pcl* CRISPR embryo at S21. Strong *Ubx* and *Abd-B* misexpression are observed, and the beginning of *abd-A* misexpression is observed.

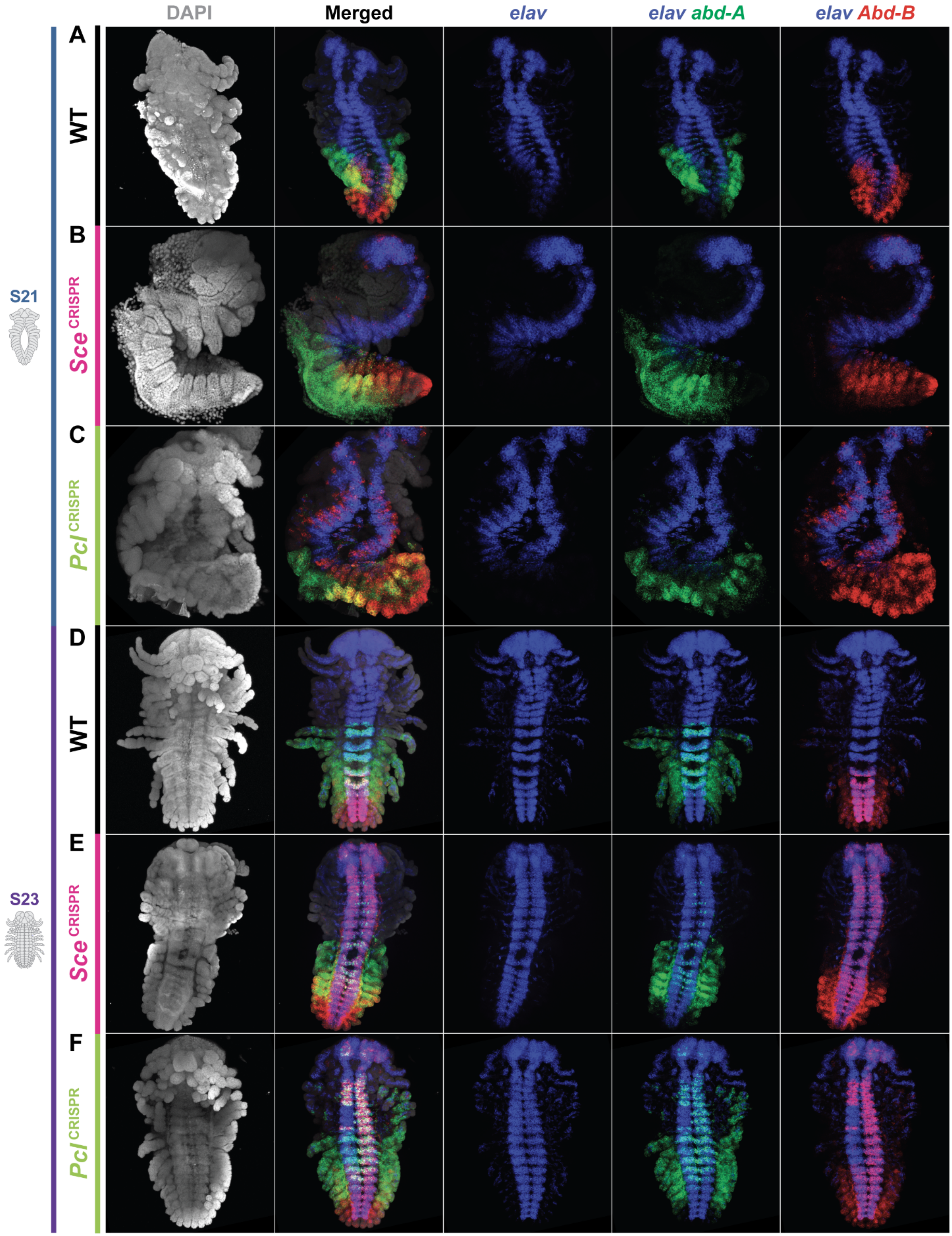

DAPI
Merged
elav
elav abd-A
elav Abd-B
A
WT
B
Sce CRISPR
C
Pcl CRISPR
D
WT
E
Sce CRISPR
F
Pcl CRISPR
S21
S23

#### Supp. Fig. 7.1: *abd-A*, *Abd-B*, and *elav* expression

A) Wildtype *Parhyale* embryo at S21 showing expression of *elav* (blue), *abd-A* (green), *Abd-B* (red), and DAPI (gray).

B) *Sce* CRISPR embryo at S21. Patches of *Abd-B* misexpression are observed at the lateral borders of *elav* expression.

C) *Pcl* CRISPR embryo at S21. Stronger *Abd-B* misexpression is observed than compared to an *Sce* mutant.

D) Wildtype *Parhyale* embryo at S23.

E) *Sce* CRISPR embryo at S23. Strong *Abd-B* misexpression is observed throughout the nervous system. *abd-A* misexpression is restricted to a few cells of the nervous system.

F) *Pcl* CRISPR embryo at S23. Strong *Abd-B* misexpression is observed throughout the nervous system. *abd-A* misexpression expands broadly through the nervous system and into the limbs.

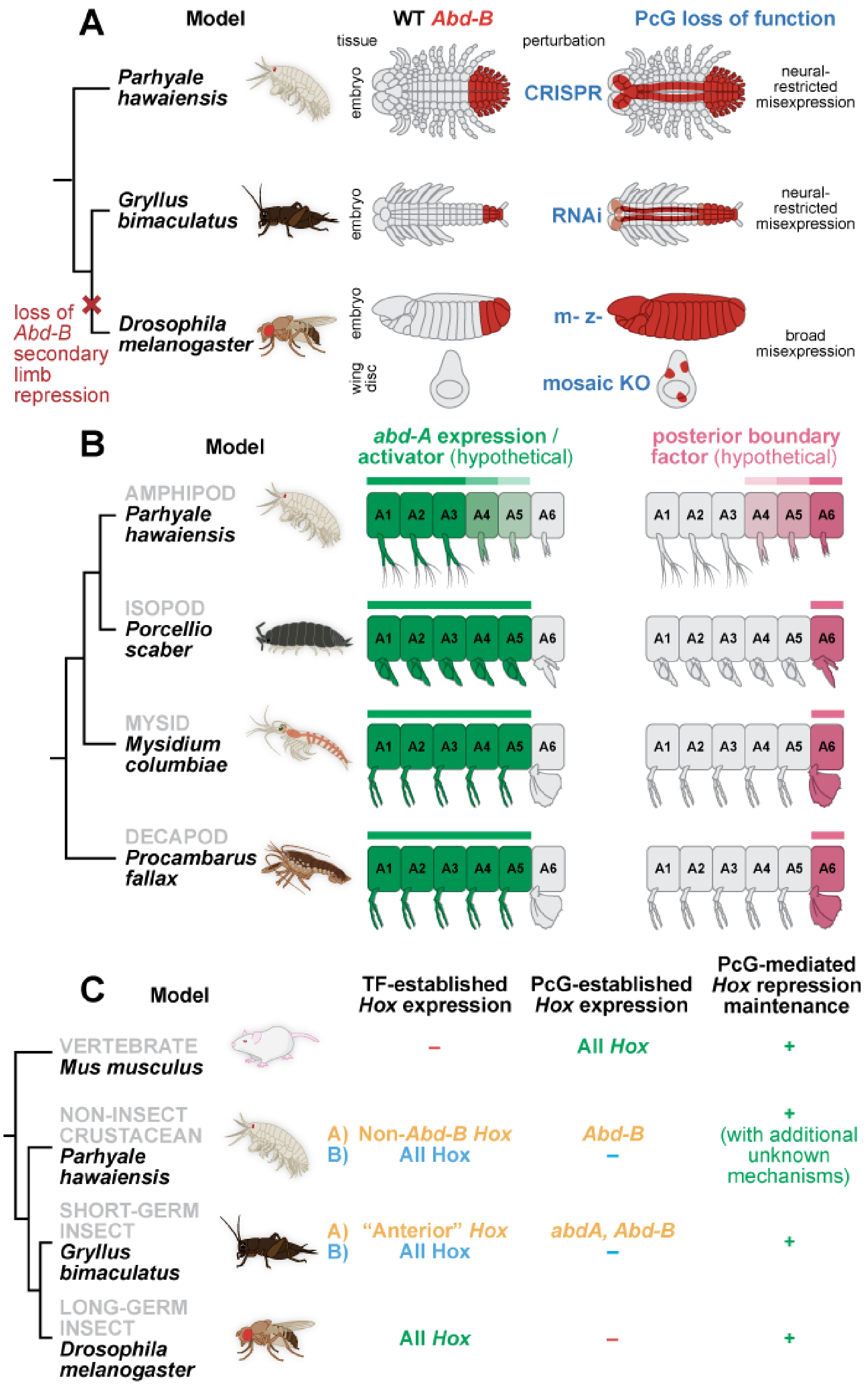
A
Model
WT Abd-B
PcG loss of function
tissue
perturbation
Parhyale hawaiensis
embryo
CRISPR
neural-restricted misexpression
Gryllus bimaculatus
embryo
RNAi
neural-restricted misexpression
loss of Abd-B secondary limb repression
Drosophila melanogaster
embryo
m- z-
broad misexpression
wing disc
mosaic KO
B
Model
abd-A expression / activator (hypothetical)
posterior boundary factor (hypothetical)
AMPHIPOD
Parhyale hawaiensis
ISOPOD
Porcellio scaber
MYSID
Mysidium columbiae
DECAPOD
Procambarus fallax
A1 A2 A3 A4 A5 A6
C
Model
TF-established Hox expression
PcG-established Hox expression
PcG-mediated Hox repression maintenance
VERTEBRATE
Mus musculus
–
All Hox
+
NON-INSECT CRUSTACEAN
Parhyale hawaiensis
A) Non-Abd-B Hox
B) All Hox
Abd-B
–
+ (with additional unknown mechanisms)
SHORT-GERM INSECT
Gryllus bimaculatus
A) "Anterior" Hox
B) All Hox
abdA, Abd-B
–
+
LONG-GERM INSECT
Drosophila melanogaster
All Hox
–
+

#### Supp. Fig. 8.1: Hypothesized *Hox* regulatory mechanisms

A) An unknown limb-specific repression mechanism for *Abd-B* in arthropods. In both *Parhyale* CRISPR knockout of PcG genes and *Tribolium* RNAi, *Abd-B* misexpression extends into the nervous system, but not the limbs. This is in contrast to *Drosophila*, where *Abd-B* misexpression is possible across broad tissue types, including imaginal tissues such as wing discs. This suggests that a potentially ancestral limb-specific repression mechanism of *Abd-B* may have been lost in the lineage leading to flies. Future experiments using PcG knockout in more diverse organisms could reveal whether such a repression mechanism is commonly found across non-dipteran pancrustaceans.

B) A posterior boundary factor for *abd-A*. CRISPR-Cas9 knockout of *Ubx* and *Abd-B*, the *Hox* genes overlapping *abd-A*, do not result in disruption of the pattern of *abd-A* expression (Jarvis et al. 2022). This suggests that *abd-A* is not regulated by the other posterior *Hox* genes. In our study, PcG knockout resulted in misexpression of *abd-A* in the anterior, but no disruption to the pattern of *abd-A* expression in the posterior. This suggests that neither *Hox* cross-regulatory interactions or PcG repression sets the posterior boundary of *abd-A*. We propose a third mechanism, a posterior boundary factor, which represses *abd-A* at the posterior end of the abdomen. We hypothesize that such a factor may also be responsible for the shifts in the *abd-A* expression boundary between different groups of crustaceans.

C) Summary of *Hox* regulatory mechanisms at different phases of development across bilaterians. In *Mus*, *Hox* gene expression establishment is linked to PcG function, whereas in *Drosophila*, PcG repression appears to occur after an early transcription factor-dependent phase of regulation. In examining the *Parhyale* and *Gryllus* data, two possible interpretations emerge: either both *Parhyale* and *Gryllus* have TF-dependent regulation for some *Hox* genes and not others, or both *Parhyale* and *Gryllus* have concurrent TF-dependent and PcG-dependent regulatory mechanisms, which are distinguishable only at very early stages of *Hox* misexpression in PcG mutants. If the latter interpretation is true, this would suggest that arthropods and vertebrates have evolved fundamentally different mechanisms of early *Hox* boundary establishment. Additional experiments with representatives across bilaterians will be needed to determine whether the ancestral bilaterian used PcG- or TF-dependent *Hox* boundary establishment.

### Supp. Table. 1: Select *Drosophila* PcG/TrxG homeotic phenotypes

| Family | *Drosophila* Protein | Genotype | Phenotype(s) | Source(s) |
|---|---|---|---|---|
| **PcG** | Ph | *Ph-d* $^{m-z-}$ *Ph-p* $^{m-z-}$ | A1-7 segments resemble A8 segment. Thoracic segments have denticles characteristic of abdominal segments. | McKeon et al., 1994; Dura et al., 1987; Doe et al., 1988 |
| | Psc | *Psc* $^{m-z-}$ | Anterior segments transformed into A8 segment. *Hox* misexpression. Other early developmental defects. | Martin and Adler, 1993; Morimoto et al., 2016 |
| | Sce | *Sce* $^{m-z-}$ | Anterior→posterior transformations. No segmentation defects. *Ubx* misexpression. | Breen and Duncan, 1986; Fritsch et al., 2003 |
| | Pc | *Pc* $^{m+z-}$ or *Pc* $^{m-z-}$ | Transformation of all thoracic and abdominal segments to A8 segment identity. *En* misexpression. | Chiang et al., 1995; Kassis and Brown, 2013; Haynie, 1983; Lawrence et al., 1983 |
| | Scm | *Scm* $^{m-z-}$ | Anterior→posterior homeotic transformation of most segments. No segmentation defects. | Breen and Duncan, 1986 |
| | Su(z)12 | *Su(z)12* $^{m+z-}$ | All abdominal, thoracic, and some head segments are transformed into the A8 segment. *Abd-B* misexpression in every segment. Antennae partially transformed into legs. Smaller wings. | Birve et al., 2001 |
| | E(z) | *E(z)* $^{m+z-}$ | All thoracic and abdominal segments transformed to more posterior segments. | Phillips and Shearn, 1990 |
| | Pcl | *Pcl* $^{m-z-}$ | A7 segment partially transformed into A8 segment. Dorsal setal belt in head. Wart-like sense organs (normally in A8) in the anterior abdominal segments. Defects in even numbered segments. | Breen and Duncan, 1986 |
| | Esc | *Esc* $^{m-z-}$ | Transformation of all abdominal segments to A8 segment identity. | Wang et al., 2006; Kurzhals et al., 2008; Ohno et al., 2008 |
| **TrxG** | Trx | *Trx* $^{m-z-}$ | Transformation of T1-3 segments into T2 segment identity. Posterior→anterior transformations of abdominal segments. Reduced *Ubx* and *abd-A* protein levels. Haltere to wing transformation (due to decreased *Ubx* expression). Posterior abdominal and genital structures transformed to more anterior identities (due to decreased *abd-A* and *Abd-B* expression). | Gildea et al., 2000; Breen and Harte, 1991; Ingham, 1983; Klymenko and Müller, 2004; Tripoulas et al., 1994; Mazo et al., 1990; Ingham, 1981; Ingham and Whittle, 1980 |
| | Ash2 | *Ash2* $^{m-z-}$ | Various posterior→anterior homeotic transformations including: haltere to wing transformation, leg 3 to leg 2 transformation, and genitalia to leg/antenna transformation. | Shearn, 1989 |
| | Ash1 | *Ash1* $^{m-z-}$ | Various posterior→anterior homeotic transformations including: posterior wing to anterior wing transformation, haltere to wing transformation, leg 3 to leg 2 transformation, and genitalia to leg/antenna transformation. Loss of *En* function. Reduction of *Hox* expression. | Shearn, 1989; Tripoulas et al., 1994; Klymenko and Müller, 2004 |
| | Kis | *Kis* $^{m+z-}$ clones | Homozygous clone of *Kis* $^{m+z-}$ tissue in A5 segment is transformed to a more anterior identity (loss of *Abd-B* function). Some clones showed transformations of first leg towards a second leg identity (loss of *Scr* function). | Daubresse et al., 1999 |
| | | *Kis* $^{m-z-}$ | Pair-rule segmentation defects. | Daubresse et al., 1999 |
| | Mor | *Mor* $^{z-}$ | Embryonic lethal. Head defects that resemble defects seen in *Dfd* $^{z-}$ flies. | Harding et al., 1995 |
| | | *Mor* $^{z-}$ clones | Clones of *Mor* $^{z-}$ cells caused transformations of metanotum and haltere to mesonotum and anterior wing, respectively. Posterior wing to anterior wing transformation (loss of *En* function). | Brizuela and Kennison, 1997 |
| | Brm | *Brm hypomorph* | Reductions in the number of sex comb teeth. Transformations of A5 segment to a more anterior identity. Abnormalities similar to those seen in ANTP-C and BX-C loss-of-function mutants. | Tamkun et al., 1992 |
| | | *Brm* $^{m-}$ | Early embryonic defects. | Brizuela et al., 1994 |
| | Fs1h | *Fs1h* $^{m-z-}$ *temp. sen.* | Homeotic defects: anterior metanotum to anterior mesonotum transformation, and anterior haltere to anterior wing transformation. Mimics loss of *Ubx* function. Mutants often missing legs, halteres, or tergites. | Zalokar et al., 1975 |

### Supp. Table. 2: Select *Mus* PcG/TrxG homeotic phenotypes

| Family | *Drosophila* Protein | Genotype | Phenotype(s) | Source(s) |
|---|---|---|---|---|
| **PcG** | *Ph* | *PHC1-/-* | Perinatal lethality. Anterior→posterior skeletal homeotic transformations. | Isono et al., 2005; Takihara et al., 1997 |
| | *Psc* | *PCGF2-/-* | Anterior→posterior transformation of axial skeleton. Growth retardation. | Akasaka et al., 2001 |
| | | *PCGF4-/-* | Anterior→posterior transformation of axial skeleton. | Akasaka et al., 2001 |
| | | *PCGF2-/- PCGF4-/-* | Embryonic lethal. | Akasaka et al., 2001 |
| | *Sce* | *RING1A+/-* or *-/-* | Anterior→posterior transformations of vertebrae. Rib abnormalities. | del Mar Lorente et al., 2000 |
| | | *RING1B-/-* | Anterior→posterior homeotic transformations of the axial skeleton. | Suzuki et al., 2002 |
| | *Pc* | *CDX2-/-* | Malformations of axial skeleton. Reduced viability. Poor growth. Male to female sex reversal. T7 vertebra transformed into T8 vertebra. | Coré et al., 1997 |
| | | *CDX4-/-* | Complete neonatal lethality. Severe thymus hypoplasia. | Coré et al., 1997 |
| | | *CDX7-/-* | Increased body length. Developed liver and lung adenomas and carcinomas. | Forzati et al., 2012 |
| | *Scm* | *Scmh1* $^{SPM-/-}$ | Anterior→posterior vertebral transformations. Male infertility. | Takada et al., 2007 |
| | | *Scmh1-/-* | Abnormal hematopoiesis. Normal fertility and skeleton. | Yasunaga et al., 2013 |
| | *Su(z)12* | *Su(z)12-/-* | Embryonic lethal. | Li et al., 2011 |
| | | *Su(z)12+/-* | Anterior tuberculums (characteristic of C6 vertebra) associated with C5 vertebra. Anteriorly shifted sternum. Spinous process (characteristic of T2 vertebra) associated with T1 vertebra. | Li et al., 2011 |
| | *E(z)* | *EZH2 -/-* | Embryonic lethal. | O'Carroll et al., 2001 |
| | | *EZH2 cond. loss* | Severe growth retardation of neonate. Gastrulation failure. | O'Carroll et al., 2001 |
| | *Pcl* | *Pcl2 -/-* | Anterior→posterior transformation of axial skeleton. | Wang et al., 2007 |
| | *Esc* | *EED -/-* | Early gastrulation defect. | Wang et al., 2002 |
| | | *EED hypomorph* | Anterior expression boundaries of several *Hox* genes were shifted rostrally by one segment. | Wang et al., 2002 |
| **TrxG** | *Trx* | *Mll -/-* | *Hoxa7* expression is activated normally but is not maintained. Branchial arch dysplasia. Aberrant segmental boundaries of spinal ganglia and somites. | Yu et al., 1998 |
| | *Ash2* | *Ash2l -/-* | Die early during gestation. | Stoller et al., 2010 |
| | *Ash1* | *Ash1l -/- (gene trap)* | Corpus to caput transformation. Widening of the cauda. Transformations similar to posterior→anterior transformations that occur in *Hoxa10-/-* mutant males. | Brinkmeier et al., 2015 |
| | *Kis* | *CHD7 -/-* | Embryonic lethal. | Sperry et al., 2014 |
| | *Brm* | *Brm -/-* | Increased cell proliferation. | Reyes et al., 1998 |
| | *Fs1h* | *BRD2 -/-* | Embryonic lethal. | Shang et al., 2009 |

#### Supp. Table. 3: *Parhyale* PcG genes

| Complex | *Parhyale* Gene Name | *Drosophila* Protein | *Mus* Proteins | Best *Parhyale* Transcript | Txome | *Parhyale* IGV Address |
|---|---|---|---|---|---|---|
| **PRC1** | Par-haw_ph | ph-d [Q9W523], ph-p [P39769] | PHC1 [Q64028], PHC2 [Q9QWH1], PHC3 [Q8CHP6] | mikado.phaw_50.283863eG7.1 | Sun-Mikado | phaw_50.283863e:707,281-945,757 |
| | Par-haw_Psc-1 | Psc [P35820] | PCGF1 [Q8R023] | mikado.phaw_50.283028bG288.1 | Sun-Mikado | phaw_50.283028b:22654336-22754322 |
| | Par-haw_Psc-2 | Su(z)2 [P25172] | PCGF2 [P23798] | mikado.phaw_50.283028bG712.1 | Sun-Mikado | phaw_50.283028b:21,822,602-21,858,763 |
| | | | PCGF3 [Q8BTQ0], PCGF4 [P25916], PCGF5 [Q3UK78], PCGF6 [Q99NA9] | | | |
| | Par-haw_Sce | Sce [Q9VB08] | RING1A [O35730], RING1B [Q9CQJ4] | mikado.phaw_50.283875aG27.1 | Sun-Mikado | phaw_50.283875a:2670126-2804666 |
| | Par-haw_Pc | Pc [P26017] | CBX2 [P30658], CBX4 [O55187], CBX6 [Q9DBY5], CBX7 [Q8VDS3], CBX8 [Q9QXV1] | mikado.phaw_50.000214fG102.1 | Sun-Mikado | phaw_50.000214f:7,059,547-7,067,561 |
| **dRAF** | includes Par-haw_Psc-1*, Par-haw_Psc-2*, Par-haw_Sce* (see above) | | | | | |
| | Par-haw_Kdm2-isoform_a | Kdm2 [Q9VHH9] | Kdm2a [P59997] | STRG.39002.1 | Sun-StringTie2 | - |
| | Par-haw_Kdm2-isoform_b | | | mikado.phaw_50.283869cG487.1 | Sun-Mikado | phaw_50.283869c:39,738,428-39,800,237 |
| **PR-DUB** | Par-haw_Calypso | Calypso [Q7K5N4] | Bap1 [Q99PU7] | comp166625_c1_seq3 | Trinity-Limb | phaw_50.000214e:2,663,327-2,803,232 |
| | Par-haw_Asx | Asx [Q9V727] | ASXL1 [P59598], ASXL2 [Q8BZ32], ASXL3 [Q8C4A5] | mikado.phaw_50.283821G75.2 | Sun-Mikado | phaw_50.283821:4,766,576-4,819,515 |
| **Scm** | Par-haw_Scm | Scm [Q9VHA0] | SCMH1 [Q8K214], SCML2 [B1AVB5], SCML4 [Q80VG1] | comp171618_c0 | Trinity-Limb | phaw_50.007301a:14,931,316-14,976,242 |
| **PRC2.1** | Par-haw_Su(z)12 | Su(z)12 [Q9NJG9] | SUZ12 [Q80U70] | mikado.phaw_50.283867cG387.1 | Sun-Mikado | phaw_50.283867c:24503623-24641721 |
| | Par-haw_E(z) | E(z) [P42124] | EZH1 [P70351], EZH2 [Q61188] | mikado.phaw_50.001137bG316.1 | Sun-Mikado | phaw_50.001137b:17254399-17416111 |
| | Par-haw_Caf1 | Caf1 [Q24572] | Rbbp4 [Q60972] | mikado.phaw_50.283826G445.1 | Sun-Mikado | phaw_50.283826:29293007-29339138 |
| | Par-haw_Pcl | Pcl [Q24459] | PCL2 [Q02395], PCL3 [Q9CXG9] | mikado.phaw_50.283817cG134.1 | Kao-Mikado | phaw_50.283817c:3067340-3096584 |
| | Par-haw_HDAC1 | Rpd3 [Q94517] | Hdac1 [O09106] | comp160143_c0_seq1 | Trinity-Limb | - |
| | Par-haw_Esc | Esc [Q24338] | EED [Q921E6] | mikado.phaw_50.283863aG40.1 | Sun-Mikado | phaw_50.283863a:3,188,185-3,235,909 |
| **PRC2.2** | includes Par-haw_Su(z)12*, Par-haw_E(z)*, Par-haw_Caf1*, Par-haw_Esc* (see above) | | | | | |
| | Par-haw_Jarid2 | Jarid2 [Q9VT00] | Jarid2 [Q62315] | mikado.phaw_50.283865bG254.1 | Kao-Mikado | - |
| | Par-haw_Jing | Jing [Q7KHG2] | Aebp2 [Q9Z248] | mikado.phaw_50.282260aG393.1 | Sun-Mikado | - |
| **PhoRC** | Par-haw_pho | Pho [Q8ST83] | YY1 [Q00899] | mikado.phaw_50.000289aG44.1 | Sun-Mikado | phaw_50.000289a:1971807-1992607 |
| | Par-haw_Sfmbt | dSfmbt [Q9VK33] | SFMBT1 [Q9JMD1], SFMBT2 [Q5DTW2] | mikado.phaw_50.283867cG203.1 | Sun-Mikado | phaw_50.283867c:14916319-14971921 |

#### Supp. Table. 4: *Parhyale* TrxG genes

| Complex | *Parhyale* Protein Name | *Drosophila* Protein | *Mus* Protein | Best *Parhyale* Transcript | Txome | *Parhyale* IGV Address |
|---|---|---|---|---|---|---|
| dCOMPASS-like | Par-haw_Trx (SET) | Trx [P20659] | KMT2B [O08550] | mikado.phaw_50.282275aG400.1 | Sun-Mikado | phaw_50.282275a:22,898,633-23,055,618 |
| | Par-haw_Trx (DBD) | | | mikado.phaw_50.282275aG397.1 | Sun-Mikado | - |
| | Par-haw_Ash2 | Ash2 [Q94545] | Ash2l [Q91X20] | comp174477_c0_seq5 | Trinity-Limb | phaw_50.282695g:2178005-2220752 |
| dCOMPASS | includes Par-haw_Ash2 (see above) | | | | | |
| | Par-haw_Set1 | Set1 [Q5LJZ2] | Setd1a [E9PYH6] | mikado.phaw_50.283355aG178.1 | Sun-Mikado | phaw_50.283355a:9493556-9551696 |
| Trr | Par-haw_Trr | Trr [Q8IRW8] | Kmt2c [Q8BRH4] | mikado.phaw_50.282861eG448.1 | Kao-Mikado | phaw_50.282861e:9,250,785-9,312,955 |
| TAC1 | includes Par-haw_Trx (see above) | | | | | |
| | Par-haw_nej | nej [M9MS40] | Crebbp [P45481] | mikado.phaw_50.282695dG124.1 | Sun-Mikado | phaw_50.282695d:7790068-7814604 |
| | Par-haw_Sbf1 | Sbf [Q9VGH9] | SBF1 [Q6ZPE2], SBF2 [E9PXF8] | mikado.phaw_50.283826G523.1 | Sun-Mikado | phaw_50.283826:34087962-34200876 |
| Ash1 | Psr-haw_Ash1 | Ash1 [Q9VW15] | Ash1l [Q99MY8] | mikado.phaw_50.007301aG268.1 | Kao-Mikado | phaw_50.007301a:3,925,168-4,034,186 |
| Kis | Par-haw_Kis | Kismet [Q9Y1L3] | Chd6 [A3KFM7], Chd7 [A2AJK6] | mikado.phaw_50.283870G403.1 | Sun-Mikado | phaw_50.283870:21945760-22131964 |
| PBAP | Par-haw_Osa | Osa [Q8IN94] | Arid1a [A2BH40], Arid1b [E9Q4N7] | mikado.phaw_50.283815cG377.1 | Sun-Mikado | phaw_50.283815c:19852707-19939426 |
| | Par-haw_Mor | Mor [Q7KPY3] | Smarcc2 [Q6PDG5] | mikado.phaw_50.283823aG160.1 | Sun-Mikado | phaw_50.283823a:6311700-6337044 |
| | Par-haw_Snr1 | Snr1 [Q24090] | Smarcb1 [Q9Z0H3] | mikado.phaw_50.282275bG198.1 | Kao-Mikado | phaw_50.282275b:3482023-3497731 |
| | Par-haw_Brm | Brm [P25439] | Smarca2 [Q6DIC0], Smarca4 [Q3TKT4] | mikado.phaw_50.283873aG49.1 | Sun-Mikado | phaw_50.283873a:2837385-2858196 |
| BAP | includes Par-haw_Mor*, Par-haw_Snr1*, Par-haw_Brm* (see above) | | | | | |
| | Par-haw_SAYP | SAYP [Q9VWF2] | Phf10 [Q8WUB8] | mikado.phaw_50.000135aG190.1 | Kao-Mikado | phaw_50.000135a:3600922-3684882 |
| Other | Par-haw_Fs1h | Fs1h [P13709] | - | comp_163013 | Trinity-Limb | phaw_50.283821:4,406,782-4,518,976 |
| | Par-haw_Kto | Kto [Q9VW47] | Med12l [Q8BQM9] | comp_154627 | Trinity-Limb | phaw_50.283867c:20,195,413-20,284,996 |
| | Par-haw_Skd | Skd [Q7KTX8] | Med13 [Q5SWW4] | comp_160166 | Trinity-Limb | phaw_50.283867c:20,195,413-20,284,996 |

#### Supp. Table. 5: CRISPR guide sequences

| Family | Gene | CRISPR Guide ID | CRISPR Guide Sequence |
|---|---|---|---|
| PcG | Par-haw_Psc-1 | Syn078_Psc-1_g1 | AAAGCACACACACGAAGTGG |
| | | Syn079_Psc-1_g2 | AGCAACAGAGTGTCGCAGTG |
| | Par-haw_Psc-2 | Syn039_PhawPsc2_g1 | GTGAGAGGAATTCGGCGTAG |
| | | Syn040_PhawPsc2_g2 | CACAGATGAGGTGTGCGTTT |
| | Par-haw_Sce | Syn041_PhawSce_g1 | AACAGCAGAACTACTGTTGT |
| | | Syn042_PhawSce_g2 | TGAACTGCATCGAACTCCTC |
| | Par-haw_Pc | Syn059_Pc_g1 | GAATACCTTGTAAAATGGAA |
| | | Syn060_Pc_g2 | TTTAGATGGACGCCTGCTTG |
| | Par-haw_E(z) | Syn084_Ez_g1 | ATCATCAATGAAGCCCGTCT |
| | | Syn085_Ez_g2 | GCCTCGGGCAACAAGGCCAG |
| | Par-haw_Pcl | Syn051_PhawPcl_g1 | CTCGGGCGCACTTGCAAGCG |
| | | Syn052_PhawPcl_g2 | CAGCCAACACAAGAGCAGCT |
| | Par-haw_Esc | Syn082_Esc_g1 | TTGAATGCCACACCAAACAG |
| | | Syn083_Esc_g2 | GTGTACCAGTGCGAGAAGCA |
| | Par-haw_pho | Syn086_pho_g1 | CGACTATGCAGAGTACATGG |
| | | Syn087_pho_g2 | GGGTCACTGAGATCAACATC |
| TrxG | Par-haw_Trx | Syn043_PhawTrx_g1 | GAGGATCCATGTGGTAATGG |
| | | Syn044_PhawTrx_g2 | GGGATCCGGACAGAGGAGGA |
| | | Syn057_PhawTrx_rg1 | TGTCAGCGACTGGTTGCCAT |
| | | Syn058_PhawTrx_rg2 | GACCAAGTGGTGAAGTCACT |
| | | Syn061_Trx_rd3g1 | TCGTAACACCCCTGACACAG |
| | | Syn062_Trx_rd3g2 | AGTGCGGTCAGTGCAAGGAA |
| | | Syn076_Trx-SET_g1 | GTCCGAGGATGGCTTCAAAG |
| | | Syn077_Trx-SET_g2 | GTGGCAGACGGTCTTTGATG |
| | Par-haw_Ash2 | Syn080_Ash2_g1 | GCCCCTTGAGACACCATGAG |
| | | Syn081_Ash2_g2 | AGCCTGATGGGTCACATGTA |
| | Par-haw_Kis | Syn055_PhawKis_g1 | GGCAGACGTCTCGAGCACTG |
| | | Syn056_PhawKis_g2 | CTCAGACACCCAAAAAGAAG |
| | | Syn063_Kis_rd2g1 | ACGGAAGAGGAATTACTCAA |
| | | Syn064_Kis_rd2g2 | AACAGCAACCTTCTTTCCTG |
| | Par-haw_Brm | Syn053_PhawBRM_g1 | TGGCCAGAGCCCAAGCCCTG |
| | | Syn054_PhawBRM_g2 | GGAGGAGGAGGCATTGGTGA |
| | Par-haw_Fs1h | Syn069_fs1h_g1 | AAGCTGCCTGGCGATAAGCT |
| | | Syn070_fs1h_g2 | GGATTGGAATCTCTGAGAGA |
| | Par-haw_Kto | Syn065_kto_g1 | TCCTGCTAAATCTTTGAACC |
| | | Syn066_kto_g2 | AACCAGATGGCACGCTGCGT |
| | Par-haw_Skd | Syn067_skd_g1 | CTGCCACAAGACTTGGACTG |
| | | Syn068_skd_g2 | CAACTGCAGTATTTCCACCT |
